# A solid-state heater-imager for quantitative evaluation of colorimetric isothermal nucleic acid amplification on paper

**DOI:** 10.64898/2026.03.03.709423

**Authors:** Bibek Raut, Gopal Palla, Dan Van Nguyen, Rachel A Munds, Abdullah Bayram, Virendra Kumar, Bilal Ahmed, Cindy Mayorga, Aaron Ault, Aaron Gilbertie, J. Alex Pasternak, Mohit S. Verma

## Abstract

Maintaining precise isothermal conditions in portable nucleic acid amplification tests (NAATs) is critical for reproducible results but remains challenging with conventional single-sided thin-film heaters, which exhibit temperature gradients and strong dependence on ambient conditions. To close this gap, we engineered ThermiQuant™ VitroMini, a dual-sided heater design that achieves volumetric-level temperature uniformity using thin-film heaters while preserving optical transparency for real-time colorimetric loop-mediated isothermal amplification (LAMP) analysis on microfluidic paper-based analytical devices (µPADs). The device integrates two independently regulated indium tin oxide (ITO) heaters (8 Ω each) controlled by independent proportional–integral–derivative (PID) algorithms. Heaters were evaluated under controlled ambient environments of 4 °C (refrigerated), 23 °C (room temperature), and 50 °C (oven). Analytical tests were performed using a colorimetric LAMP assay targeting the SARS-CoV-2 *orf7ab* gene on µPADs preloaded with dried LAMP reagents, with time-lapse images (30 seconds interval) analyzed via Amplimetrics™ software. VitroMini maintained 65 ± 0.5 °C across 4 to 50 °C ambient conditions and achieved a limit of detection of 34 copies/reaction (4.5 copies/µL) and limit of quantification of 1000 copies/reaction (133 copies/µL), with quantification time (Tq) linearly correlated with log_10_ DNA concentration. Dual-sided heating eliminated temperature bias, condensation artifacts, and ambient-dependent variability while preserving optical transparency for real-time quantitative LAMP reaction. ThermiQuant™ VitroMini bridges the gap between benchtop volumetric heaters and portable diagnostic devices, offering a compact, low-power platform for quantitative colorimetric molecular analysis on paper with potential for decentralized and field-deployable applications.

## Introduction

Single-sided thin-film heaters are widely used in portable isothermal nucleic acid amplification tests (NAATs), but they must operate above the target reaction temperature to compensate for heat loss to the environment^1–5^. The magnitude of this offset depends on ambient conditions, and the resulting temperature gradients orthogonal to the reaction surface are often sufficient to induce localized condensation, reducing assay reproducibility across different ambient temperatures. To address this limitation, we developed a dual-sided indium tin oxide (ITO) thin-film heater design that eliminates the need for temperature offsets, maintains ±0.5 °C temperature uniformity at the 65 °C setpoint across 4 to 50 °C ambient conditions, and provides a transparent optical window for real-time (every 30 s) colorimetric LAMP monitoring.

Isothermal nucleic acid amplification tests (NAATs) such as loop-mediated isothermal amplification (LAMP) have gained broad attention as rapid, minimal-equipment alternatives to polymerase chain reaction (PCR) for molecular diagnostics in low-resource or field settings^6–12^. The integration of LAMP into microfluidic paper-based analytical devices (µPADs) enables low-cost, disposable assay formats that can be easily adapted for one-pot sample preprocessing, amplification, and detection^8,9^. LAMP operates at a constant temperature (typically 60–65 °C) and, when coupled with colorimetric reporters, produces visible color shifts that in principle allow naked-eye interpretation^10,13–16^. However, at low analyte concentrations near the limit of detection (LOD), the color change can be spatially heterogeneous, leading to ambiguous visual reads^17–20^. Therefore, a minimal instrument capable of maintaining precise isothermal conditions while continuously tracking color change remains essential for reliable PON diagnostics using colorimetric µPAD-LAMP assays.

Several research groups (as summarized in Table S1) have explored a wide range of heater designs for isothermal NAATs, which can be broadly divided into two categories: volumetric and thin-film heating. Volumetric heaters, such as circulating water baths or metal heat blocks, deliver heat from all sides of the reaction chamber, enabling rapid heat transfer and excellent temperature uniformity (typically ± 0.1–0.5 °C at 60–65 °C). Transparent water baths additionally allow full field-of-view optical imaging through the medium. Leveraging this capability, we previously developed several water bath–based platforms, including Field Applicable Rapid Microbial (FARM)-LAMP^16^, IsoHeat^21^, and ThermiQuant™ variants such as MegaScan^20^ and AquaStream^22^ (the latter two currently reported as prepri**n**ts), which demonstrated uniform temperature distribution (± 0.**5** °C) and robust quantitative evaluation of µPAD-LAMP assays through time-lapse image analysis. However, these systems rely on water baths and are therefore bulky (often >5 kg, benchtop format) and require routine maintenance. Other groups have employed metal heat blocks, primarily for liquid assays and utilized tiny op**t**ical apertures (usually < 2 mm diameter) for optical sensing during incubation^23^. While effective for uniform heating, **s**uch configurations provide limited optical access, which is insufficient for µPAD-based assays that exhibit spatially heterogeneous color development and require full-field imaging.

To enable portable and compact heating, most recent colorimetric LAMP instruments have adopted thin-film resistive heaters, often integrated directly onto printed circuit boards (PCBs) or fabricated as metal-foil or conductive-film layers^3,24–26^. This design simplifies device construction, reduces power consumption, and allows imaging or sensing from the non-opaque side of the heater, providing a clear optical window for real-time colorimetric readout. However, because heating occurs from only one surface while the other surface is at ambient temperature, a thermal gradient develops orthogonal to the heater plate, requiring the heater to operate at a higher setpoint to maintain the desired reaction temperature. For example, in Ubiquitous NAAT (UbiNAAT), a heater setpoint of 73 °C was required to maintain a 63 °C reaction temperature^3^.

This through-thickness gradient also creates asymmetric water vapor conditions within the cartridge. Specifically, the µPAD surface in contact with the heater is warm, while the opposing surface, exposed to ambient air, remains relatively cooler. When small air gaps are present (>0.3 mm in our case) between the cartridge ceiling and the µPAD, water vapor generated during heating becomes trapped in this confined space and condense on the cooler surface. In our experience, even small mechanical gaps (0.3 mm) between the cartridge ceiling and the µPAD surface were enough to induce condensation forming tiny droplets at the cooler surface, affecting both LAMP kinetics and image clarity (presented in detail in the results section). Although this phenomenon has not been widely discussed in literature, it represents a practical challenge we have frequently encountered during device operation and must be carefully managed to ensure consistent µPAD-LAMP assay performance. Moreover, we have noticed that at high concentrations of the target, the evaporation and condensation do not alter assay performance noticeably; however, when the target concentration is close to the LOD, the assay becomes unreliable and thus the effective LoD becomes worse than what might be possible if a uniform heating system was used.

Please do not adjust margins

Here, we present ThermiQuant™ VitroMini (Figure 1), an indium tin oxide (ITO)–based thin-film heater system that advances conventional single-sided designs through a dual-sided architecture, achieving volumetric-like thermal precision (65 ± 0.5 °C) with minimal sensitivity to ambient conditions. Compared to volumetric water-bath systems, which provide excellent thermal uniformity but are bulky, require liquid handling, and are less suited for portable use, VitroMini achieves comparable temperature precision (65 ± 0.5 °C) in a compact, low-power, and maintenance-free format. Compared to metal block heaters, which typically offer limited optical access, the VitroMini preserves full-field optical transparency for real-time imaging of µPAD-based assays. Furthermore, compared to conventional single-sided thin-film heaters, which suffer from temperature gradients and ambient sensitivity, the dual-sided design enables volumetric-like thermal performance with minimal dependence on environmental conditions. In controlled tests across 4–50 °C environments, ThermiQuant™ VitroMini maintained stable performance, with only a 1.2-min difference in ramp-up time to reach 65 °C between the coldest (4 °C) and hottest (50 °C) conditions. When tested with a colorimetric LAMP assay targeting the SARS-CoV-2 *orf7ab* g**e**ne, VitroMini achieved a LOD95 of 34 copies/reaction (4.5 copies/µL), consistent with prior volumetric systems^16,20–22^, and demonstrated a limit of quantification (LOQ) of 1,000 copies/reaction (133 copies/µL), enabling quantitative analysis via log-linear calibration of quantification time (Tq) against log_10_ DNA concentration. Thus, ThermiQuant™ VitroMini, combining precise thermal regulation (“Therm”) and integrated imaging (“i”) for quantitative (“Quant”) colorimetric analysis on a glass-based (“Vitro”) and compact (“Mini”) platform, offers precision, portability, and environm**e**ntal resilience with potential for applications in clinical diagnostic**s**, as well as agricultural, veterinary, and environmental testing.

**Fig. 1.**
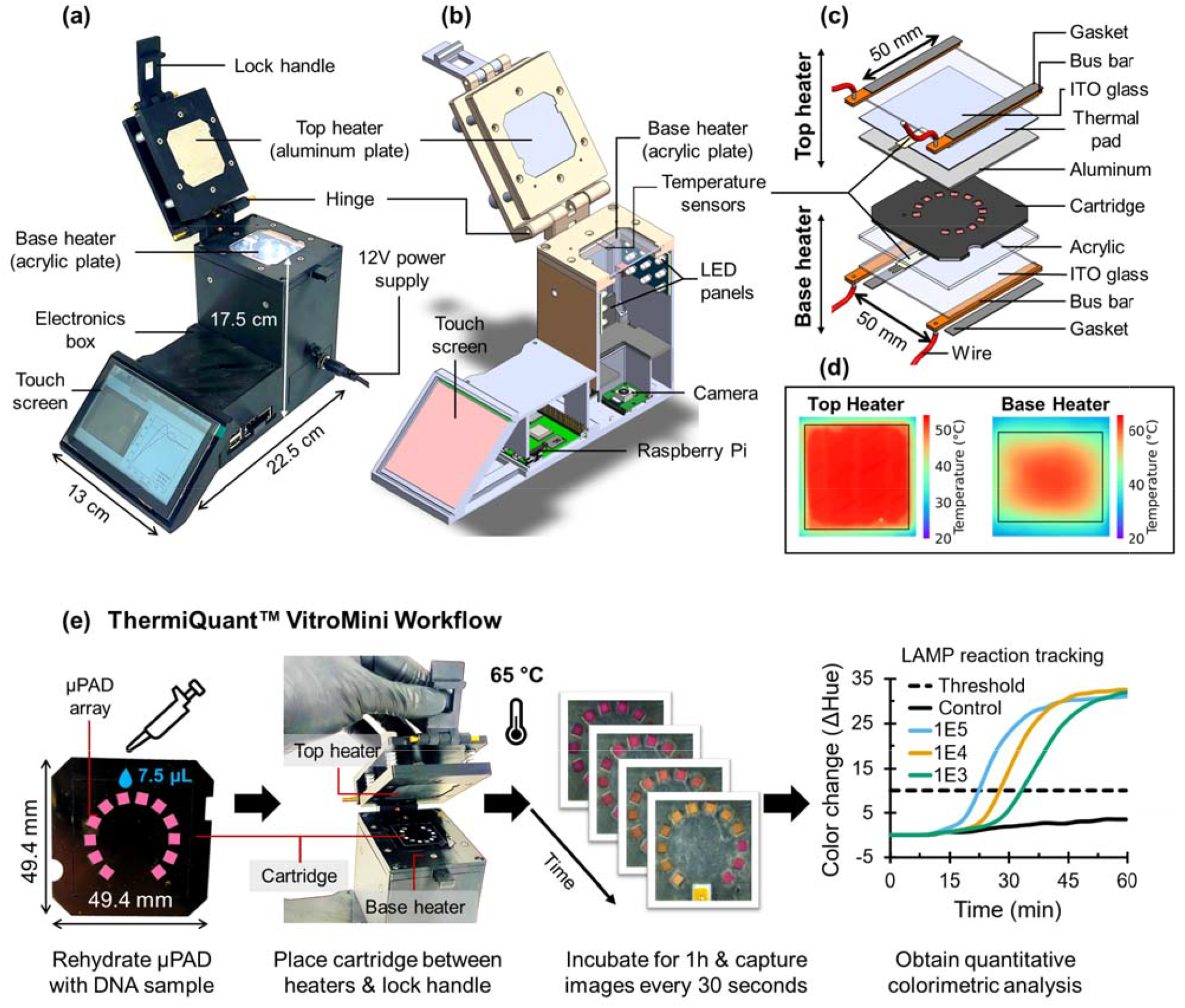
Design and workflow of the ThermiQuant™ VitroMini for quantitative colorimetric LAMP analysis on microfluidic paper-based analytical devices (µPADs). (A) Photograph of the assembled instrument. (B) Sectional side view showing internal components, including the camera, dual heater units, and LED illumination panel. (C) Exploded schematic of the dual-sided heater assembly with top and base indium tin oxide (ITO)–coated glass layers and a 1.5-mm thick cartridge containing 12 µPADs sandwiched between them. Each heater includes a thermistor positioned 13 mm from the center for independent temperature feedback control. (D) Thermal maps of the top and base heaters after 5 min of heating at 12 V (8 Ω, 18 W each), obtained by heating each heater separately. The top heater was mounted on an aluminium plate to promote uniform heat distribution, while the base heater was attached to an acrylic plate to preserve optical transparency for image capture. (E) Workflow for µPAD-LAMP experiments using VitroMini. Pre-dried µPADs are rehydrated with sample and sealed with transparent film, then sandwiched between the two heaters. The reaction is incubated at 65 °C for 1 h with imaging every 30 s, and the resulting time-lapse data are converted into hue-versus-time amplification curves for quantitative analysis.

## Methods

### ThermiQuant™ VitroMini design and assembly

The mechanical parts of ThermiQuant™ VitroMini instrument (Figure 1 and 2) were designed in SolidWorks (Dassault Systèmes SolidWorks Corp., USA) and 3D printed using a Bambu Lab X1 Carbon 3D printer (Bambu Lab, China) with black polycarbonate filament. The custom PCB boards were fabricated by JLCPCB (China) and designed using their EasyEDA software and manually assembled in house. Design files and software are provided in SI and bill of material (BOM) in Table S2. Instrument assembly guide is shown in Figure S1 and Figure S2 and described in Note 1.1. Instrument and software workflow and user guide is shown in “Video1_Instrument_Demo_SD.mp4” in SI. Note 1.2 in the SI contains the full video transcript, and Figure S3 presents key video screenshots described in Note 1.2.

**Fig. 2.**
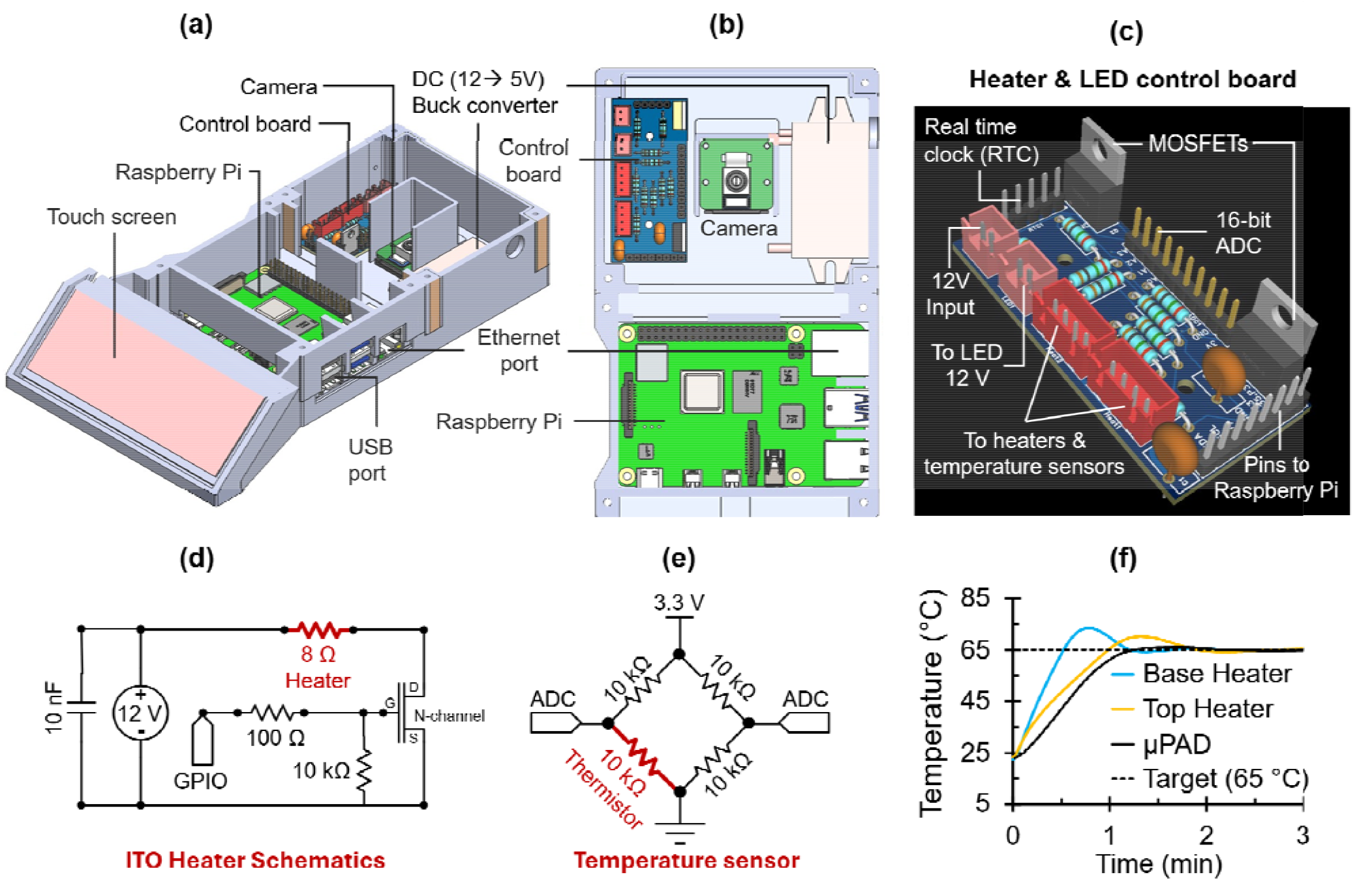
Electronics design and heater control of the ThermiQuant™ VitroMini. (A) Sectional isometric view of the instrument base showing the control board, camera module, and Raspberry Pi 4B controller. (B) Flattened top view of the assembly shown in A. (C) Schematic of the custom-designed printed-circuit-board (PCB) control module integrating heater, illumination, and sensor interfaces. (D) Electronic schematic of the heater-control circuit. Each heater is powered by 12 V and regulated by an n-channel MOSFET driven by a pulse-width-modulated (PWM) signal from the Raspberry Pi’s GPIO pins, which modulates the duty cycle to control the average power delivered to the 8 Ω resistive ITO heater. (E) Temperature-sensing circuit based on a 10 kΩ thermistor arranged in a Wheatstone-bridge configuration for precise temperature measurement and feedback. (F) Representative temperature profiles showing independent proportional–integral–derivative (PID)-regulated control of the top and base heaters, along with the corresponding temperature measured at the µPAD location inside the cartridge when sandwiched between the two heaters during heating from room temperature (23 °C) to 65 °C.

Each indium tin oxide (ITO) heater was independently regulated using a proportional–integral–derivative (PID) control loop implemented through Python script (“VitroMini.py”, see SI) on the Raspberry Pi, with optimized gains of P = 2.0, I = 0.5, and D = 1.0 for both heaters. The schematic of the heater control circuitry is shown in Figure 2D. The resistance of each ITO heater measured 8 Ω after assembly. One terminal of the heater was connected to the 12 V supply, while the other was connected to the drain of an n-channel MOSFET, whose source was grounded. A 10 nF capacitor was placed between the 12 V and ground lines to suppress high-frequency electrical noise. The MOSFET gate was connected in series to a 100 Ω resistor, which limited inrush current and protected the Raspberry Pi’s GPIO pin from voltage spikes. A 10 kΩ pull-down resistor connected the MOSFET gate to ground to ensure the gate remained in the off state when no control signal was applied. Temperature measurement was performed using a Wheatstone bridge circuit (Figure 2E) comprising three 10 kΩ resistors and one arm containing the 10 kΩ thermistor from the heater assembly. The Wheatstone-bridge was powered by the 3.3 V rail of the Raspberry Pi, and the differential voltage across the bridge was read by a 16-bit ADS1115 analog to digital converter (ADC, Amazon, B0CNV9G4K1). The thermistor voltage was converted to temperature using the Beta-parameter equation:

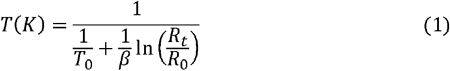

where *T*(*K*) is the absolute temperature in kelvins, *T*_0_ is the reference temperature of 298.15 *K, R*_*t*_ is the measured thermistor resistance (Ω) at temperature *T, R*_0_ is the nominal resistance at 25 °C which is 10 kΩ and β is the thermistor Beta coefficient, 3950 Kfor the MF55 series thermistors.

For illumination, two LED panels were custom-designed and mounted on opposite side walls inside camera enclosure, as shown in Figure 1B. The LED panels were powered directly from the 12 V line of the power management module. For image acquisition, an autofocus IMX519 camera module (Arducam, B0371) was integrated into the system. Custom Raspberry Pi software (“VitroMini.py”, see SI) controlled time-lapse image capture at camera gain of 2,4,6,8 and exposure time of 1 ms while maintaining isothermal conditions within the heater module. Images were recorded at a resolution of 640 × 480 pixels every 30 seconds, with the first frame discarded to allow the camera to stabilize.

### µPAD fabrication and LAMP reagent preparation

µPADs were fabricated following previously published methods^20^, and the colorimetric LAMP reagent composition was based on previously published formulations with primers listed in Table S3^20,27^; both are described in Note 1.3 of the SI.

### Cartridge fabrication, LAMP reagent loading, and assembly of µPADs onto cartridge

An acrylic cartridge was fabricated to accommodate twelve µPADs arranged in a circular pattern (26 mm diameter), designed to minimize reagent evaporation during 1 h incubation at 65 °C. The detailed methods are provided in Note 1.4 of SI.

### Synthetic *orf7ab* DNA preparation and copy number calibration

Synthetic DNA corresponding to the SARS-CoV-2 *orf7ab* target sequence (Table S4; NCBI Reference Sequence: NC_045512.2) was quantified using digital PCR (dPCR) following a previously reported protocol (primers in Table S5), with details provided in Note 1.5 of SI.

### Paper LAMP reaction test in ThermiQuant™ VitroMini and in ThermiQuant™ MegaScan

Three configurations of paper-based LAMP experiments were conducted to evaluate the performance of the ThermiQuant™ VitroMini: (i) dual-sided heating, (ii) single-sided heating, and (iii) operation with a previously reported ThermiQuant™ MegaScan, a water bath based volumetric heating instrument^20^.

For dual-sided heating experiments, four cartridges were prepared, as each cartridge could accommodate only twelve µPAADs. Each µPADs had been preloaded with the LAMP final mix and dried. The dPCR-quantified synthetic DNA was serially diluted to generate a range of concentrations (in copies/reaction) with no-template controls (NTCs) and used in three batches with three technical replicates. Batch 1: 10^6^, 10^5^, 10^4^, NTC1; Batch 2: 500, 250, 100, NTC2; Batch 3: 10^3^, 50, 25, NTC3. One additional batch was included for comparing against single-sided heating (10^3^, 100, 50, NTC). A 7.5 µL aliquot from each dilution was pipetted onto the µPADs, with three technical replicates per concentration. Each batch also included triplicate NTCs (NTC1, NTC2, NTC3), where an equivalent volume of nuclease-free water was pipetted to rehydrate the µPADs. After sample loading, the cartridge was sealed with PCR plate sealing film and placed inside the ThermiQuant™ VitroMini. The lid containing the top heater was then closed, and the onboard software initiated automatic temperature control for 1 h at 65 °C. Time-lapse images were captured at different camera gain settings of 2, 4, 6, and 8 and time-lapse imaging every 30 seconds throughout the incubation. The cartridge was retrieved upon completion of the run and disposed in biohazard bag.

The single-sided heating experiments followed the same protocol as above, with three modifications: (i) electrical connections to the base heater were disconnected, while temperature sensors remained connected to prevent software errors; (ii) two concentration groups were tested at a top heater setpoint of 65 °C: (10^5^, 10^4^, 10^3^ copies/reaction, and NTC) and (10^3^, 100, 50 copies/reaction and NTC). The latter was also tested at a higher setpoint of 70 °C; and (iii) each condition was run with three technical replicates.

A third experiment was conducted using the ThermiQuant™ MegaScan instrument. The experimental procedure was largely identical to the VitroMini protocol (first configuration), with the following adjustments: (i) cartridges were immersed in a water bath maintained at 65 °C and (ii) time-lapse images were captured using the MegaScan’s imaging system (flatbed scanner). The same concentration range (10^5^, 10^4^, 10^3^ copies, and NTC) and triplicate setup were used to ensure comparability across test configurations.

### Determination of LOD95 and LOQ

The definitions of LOD95 and LOQ used in this study are provided in SI Note 1.6.

### Software Image analysis workflow

Time-lapse images captured by the ThermiQuant™ VitroMini were processed using an updated version of the custom Python-based Amplimetrics™ software (Amplimetrics™ 2.0, see SI). Detailed user guide for the software is provided in SI Note 1.2.

### Heater characterization

evaluated through a series of tests assessing heat distribution, uniformity, and stability. First, a TOPDON TC001 thermal camera (Amazon, B0B7LMB22Q), was used to visualize temperature profiles across both ITO-coated heater plates (Figure 1D), confirming the overall heat distribution pattern. Next, four K-type thermocouples were positioned diagonally across an acrylic cartridge at 14 mm intervals (Figure 3A). Temperature readings were continuously recorded for 20 min using a multichannel data logger (AZ Instruments, Amazon, USA) at 1 second interval to determine equilibration time and temperature spatial variation.

**Fig. 3.**
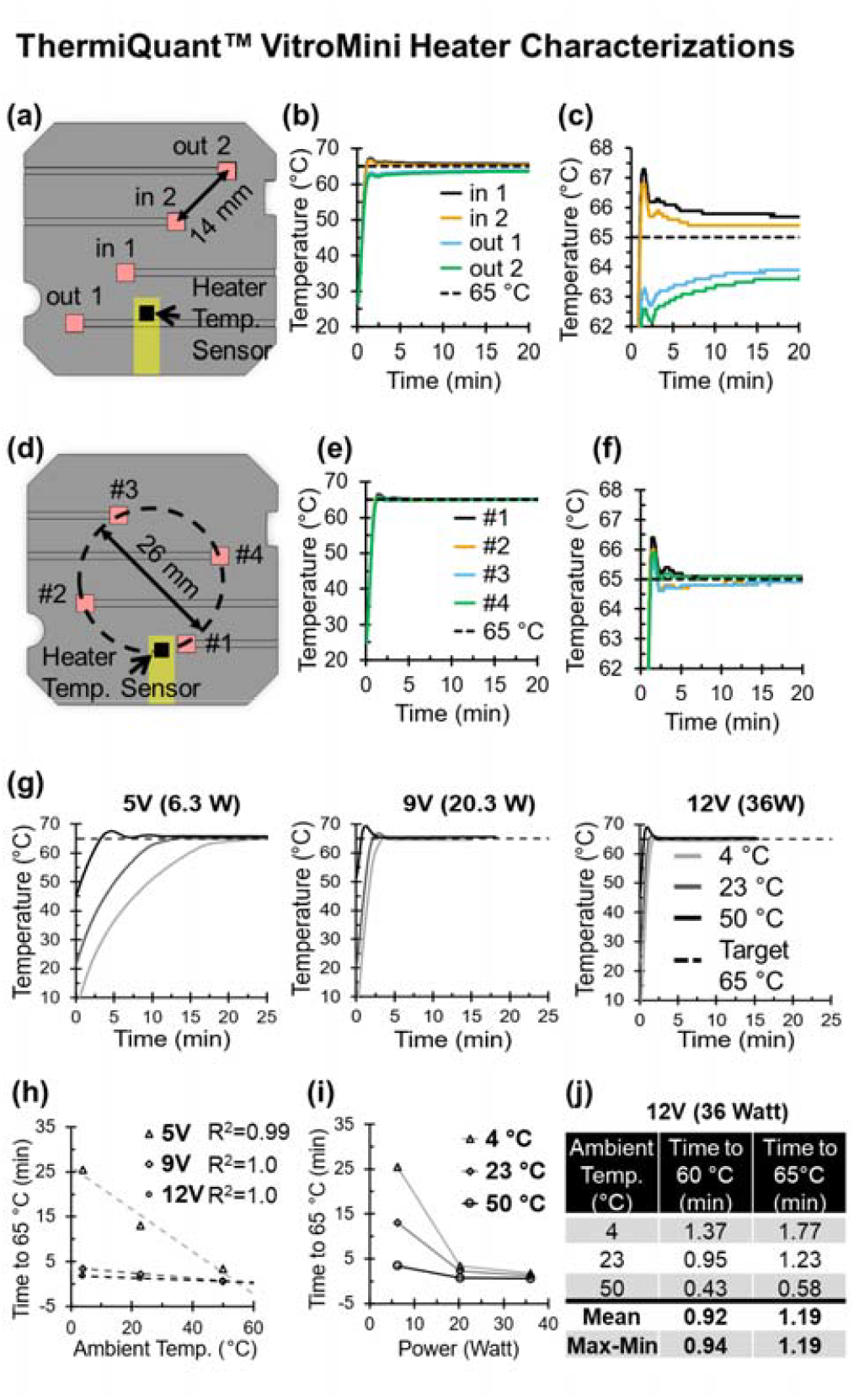
Characterization and optimization of dual-sided heater performance under varying ambient temperatures and power inputs. (A) Test cartridge containing four thermistor probes, two inner (in) and two outer (out), positioned diagonally 14 mm apart and sandwiched between the dual heaters. (B) Temperature-time response recorded at each probe location with (C) magnified view. (D) Optimized probe configuration with four sensors positioned equidistantly (13 mm) from the center along the heater perimeter. (E) Corresponding temperature-time response at each probe position with (F) magnified view. (G) Average temperature measured across four equidistant probes (13 mm from center) at ambient temperatures of 4, 23, and 50 °C under combined power inputs of 5 V (6.3 W, left), 9 V (20.3 W, middle), and 12 V (36 W, right). (H) Time required to reach 65 °C from each ambient condition plotted against ambient temperature with linear regression fit (5V: y = −0.47x + 26.1; 9V: y = −0.059x + 3.6; 12V: y = −0.026 +1.85). (I) Time to first reach 65 °C plotted as a function of total power dissipated by both heaters. (J) Summary table showing times required to reach 60 °C and 65 °C from each ambient condition under 12 V (36 W) operation. The 12 V configuration was selected as the final operating condition for the ThermiQuant™ VitroMini system.

To further assess temperature uniformity under realistic operating conditions, the thermocouples were reconfigured in a circular arrangement 13 mm from the cartridge center (Figure 4A). Using this circular configuration, the system was tested under three ambient conditions: 4 °C (refrigerated), 23 °C (room temperature), and 50 °C (oven) and heater operated at 5 V, 9 V, and 12 V to evaluate heating performance and thermal stability.

**Fig. 4.**
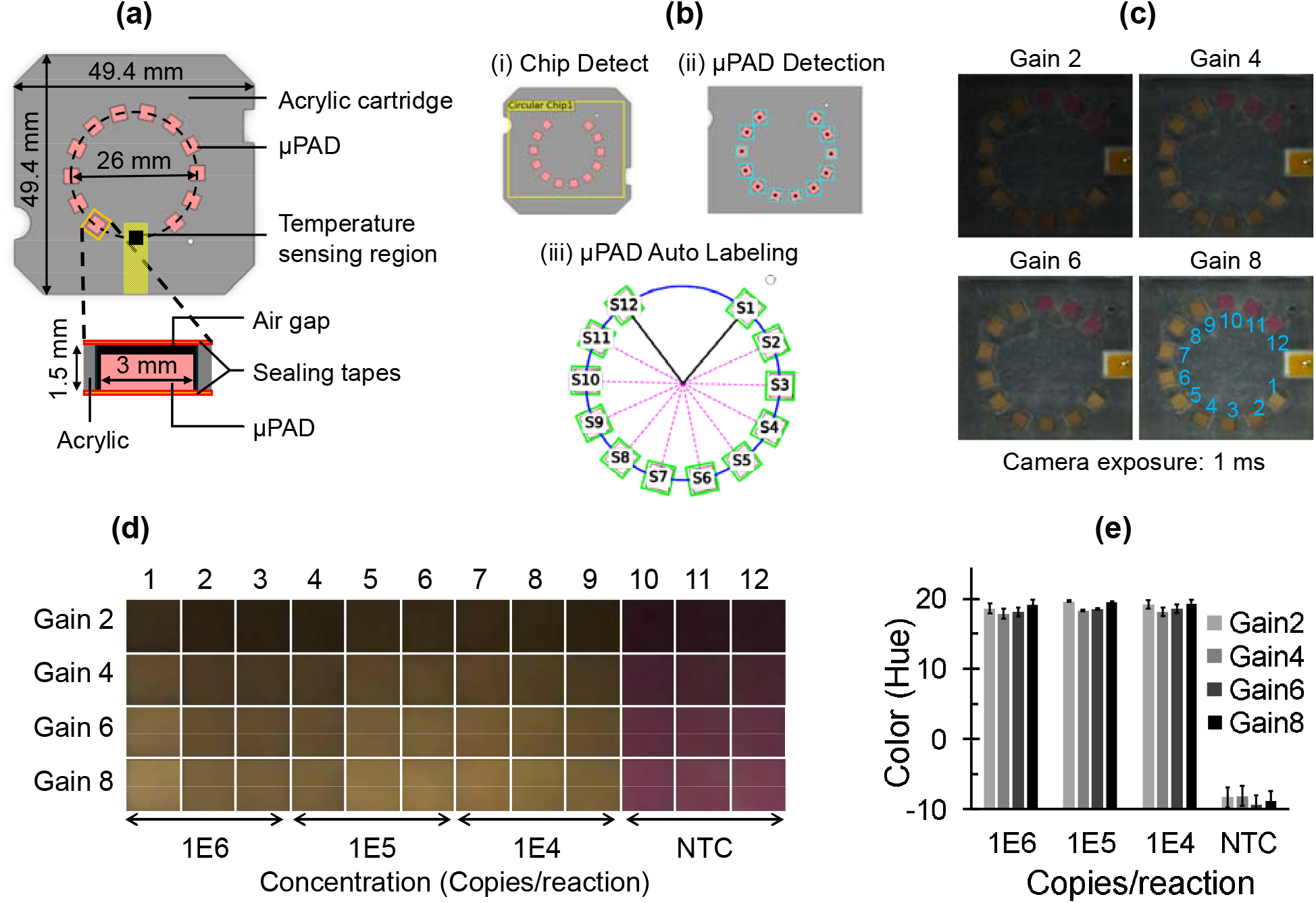
Characterization of the imaging and color-detection performance of the ThermiQuant™ VitroMini. (A) Laser-cut cartridge containing 12 µPADs arranged in a circular layout. Inset: orthogonal cross-section of a 3 × 3 mm µPAD embedded within a 1.5-mm-thick acrylic cartridge and sealed with transparent adhesive film on both sides to prevent evaporation. (B) Automated image-processing workflow comprising three sequential steps: (i) chip detection, (ii) identification of µPAD positions and centroids, and (iii) calculation of median object size followed by reorientation, bounding, and automatic numbering of all 12 µPADs. (C) Endpoint (60 min) µPAD-LAMP images captured at varying camera gain settings with a fixed exposure time of 1 ms. (D) Cropped image patches from C corresponding to 10^6^, 10^5^, 10^4^ copies/reaction, and a no-template control (NTC), each shown with three technical replicates per concentration. (E) Bar chart showing color analysis based on rescaled hue values (Mean ± SD) for the cropped µPAD patches in D.

## Results

### ThermiQuant™ VitroMini is a portable instrument for colorimetric loop-mediated isothermal amplification (LAMP) analysis

We developed the ThermiQuant™ VitroMini as a compact, dual-sided heating instrument for quantitative colorimetric LAMP analysis on µPADs (Figures 1–2). The device measures 22.5 × 13 × 1.5 cm, weighs 0.95 kg, and functions as a portable, stand-alone analytical platform (Figure 1A–B). Its heating module integrates two ITO-coated glass heaters (8 Ω each) attached to distinct substrates: a 1 mm-thick aluminum plate for the top heater and a 1.5 mm-thick acrylic plate for the base heater (Figure 1C). The thermal profile (Figure 1D) showed that the aluminum-backed heater produced a uniform temperature distribution, whereas the acrylic-backed heater displayed a radial gradient with the center hotter than the periphery, due to their contrasting thermal conductivities (~205 W ^−1^K^−1^-for aluminum vs. ~0.2 W m^−1^-K^−1^-for acrylic). This asymmetric thermal behavior can compromise isothermal conditions during LAMP reactions when multiple µPADs are distributed across the heater surface. To address this issue, we arranged twelve µPADs in a circular layout aligned with the heater’s radial isotherms, achieving temperature uniformity with a standard deviation of less than ±0.5 °C at the 65 °C setpoint. This configuration provided both stable isothermal performance for LAMP amplification and a transparent optical window for real-time imaging during incubation.

To ensure precise and independent temperature regulation of both heating surfaces, we implemented a custom electronic control system and evaluated its performance. Each heater incorporated a 10K B3950 NTC thermistor positioned 13 mm from the center and connected in a Wheatstone-bridge circuit for temperature sensing (Figure 2E). The Wheatstone bridge converts thermistor resistance changes into proportional voltage signals while minimizing the effects of wire resistance, supply voltage fluctuation, and minor thermal drift in circuit components. It also enhances the linearity of the voltage–temperature response within the typical 40–80 °C range, encompassing the 65 °C operating setpoint. Independent PID controllers implemented on a custom PCB (Figure 2C) regulate both heaters using 3.3 V PWM signals from the Raspberry Pi GPIO pins, which drive 12 V through n-channel MOSFETs to modulate power delivery (Figure 2D). As shown in Figure 2F, both heaters reached 65 °C from 23 °C within 2 min with a small initial overshoot (< 5 °C). The base heater reached its setpoint about 0.5 min faster than the top due to its lower thermal mass. In contrast, the temperature at the µPAD plane showed a smooth, damped response with minimal overshoot (<1 °C) and reached a stable 65 °C within 1.5 min. This damping behavior results from the transient heat transfer through multiple layers, glass, aluminum, acrylic, and the cartridge, each with distinct thermal conductivity and heat capacity. As heat propagates through these materials, energy absorption and delayed heat conduction smooth rapid initial heater overshoot, producing a stable and uniform isothermal environment at the µPAD location.

With precise temperature regulation established, the VitroMini provides a streamlined end-to-end workflow that semi-automates incubation, imaging, and quantitative analysis for µPAD-based colorimetric LAMP assays. The operational sequence (Figure 1E) involves rehydrating pre-dried LAMP reagents on µPADs with 7.5 µL of DNA sample, sealing the cartridge with transparent film, and placing it between the two heaters. The current workflow assumes that nucleic acid input is obtained through minimally processed samples or upstream preparation steps performed outside the instrument, enabling compatibility with a wide range of sample types and preprocessing requirements. Once the onboard software is initiated, the instrument autonomously incubates reactions at 65 °C for 1 h, capturing time-lapse images every 30 s. After each run, the image set is transferred from the VitroMini to a separate computer running the Python-based Amplimetrics™ software. Amplimetrics™ automatically detects the reaction zones, extracts hue values, and generates hue-versus-time amplification curves for quantitative interpretation (Figure 1E, last panel). Beyond basic user actions such as sample loading, cartridge sealing, and image file transfer, all operational steps, including heating, imaging, and analysis, are fully automated, minimizing user intervention and ensuring reproducible, quantitative colorimetric readouts. As the scope of this work focused on the heater design and imaging performance, image analysis was performed off-board; however, our previous ThermiQuant™ AquaStream^22^ platform have demonstrated the feasibility of on-board processing on a Raspberry Pi 4B. Future iterations of VitroMini will incorporate expanded embedded software, including optical character recognition (OCR), barcode, or QR-code–based cartridge identification and automated µPAD localization, and integration with cartridge-based sample preparation and distribution modules.

### Dual-sided heating with circular µPAD layout maintains ±0.5 °C uniformity from 4 °C to 50 °C ambient conditions

We evaluated the dual-sided ITO heater system of ThermiQuant™ VitroMini (Figure 1C) to assess its thermal uniformity and stability under varying ambient conditions and power inputs (Figure 3). A custom test cartridge equipped with four thermocouple probes–two inner (in) and two outer (out) positioned diagonally 14 mm apart–was sandwiched between the heaters (Figure 3A). In this configuration, the inner probes consistently recorded higher temperatures than the outer probes (Figure 3B and 3C). Even after 20 min of heating, a residual 2–3 °C difference remained, indicating a non-uniform temperature distribution across different µPAD locations within the cartridge. We attribute this temperature gradient to the thermal non-uniformity of the base heater, where the low-conductivity acrylic layer produces a radial heat distribution pattern (Figure 1D). The aluminum-backed top heater, with high thermal conductivity (::205 W m^-1^ K^-1^), distributes heat efficiently, whereas the acrylic–ITO base heater, with nearly 1,000-fold lower thermal conductivity (≈0.2 W m^-1^ K^-1^), limits lateral heat transfer, causing faster heat loss near the outer edge than in the center. This mismatch in thermal behavior results in a center-hotter-than-edge temperature profile across the test cartridge.

Based on these observations, we hypothesized that if the µPADs were aligned in a circular arrangement matching the radial heat distribution, a more uniform temperature could be achieved across different µPAD locations. To test this, we redesigned the test cartridge by positioning four thermocouples equidistantly (13 mm from the center) along a circular path (Figure 3D). In this configuration, both heaters reached 65 °C within 2 min, and the standard deviation across all probes was < 0.5 °C (Figure 3E and 3F). These results confirmed that aligning the µPADs with the heater’s radial isotherms minimizes lateral temperature gradients, providing a uniform thermal field suitable for LAMP reaction. This circular configuration was therefore adopted as the final cartridge design (Figure 1E & 4A).

Using the optimized circular four-probe cartridge, we next examined how the time required to reach the target temperature (65 °C) varied under different ambient conditions and power inputs (Figure 3G–J). The instrument was placed in controlled environments of 4 °C (freezer), 23 °C (room temperature), and 50 °C (oven) and allowed to equilibrate to each test ambient condition prior to heating. The heaters were operated at standard DC voltage levels of 5 V (6.3 W), 9 V (20.3 W), and 12 V (36 W) and regulated by independent PID controllers as described previously (Figure 2C–F). Each ITO heater had a resistance (R) of 8 Ω; since the two heaters were powered in parallel, the effective resistance was 4 Ω, resulting in power dissipation of V^2^ /R = 6.3, 20.3, and 36 W for 5, 9, and 12 V, respectively. The average temperature from the four probes was used to evaluate heating performance under each ambient condition.

Across all tests, the temperature probes consistently reached the 65 °C setpoint, although the time required varied substantially with both power input and ambient temperature. At the lowest voltage (5 V) and coldest condition (4 °C), the cartridge required 25.5 min to reach 65 °C, whereas at the highest voltage (12 V) under the same 4 °C condition, it reached the target temperature in only 1.77 min. The relationship between time-to-setpoint and ambient temperature, shown in Figure 3H, was highly linear, with coefficients of determination (R^2^) of 0.99 for 5 V and 1.00 for both 9 V and 12 V. This strong correlation indicates that, when voltage and heater resistance are fixed, the time required to reach the setpoint can be accurately predicted for each ambient temperature.

In contrast, the relationship between time-to-setpoint and total power dissipation was nonlinear (Figure 3I). For example, at 4 °C, the time to reach the setpoint decreased sharply from 25.5 min at 5 V (6.3 W) to 3.4 min at 9 V (20.3 W) and 1.77 min at 12 V (36 W), exhibiting an approximately exponential decline in heating time with increasing power. This nonlinear behavior indicates that higher power inputs more effectively overcome ambient heat losses, enabling faster net heat accumulation within the cartridge.. As summarized in Figure 3J, at 12 V the system reached 60 °C an**d** 65 °C in 0.92 ± 0.94 min and 1.19 ± 1.19 min, respectively, across all ambient conditions (where “±”denotes the maximum-minimum range, not the standard deviation). Because the 12 V (36 W) configuration provided the fastest and most consistent heating performance across the full 4 to 50 °C ambient range, it was selected as the standard operating voltage for the ThermiQuant™ VitroMini system.

### Automated µPAD imaging and hue-based color analysis remain consistent across illumination conditions

We evaluated the imaging and color-analysis performance of the ThermiQuant™ VitroMini system using phenol-red-based colorimetric LAMP reactions on µPADs (Figure 4). We fabricateed a 12-µPAD cartridge with µPADs positioned along a 26 mm-diameter circle, aligned with the uniform thermal zone of the dual heaters (Figure 4A). Each µPAD contained pre-dried LAMP reagents that swelled from approximately 0.8 mm to 1.2 mm in thickness after rehydration with 7.5 µL of DNA sample. The µPADs were sealed within a 1.5 mm-thick acrylic cartridge using transparent adhesive PCR film to prevent evaporation while maintaining optical clarity for full-field imaging during incubation. This circular layout ensured even temperature distribution (within ±0.5 °C) and consistent imaging throughout the one-hour LAMP reaction.

We developed an automated image-processing pipeline to detect and track all 12 µPADs in the first frame (Figure 4B). The algorithm detected the cartridge, identified µPAD centroids, and generated bounding boxes with automatic numbering, eliminating manual region selection. In all repeated experiments, the detection algorithm consistently identified all µPADs with 100 % accuracy, providing reliable positional alignment for quantitative color analysis.

Hue-based color metrics are inherently robust to illumination variation because the hue channel captures chromatic information separately of intensity and saturation. To verify this behavior in our system, we imaged phenol-red-based colorimetric LAMP reactions using three DNA concentrations (10^6^, 10^5^, and 10^4^ copies/reaction) and NTCs, each with three technical replicates. Endpoint images (60 min) were captured at multiple camera gain settings (using single LED panel) while keeping the exposure time fixed at 1 ms (Figure 4C). Rather than adjusting LED brightness directly, we modulated the camera gain in 5-second intervals to simulate illumination changes through a software-controlled approach. As shown in Figure 4C, µPAD visibility decreased at lower gains (e.g., gain 2–4), making it difficult to visually distinguish between positive (yellow) and negative (red) reactions. Despite this issue, the image-processing algorithm correctly detected and labeled all µPADs at every gain level (Figure 4D). More importantly, hue analysis (Figure 4E) showed that hue values for each concentration group remained consistent within ±2 hue units across all gain settings. This result confirms that the hue channel is minimally affected by illumination variation under front-illumination conditions and that hue-based analysis inherently compensates for moderate differences in illumination intensity or camera sensitivity. Consequently, small inter-instrument variations in LED brightness or optical calibration are unlikely to affect the accuracy of colorimetric LAMP evaluation. Although any gain could be used for automated analysis, we selected gain 8 as the default setting when using single LED panel (as in this experiment) and gain 4 (with exposure time 10 ms) when using two LED panels (implemented in a later system revision) to provide optimal visual contrast for human observation while maintaining full algorithmic accuracy.

### ThermiQuant™ VitroMini enables quantitative colorimetric LAMP detection with LOD95 of 34 copies/reaction and LOQ of 1000 copies/reaction

We next evaluated the analytical performance of the ThermiQuant™ VitroMini for quantitative colorimetric LAMP analysis on µPADs (Figure S4). The LAMP assay chemistry used here has been previously implemented for reverse transcription LAMP (RT-LAMP) for SARS-CoV-2 RNA using clinical nasopharyngeal swab samples^20^. In this study, we used synthetic DNA to isolate and evaluate the thermal and imaging performance of the device without introducing additional variability from upstream sample preparation or reverse transcription efficiency. We tested serial dilutions of synthetic SARS-CoV-2 *orf7ab* DNA ranging from 10^6^ to 25 copies per reaction, using three technical replicates per concentration and nine NTCs obtained from three independent runs. Figure S4A shows representative cropped images captured at 10 min intervals over a 60 min reaction period. The color transition from red (negative) to yellow (positive) became progressively slower with decreasing template DNA concentration, reflecting concentration-dependent amplification kinetics.

We processed the time-lapse images using Amplimetrics™ software, which generated hue-versus-time trajectories for each µPAD (Figure S4B). Positive reactions exhibited characteristic sigmoidal curves, whereas negative reactions remained linear. We set a 10-hue-unit threshold to classify reactions as positive or negative. Figure S4C shows the probit fit used to determine the limit of detection at 95% probability (LOD95)^28^, yielding 34 copies per reaction.

We next analyzed the hue-time trajectories from positive reactions to construct a quantitative calibration curve. For each amplification curve, we calculated the first derivative of hue versus time and fitted it to a Voigt function. We then performed a second-derivative analysis on the Voigt fit to identify the time of maximum slope change, following our previously reported approach^20^. We defined this time point as the quantification time (Tq), corresponding to the inflection point in the amplification curve’s exponential-linear phase. Figure S4D shows the coefficient of variation (CV) of Tq at each standard concentration, with the limit of quantification (LOQ = 1,000 copies/reaction) defined as the lowest concentration at which CV is below 10%. This threshold was selected based on prior reports indicating that LAMP assays typically exhibit acceptable repeatability within a CV range of 5–10% across reactions and runs^29–31^. Figure S5A summarizes the individual Tq values along with their means and standard deviations (SD) across all tested concentrations. The resulting Tq-log_10_ concentration relationship (Figure S5B) displayed a strong log-linear correlation (R^2^ = 0.95).

The observed LOD95 of 34 copies per reaction agrees with our previously reported results using the same LAMP chemistry and *orf7ab* target^20,22^. These findings confirm that VitroMini achieves analytical performance comparable to our earlier volumetric water-bath systems (± 0.5 °C temperature uniformity) while highlighting the inherent heterogeneity of LAMP reaction kinetics near and below the LOQ, where stochastic amplification introduces greater variability in Tq among replicates.

### Dual-sided heating achieves volumetric-equivalent LAMP performance and eliminates condensation artifacts

We next performed two sets of comparisons to evaluate the performance of ThermiQuant™ VitroMini. First, we benchmarked its dual-sided heating configuration against our previously reported ThermiQuant™ MegaScan volumetric water-bath system to assess equivalence in quantitative LAMP performance. Second, we compared VitroMini’s dual- and single-sided heating modes to determine how heating geometry influences reaction uniformity and reproducibility (Figure 5).

**Fig. 5.**
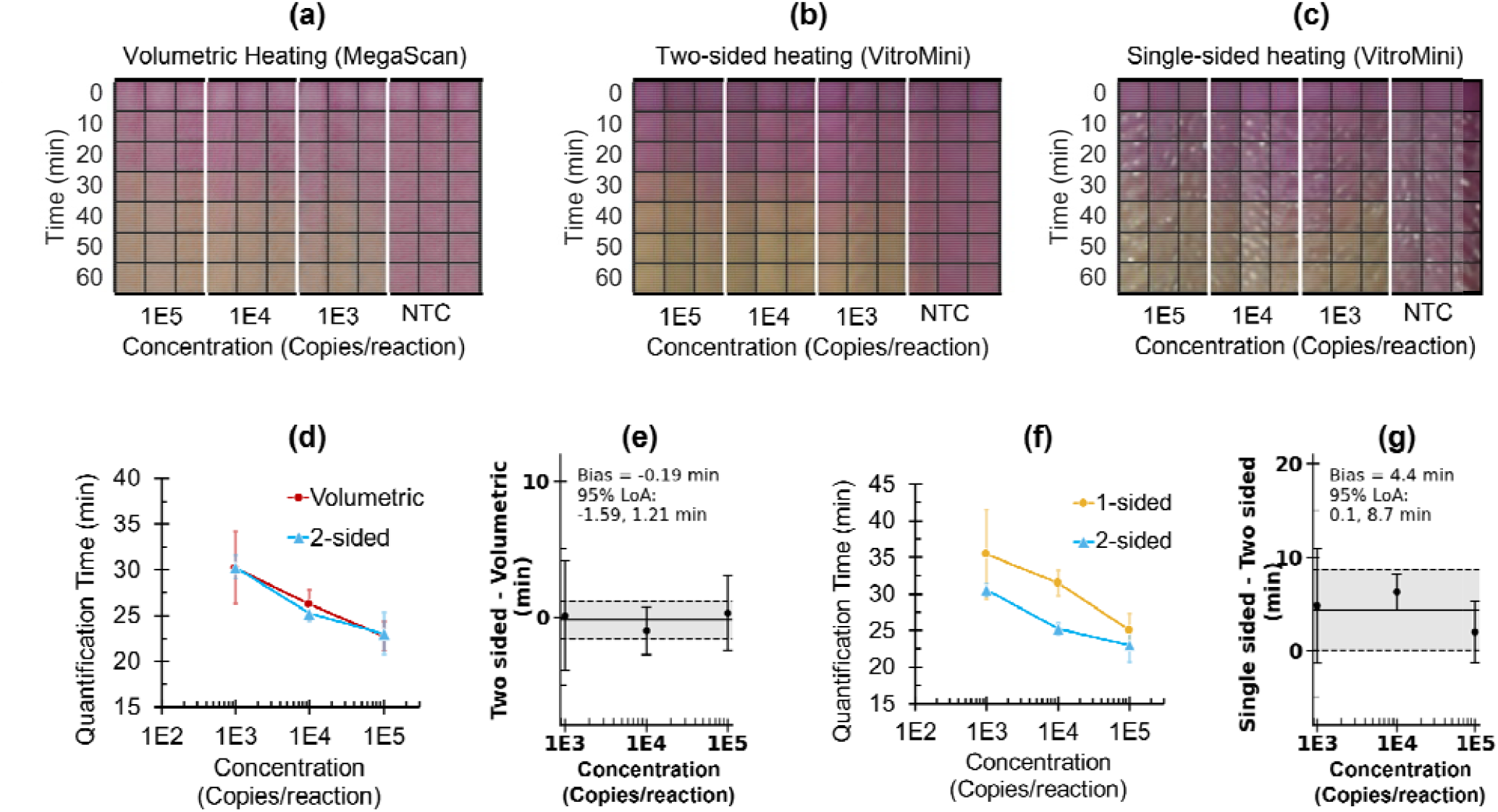
Cross platform comparison of colorimetric LAMP performance across different heating config rations Colorimetric LAMP reactions were performed using three synthetic A concentrations 1E5, 1E4, an 1E3 copies per reaction, each with triplicate replicates except 1E3 which has n=6 of the SARS-CoV-2 *orf*7*ab* target sequence. (A) Time lapse images capture using the ThermiQuant™ MegaScan, which employs volumetric heating via a water bath an high-resolution flatbed scanning. (B) Corresponding images capture using the ThermiQuant™ VitroMini this work with dual-sided heating and (C) single-sided heating. (D) Quantification time (Tq) versus DNA concentration for volumetric and dual-sided heating configurations. (E) Bland-Altman residual plot comparing differences in Tq between dual-sided and volumetric heating. (F) Tq versus DNA concentration for single and dual-sided heating configurations. (G) Bland-Altman residual plot comparing differences in Tq between single- and dual-sided heating. Error bars in the Tq versus concentration plots show mean ± SD. In the Bland-Altman plots, error bars represent the propagate standard deviation of the difference between the two measurements, calculated as 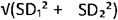.

Representative time-stamped images of µPAD-LAMP reactions (10^5^, 10^4^, and 10^3^ copies/reaction, each in triplicate) are shown in Figure 5A–C for volumetric heating (MegaScan)^20^, VitroMini (dual-sided heating, this work), and VitroMini (single-sided heating, this work), respectively. The MegaScan and dual-sided VitroMini produced visually consistent amplification patterns and uniform reaction endpoints as well as their Tq values were similar across the tested concentration (Figures 5D-E), whereas the single-sided configuration displayed uneven color development and localized condensation artifacts on the µPAD surface (Figure 5). These artifacts, absent in the dual-sided and volumetric systems, arose from vertical temperature gradients between the µPAD (in contact with the heater and its ambient exposed surface near the optical window). The resulting surface cooling at the colder end led to water vapor condensation within the first 10 minutes of incubation (Figure 5C). Such condensed tiny water droplets can locally concentrate or dilute reagents, producing spatially heterogeneous color change and reduced reproducibility in single-sided heating setups.

To further assess the impact of heating geometry on performance near the LOD, we conducted colorimetric µPAD-LAMP assays at three input concentrations (1E3, 1E2, 5E1 copies per reaction) and NTCs, using cartridges of two thicknesses (1.5 mm and 2.0 mm) (Figure S6). At 1E3 copies per reaction, all replicates yielded positive amplification across both heating modes and cartridge thicknesses. However, at lower concentrations (1E2 and 5E1 copies per reaction), single-sided heating exhibited incomplete yellow color development with residual red regions indicative of partial amplification. This behavior was observed in both cartridge formats, with improved performance in the 1.5 mm cartridge compared to the 2.0 mm cartridge. Hue-versus-time trajectories further showed that dual-sided heating maintained consistent amplification kinetics across replicates, whereas single-sided heating exhibited reduced amplification near the LOD (Figure S6B).

Quantitative comparison of amplification kinetics confirmed these observations. Tq versus input concentration (Figure 5D) and Bland-Altman analysis (Figure 5E) showed that dual-sided heating closely matched volumetric heating, with a mean bias of −0.19 min and 95% limits of agreement (−1.59 to 1.21 min), indicating near-equivalent performance. In contrast, single-sided heating produced systematically delayed Tq values and increased variability (Figure 5F), with a negative bias of up to −4.43 min and substantially broader limits of agreement (Figure 5G), reflecting reduced reproducibility. Together, these results demonstrate that dual-sided heating minimizes condensation artifacts and achieves volumetric-equivalent performance, while single-sided heating degrades reproducibility, particularly near LOD.

### Through-thickness thermal gradients in single-sided heating drive condensation and degrade assay performance

To understand the origin of the observed LAMP performance differences between single sided and dual sided heating, we quantified through-thickness thermal gradients under single-sided, dual-sided, and elevated-temperature single-sided heating configurations (Figure S7). We used a 2 mm thick acrylic cartridge and positioned thermocouples at the top (near the heater) and at the base (ambient-facing side), with µPADs placed between them. We selected the 2-mm thickness to compensate the ~0.5 mm thermocouple probe head while maintaining a 1.5 mm gap between the probes, and we averaged measurements from two probe pairs.

Under single-sided heating (top heater set to 65 °C), the top surface approached ~63 °C (2 °C below the set point because of temperature difference within the 0.5 mm diameter of the probe as well as temperature offset of top heater) while the base remained at ~59 °C, resulting in a temperature difference of ~4 °C across the 1.5 mm thickness after stabilization (>10 min), corresponding to ~2.7 °C/mm (Figure S7A,C). In contrast, dual-sided heating (both heaters set to 65 °C) reduced this offset to ~1.5 °C (~1 °C/mm) and achieved faster stabilization (~3 min versus >10 min for single-sided heating) (Figure S7B–C). During the initial transient period (<3 min), independent heater PID control produced temperature overshoot and undershoot in both configurations, but the larger and longer-lasting gradients (Figure S7C) in single-sided heating coincide with the early onset (i.e., 10 min) of condensation observed experimentally (Figure 5C).

To compensate for the lower average temperature in single-sided heating (~60–61 °C core temperature at a 65 °C setpoint), we increased the top heater setpoint to 70 °C. This adjustment raised both top and base temperatures (Figure S7D) and brought the estimated core temperature in a 1.5 mm cartridge closer to ~65 °C, although it increased further to ~66 °C after ~15 min (Figure S7E). However, the through-thickness temperature difference remained similar at both 65 °C and 70 °C setpoints (Figure S7F), indicating that increasing the setpoint does not eliminate thermal non-uniformity.

We then evaluated LAMP amplification under these elevated conditions. Hue-versus-time analysis showed that increasing the setpoint to 70 °C reduced amplification efficiency compared to operation at 65 °C across all tested concentrations (1E3, 1E2, and 5E1 copies per reaction) (Figure S7G). These results indicate that increasing the heater setpoint does not mitigate the effects of vertical thermal gradients and can instead degrade assay performance. Notably, although elevating the setpoint brings the average core temperature closer to the nominal 65 °C, it also drives the heater-facing region above the optimal LAMP temperature range (60–65 °C), leading to reduced amplification efficiency. These findings suggest that, in single-sided heating configurations, maintaining the heater setpoint at or below 65 °C is preferable in the design studied here, even if the resulting core temperature is slightly lower (~60–61 °C). In contrast, dual-sided heating achieves the target reaction temperature with a smaller gradient and avoids thermal asymmetry, resulting in consistent amplification independent of ambient conditions.

## Discussion

We demonstrate that the dual-sided ITO heater design in ThermiQuant™ VitroMini bridges the long-standing gap between compact thin-film heaters and high-precision volumetric systems. Volumetric instruments such as FARM-LAMP^16^ and LARI^23^ achieve excellent temperature uniformity (±0.3–0.5 °C) through water- or metal-based heat transfer, but their large size (>3 kg) and high-power requirements (>24 V) limit portability. In contrast, single-sided thin-film platforms, such as UbiNAAT^3^, qcLAMP^29^, and MILP^25^, have gained popularity for point-of-need diagnostics because of their simplicity, compactness, and low cost. However, these systems introduce two persistent limitations: through-thickness thermal gradients and sensitivity to ambient conditions.

In single-sided designs, only one face of the cartridge receives heat, while the opposite face dissipates heat to the environment, producing a vertical temperature gradient across the reaction volume. This through-thickness thermal gradient forces the heater to operate above the nominal reaction temperature to achieve optimal conditions within cartridges typically 1–5 mm thick and fabricated from low-conductivity materials such as acrylic. Our results show that this configuration also introduces condensation artifacts due to cooling at the ambient-facing surface, leading to spatially heterogenous amplification (Figure 5C). In contrast, VitroMini’s dual-sided configuration encloses the cartridge between two independently regulated ITO heaters operating at the same setpoint, minimizing this asymmetry. As a result, the reaction zone equilibrates uniformly to the target temperature, independent of external conditions, and avoids condensation artifacts observed in single-sided systems (Figure 5A–B). Consistent with these observations, dual-sided heating matched volumetric performance with minimal bias (−0.19 min) and tight limits of agreement, whereas single-sided heating exhibited delayed amplification and increased variability (Figure 5D–G).

Importantly, these limitations become more pronounced near the limit of detection. As shown in Figure S6, single-sided heating exhibited reduced amplification efficiency and increased variability at lower input concentrations (1E2 and 5E1 copies per reaction), whereas dual-sided heating maintained consistent performance. This behavior is particularly relevant for diagnostic applications, where reliable detection at low target concentrations is essential.

Attempts to compensate for thermal bias in single-sided systems by increasing the heater setpoint were ineffective. Although elevating the heater to 70 °C increased the average core temperature toward the nominal 65 °C target (Figure S7), it did not reduce the through-thickness gradient and instead degraded amplification performance across all tested concentrations. This outcome reflects a fundamental limitation of single-sided heating: increasing the setpoint to compensate for cooler regions simultaneously drives the heater-facing surface above the optimal LAMP temperature range (60–65 °C), impairing reaction efficiency. These findings indicate that, in single-sided configurations, maintaining the heater setpoint within the optimal reaction range is preferable to attempting compensation via elevated temperatures. In contrast, dual-sided heating eliminates this constraint by achieving uniform temperature distribution without exceeding the optimal reaction range.

Even after eliminating temperature bias, the heaters performance remains influenced by environmental heat loss, which affects the ramp up dynamics (time required to reach the target temperature of 65 °C). We evaluated system behavior across ambient temperatures from 4 °C to 50 °C and found that increasing power dissipation (V^2^ /R) effectively compensated for this loss. The 12 V configuration provided the best balance, achieving consistent heating with a maximum ramp-time variation of only 1.2 minutes between 4 °C and 50 °C ambient conditions. This variation is acceptable given that real-time colorimetric LAMP imaging typically acquires frames at one-minute intervals^16,26,29^. Moreover, the observed strong linear relationship between ramp-up time and ambient temperature at a fixed voltage (Figure 3H) suggests that this behavior can be modeled for predictive thermal control. Future system iterations could leverage this finding by incorporating adaptive feedback, such as an additional thermistor to monitor ambient temperature and dynamically adjust ramp profiles. This approach would be especially valuable for low-voltage, battery-powered configurations (e.g., 5 V operation). In such cases, two design optimizations would be beneficial: (i) using lower-resistance ITO coatings (<8 Ω) to increase power dissipation (for example, reducing resistance to 2 Ω at 5 V would yield approximately fourfold higher heat output), and (ii) implementing a staged heating algorithm that first ramps to ~40 °C, below the activation threshold of Warmstart® Bst polymerase, before proceeding to 65 °C.

Maintaining optical transparency while preserving isothermal uniformity introduced an inherent geometric trade-off. Because the acrylic imaging window has low thermal conductivity, lateral heat spreading was constrained, so we achieved uniform heating only when µPADs were positioned radially around the heater’s center. This design is less compatible with linear strip-style cartridges but aligns naturally with a broad class of µPAD and microfluidic architectures that use circular or concentric layouts for multiplexed reactions, radial flow, or reagent distribution^3,30^. Indeed, centrifugal microfluidic “lab-on-a-disc” systems typically employ disc-shaped cartridges with radial fluidic and reaction chambers for parallel operations^31,32^. For example, a multiplex circular-array continuous-flow PCR chip arranged 12 serpentine channels in an annular layout around the disc^33^. Thus, while the VitroMini design may not suit non-circular cartridge layouts, its circular µPAD arrangement remains a practical and transferable solution for a wide range of multiplexed diagnostic assays that demand transparent, compact, and thermally stable operation.

Nevertheless, the circular heating constraint arises primarily from radial heat loss at the edges of the heater stack. Future design refinements, such as extending the aluminium top frame to form a thermally conductive perimeter around the lower heater, could help suppress edge losses and improve lateral heat distribution. However, such modification must be balanced against constraints related to optical access, heater transparency, and fabrication variability associated with commercially available ITO coated glass substrates. The use of custom-fabricated heaters with improved coating uniformity or alternative transparent substrates (e.g., sapphire-based conductors) may further enhance thermal uniformity in future iterations. Additionally, preheating the enclosure with a small air heater could reduce the temperature gradient between the heater and its surroundings, further mitigating peripheral cooling. With these modifications, the system may better support non-circular or linear cartridge geometries while maintaining the optical access and thermal stability achieved in the current VitroMini design.

Beyond its thermal performance, ThermiQuant™ VitroMini achieves illumination-independent colorimetric analysis through the combination of front illumination and hue-based quantification. Similar findings have been reported in earlier studies that employed hue as an analytical metric for colorimetric LAMP. For example, Nguyen *et al*. (2020)^26^ showed that hue exhibited minimal variability under front illumination, although their study used a different chromogenic dye, hydroxynaphthol blue (HNB), instead of phenol red. In contrast, our previously reported ThermiQuant™ AquaStream system, which used backlit illumination through the reaction zone, displayed systematic hue shifts with changes in camera gain because absorbance dominated over reflectance, causing hue values to vary with gain^22^. In the present study using front illumination, hue values varied by less than ±2 units across a wide range of gain settings, confirming that the hue channel remained stable even when overall image brightness changed markedly (Figure 4C–D). This stability arises because hue isolates chromatic information separately of brightness and saturation, making it intrinsically robust to variations in illumination intensity or sensor gain. Such robustness is particularly important for portable instruments, where illumination uniformity can fluctuate due to power supply variation and LED aging.

Tracking hue trajectory over time extends this robustness into a quantitative analytical framework. The hue–time profiles displayed distinct kinetic patterns, sigmoidal curves for positive reactions and linear traces for negatives, enabling clear discrimination and automated classification. Among positive reactions, the corresponding Tq showed a strong log-linear relationship with input DNA concentration, allowing relative quantification of unknown samples. The measured LOD95 of 34 copies/reaction and linear calibration fit above LOQ of 1,000 copies/reaction matched those obtained from volumetric water-bath systems such as ThermiQuant™ MegaScan and AquaStream, demonstrating that VitroMini maintains laboratory-grade analytical precision despite its compact, low-power design^20,22^. However, as with qLAMP and qPCR, this quantification remains relative, relying on calibration curves, whereas digital PCR (dPCR) achieves absolute quantification through reaction partitioning and Poisson statistics^34,35^. Therefore, ThermiQuant™ VitroMini is best suited for relative quantification and kinetic benchmarking, providing a portable, reproducible platform for assay optimization, reagent validation, and decentralized molecular diagnostics.

## Conclusion

In this study, we introduced ThermiQuant™ VitroMini, a dual-sided indium tin oxide (ITO) thin-film heater that achieves volumetric-level temperature uniformity within a compact, transparent architecture optimized for real-time colorimetric LAMP analysis. The system offers three key advantages: (i) a dual-sided ITO configuration that maintains 65 ± 0.5 °C across ambient conditions from 4 °C to 50 °C with less than 1.2 min variation in ramp-up time; (ii) stable hue-based colorimetric quantification independent of camera gain; and (iii) reliable analytical performance, with quantification possible on and above 1,000 copies/reaction (LOQ) and detection down to LOD of 34 copies/reaction using a synthetic *orf7ab* target. The instrument also has two limitations: (i) its geometric compatibility currently is only useful for circularly arranged µPAD layouts; and (ii) the colorimetric methodology is presently limited to phenol red chemistry. Future work will focus on two directions: (i) further miniaturization toward a palm-sized design and all analysis on board; and (ii) expanded bioanalytical validation beyond synthetic DNA to include diverse field-derived samples, accompanied by multi-user testing to assess robustness and usability.

## Supporting information

Supporting Information Readme

Software

Design Files

## Author contributions

**BR:** Conceptualization, software, data curation, formal analysis, investigation, methodology, validation, visualization, writing – original draft preparation. **GP:** Data curation, formal analysis, investigation, methodology, validation, writing – review and editing. **DN:** Data curation, methodology. **RAM:** Data curation, writing – review and editing. **AB:** Conceptualization, methodology VK: Investigation, methodology. **BA:** Data curation. **CM:** Data curation. **AA:** Methodology. **AG:** Methodology, funding acquisition **JP:** Supervision. funding acquisition. **MSV:** Conceptualization, supervision, project administration, funding acquisition, validation, and writing – review & editing. All authors have reviewed the manuscript and approved the final version.

## Data availability

This article includes Supporting Documents and a Mendeley dataset. All raw data and software used in this study are available on Mendeley Data (10.17632/87rytxs8xy.1).

## Acknowledgements

We are grateful to Dr. Mohamed Kamel for initial preliminary data to guide the design of the device.

## Declaration of generative Al and Al-assisted technologies in the writing process

During the preparation of this work, the authors used Grammarly and ChatGPT to check for grammar errors and improve their academic writing language as well as debugging software. After using this tool/service, the authors reviewed and edited the content as needed and take full responsibility for the content of the publication.

## Copyright claim on the trademark name

ThermiQuant™ and Amplimetrics™ are copyrighted by © Purdue Research Foundation 2024

