## Supporting Information for "A solid-state heater-imager for quantitative evaluation of colorimetric isothermal nucleic acid amplification on paper"

for

**Note 1.1 ThermiQuant™ VitroMini design and assembly**

The ThermiQuant™ VitroMini instrument (shown in Figures 1 and 2) consisted of four primary functional modules: (i) a power management module supplying regulated 12 V and 5 V DC power source; (ii) a computation and control module centered on a Raspberry Pi 4B running custom Python software for temperature regulation, image acquisition, and user interaction through a 5-inch touch-screen display; (iii) a heater module containing two independently controlled heater units operated by a custom-designed printed circuit board (PCB); and (iv) an imaging and illumination module incorporating an autofocus camera and dual LED panels that provided uniform illumination during time-lapse imaging.

All 3D-printed components were designed in SolidWorks (Dassault Systèmes SolidWorks Corp., USA) and fabricated using a Bambu Lab X1 Carbon 3D printer (Bambu Lab, China) with black polycarbonate filament. The heater assembly parts were printed at 100% infill density to ensure mechanical rigidity and thermal stability, whereas all other components were printed at 15% infill density to minimize weight and printing time. All laser cutting and engraving were done on a Glowforge Plus 40 W laser cutter (Glowforge Inc., USA). Cutting 1.5 mm acrylic was carried out at 100% power and speed factor 160, while engraving 0.5 mm-deep channels on the same acrylic was performed at 30% power and speed factor 300. The custom PCB boards were fabricated by JLCPCB (China) and designed using their EasyEDA software platform and manually assembled in house. Table S2 contains bill of materials (BOM).

Power for the ThermiQuant™ VitroMini system was supplied through a 12 V, 5 A DC adapter (Amazon, B0BYVK73BP). The 12 V line directly powered the LED illumination panels and, via n-channel IRLZ44NPBF MOSFETs (Amazon, B0BW8K31BF) to two heaters. A buck converter (Amazon, B0BNQ5JNWZ) stepped down the 12 V supply to 5 V to power the Raspberry Pi 4B (Digikey, 2648-SC0194(9)-ND) computational unit, which in turn shared 5 V with the 5-inch touch-screen monitor (Amazon, B0CRR7XMCQ). The Raspberry Pi’s internally regulated 3.3 V rail supplied power to auxiliary low-voltage circuits, including the DS3231 RTC (Amazon, B01N1LZSK3), 16-bit ADS115 analog-to-digital converter (ADC, Amazon, B0CNV9G4K1), and the Wheatstone bridge circuit (Figure 2E) used for thermistor voltage-drop measurements.

The two heater units were fabricated using indium tin oxide (ITO)-coated glass plates, which functioned as resistive heaters; when voltage was applied across the ITO layer through the busbars, the resulting voltage drop generated heat through resistive heat dissipation. Each heater assembly (Figure 1C) comprised five common components: (i) one 50 × 50 mm, 0.7 mm-thick ITO-coated glass heater (Amazon, B09ZDWFY8Cr); (ii) one 10 kΩ thin-film B3950 thermistor (Amazon, B0BN8K7HPR) for temperature sensing; (iii) a pair of custom-machined copper bus bars (60 × 5 mm); (iv) a pair of 5 mm wide sealing foam tape gaskets cut to match the bus bar length (Amazon, B07L6LGMWC); and (v) a pair of Jst Xh 2.54 four pin connector (Amazon, B01DUC1S14), with two leads for the heater and two for the thermistor. The top and base heater assemblies differed in two components. In the top heater, a 0.5 mm-thick thermal pad (12 W m⁻¹ K⁻1, Amazon, B095H32JFP) with a 5 mm width x 15 mm length cutout manually for thermistor placement was positioned beneath a 1 mm-thick 6061 aluminum sheet (Amazon, B0C4Z167CX) that acted as a heat spreader; the thermistor was bonded to the aluminum surface using double-sided thermal adhesive tape (Amazon, B094VMDTH3). In contrast, the base heater employed a 1.5 mm-thick clear acrylic panel (Amazon, B0922XTTNF) with a 0.5 mm-deep engraving to accommodate the thermistor. The thermistor was bonded using double-sided thermal adhesive tape to the acrylic. Both heater assemblies were housed within 3D-printed enclosures, and the top heater incorporated an additional spring-loaded 3D printed plate with five compression springs of 10 mm outer diameter (Amazon, B0C339Q4X5) to provide uniform pressure when sandwiching a custom cartridge between the heater plates.

**Note 1.2 Video1 transcript and screenshots for “Video1_Instrument_Demo_SD.mp4”**

1. Remove the paper backing from the sealing tape and seal one side of the cartridge.
2. Using vacuum suction, pick up each dried micro pad pre-loaded with LAMP reagents and place micropads into the cartridge. Make sure the pads are correctly aligned before proceeding.
3. Using pipette, add 7.5 microliters of the test sample or negative control into each micro pad. Change tips to avoid cross-contamination.
4. Peel off the second side of the sealing tape and gently seal the cartridge to make it airtight. The sealing helps prevent the micro pads from drying out during the one-hour incubation.
5. Place the cartridge on the heater for incubation. Make sure to align the cartridge notch so it fits perfectly onto the heater base.
6. To run the software, select Start, then Accessories, and open the ITO Heater App. The software will display two windows: an image on the left and a real-time temperature graph for both heaters. Incubate the cartridge for 60 minutes.
7. Once heating is complete, remove the cartridge from the heater. The positive test results appear yellow, while the negative or control results appear red.
8. Open VS Code and choose the Amplimetrics Python environment to process timelapse images.
9. Run the software. When the window opens, select the Image Processing module and choose the folder containing the images. The first image will appear on the screen.
10. Select the chip type as Circular, then choose Generate Metadata. The software will automatically detect and label the micro pads. Next, select Process Images to begin processing. A Hue vs. Time graph will appear on the screen.
11. After image processing, three subfolders will be created inside the parent folder containing the timelapse images. The first subfolder contains cropped image patches of micro pads arranged in a line. The second subfolder, Metadata, includes the labeled image and a JSON file with the coordinates of each micro pad.
12. The third subfolder, Processed Data, contains the label for the cropped linear patch, and a csv file for the raw hue-versus-time data.
13. Next, select the Data Processing module and load the raw hue–time CSV file from the Processed Data folder.
14. The raw and processed data graphs will appear. You can manually adjust the baseline regions, then click Start Normalize and trim the experiment duration. Data is also displayed in the table format.
15. Enter the threshold for manual classification. Blue curves indicate positives; black indicate negatives.
16. Process the derivative to obtain the quantification time. The blue line represents the smoothed data and its sigmoid fit, the pink line shows the first derivative with the Voigt fit, and the red line shows the second derivative. The vertical line indicates the quantification time.
17. Save the quantification-time data as a CSV file. The data also appears in the table. Finally, save the processed hue vs time data.

**Note 1.3 µPAD fabrication and LAMP reagent preparation**

Each µPAD was fabricated using 0.83 mm-thick Grade 222 chromatography paper (Ahlstrom-Munksjö, Finland). Sheets of chromatography paper were first trimmed into 3 mm-wide strips using a leather strip-belt cutting machine (Zhixumm, B0D8THH9LW, Amazon, USA) fitted with a 3 mm blade spacing. A double-sided adhesive film (ARclean® 90178, USA) was placed on top of a 0.076 mm optically clear MELINEX® 454 polyester substrate (Tekra, USA), after which the paper strips were positioned onto the adhesive surface. The assembled sheet was then cut again in a perpendicular direction using the same cutting machine to yield individual 3 × 3 mm µPAD units. An exploded schematic of the µPAD integrated within the cartridge is presented in Figure 4A.

The colorimetric LAMP reagent composition was based on a previously established formulation^1,2^. To prepare 1,000 μL of a 2× LAMP master mix, the following components were combined: 100 μL KCl (1,000 mM; Sigma-Aldrich, P9541), 160 μL MgSO_4_ (100 mM; Sigma-Aldrich, M2773), 280 μL dNTP mix (10 mM; Fisher Scientific, FERR0182), 2.8 μL dUTP (100 mM; Fisher Scientific, FERR0133), 0.4 μL Antarctic Thermolabile UDG (1 U/μL; New England Biolabs, M0372S), 5.4 μL Bst 2.0 DNA polymerase (120 U/μL; New England Biolabs, M0537M), 20 μL phenol red (25 mM; Sigma-Aldrich, P3532), 100 μL Tween-20 (20%; Sigma-Aldrich, P9416), and 331.4 μL nuclease-free water (Fisher Scientific, 43-879-36).

For preparation of 200 μL of the final LAMP reaction mixture, 125 μL of the 2× LAMP mix was combined with 25 μL of 10× primer mix (16 μM FIP/BIP, 2 μM F3/B3, 4 μM LF/LB; final concentrations: 1.6 μM FIP/BIP, 0.2 μM F3/B3, and 0.4 μM LF/LB), 0.67 μL Bst 2.0 DNA polymerase, 1 μL betaine (5 M; Sigma-Aldrich, B03005VL), 3.13 μL bovine serum albumin (BSA, 40 mg/mL; Sigma-Aldrich, A2153), 36.0 μL trehalose (1.75 M; Thermo Scientific Chemicals, 182550250), and 9.2 μL nuclease-free water. The final mixture was adjusted to pH 7.9 with 0.1 M KOH to obtain a uniform red solution. The LAMP primers are listed in Table S3 which were reused from previous publication^1,3^.

A 7.5 μL aliquot of the final LAMP master mix was pipetted onto each µPAD and air-dried for 2 h at room temperature inside a PCR workstation (Mystaire, MY-PCR32). Relative humidity was 20-23%.

**Note 1.4 Cartridge fabrication, LAMP reagent loading, and assembly of µPADs onto cartridge**

The cartridge layout was created in SolidWorks, exported as DXF files, and refined in Adobe Illustrator (Adobe Inc., USA) prior to conversion to SVG format for laser cutting. Acrylic panels (1.5 mm thick; Outus, B09MRFHFTR, Amazon, USA) were cut using a Glowforge Plus laser cutter with optimized settings (100% power, speed factor 160). After laser cutting, cartridges were rinsed with reverse osmosis (RO) water, sequentially treated with 70% ethanol and RNase AWAY™ (Thermo Fisher Scientific, USA), and finally wiped with lint-free Kimwipes® (Kimberly-Clark, USA). For cartridge assembly, one side of the acrylic panel was first sealed using PCR plate sealing adhesive film (Fisher Scientific, AB-0558). The cartridge was then inverted, and µPADs preloaded with the LAMP master mix (and dried) were positioned onto the exposed adhesive surface of the PCR seal using handheld vacuum pickup tool (Amazon, B0D234YZ2R), ensuring firm contact between the µPADs and the adhesive film. Fully assembled cartridges intended for later use were stored in airtight plastic bags at -20 °C. In this study, all cartridges were utilized within 2 days of preparation.

**Note 1.5 Synthetic orf7ab DNA preparation and copy number calibration**

Synthetic DNA corresponding to the SARS-CoV-2 *orf7ab* target sequence (Table S4; NCBI Reference Sequence: NC_045512.2) was purchased as 4 ng lyophilized pellets (IDT, USA) and reconstituted in 40 μL of nuclease-free water (Fisher Scientific, 43-879-36) to obtain the stock solution. Digital PCR (dPCR) primers and probes were the same as previously reported^1^ which were designed using the PrimerQuest™ online tool (https://www.idtdna.com/PrimerQuest/Home/Index) and their sequences are listed in Table S5.

Each dPCR reaction was prepared in a total volume of 40 μL containing 10 μL of 4× Probe PCR Master Mix (Qiagen, 250102; final concentration 1×), 4 μL of 10× primer–probe mixture (final concentration 1×; 0.8 μM forward primer, 0.8 μM reverse primer, and 0.4 μM FAM-labeled probe; see Table S5), 0.5 μL of EcoRI-HF restriction enzyme (New England Biolabs, R3101S), 20.5 μL of nuclease-free water, and 5 μL of DNA template. Reaction mixtures were loaded onto a 26K 24-well Nanoplate (Qiagen, 250001) and analyzed using a QIAcuity One 5-plex dPCR system (Qiagen, 911021).

The amplification program comprised an initial denaturation at 95 °C for 2 min, followed by 40 cycles of 95 °C for 15 s, 55 °C for 15 s, and 60 °C for 30 s. Absolute DNA copy number quantification was performed using the QIAcuity Software Suite (Qiagen). The prepared synthetic DNA stock was stored at –80 °C until further use.

**Note 1.6 Determination of LOD95 and LOQ**

The limit of detection (LOD95) was estimated using probit regression analysis^4,5^. Binary amplification outcomes (positive/negative) were modeled as a function of the log_10_-transformed target concentration. LOD95 was defined as the concentration corresponding to a 95% predicted probability of detection. Given the limited number of replicates per concentration (n=3), this value should be considered a preliminary estimate demonstrating the capability of the instrument to perform and analyze such assays. Comprehensive analytical validation was beyond the scope of this study. For reference, regulatory guidelines (e.g., FDA recommendations) typically require testing at least 20 replicates near the anticipated detection limit and defining the LOD as the concentration at which ≥19 out of 20 replicates yield positive results^6^.

The LOQ was defined as the lowest concentration at or above the LOD for which the CV was ≤10%, with the additional criterion that the two subsequent higher concentrations also met this threshold. CV was calculated only for concentrations with at least three valid replicate measurements. Thus, determination of the LOQ required three consecutive concentrations satisfying the CV threshold, beginning at the candidate LOQ level. Although LOQ is commonly defined as the lowest concentration meeting a predefined precision threshold (e.g., CV ≤10%), under the assumption that variability decreases with increasing concentration, this trend was not consistently observed in our data, with elevated CV values at higher concentrations likely influenced by the limited number of replicates (n=3). Therefore, the additional requirement was introduced to ensure robustness against non-monotonic variability and potential outliers. It should be noted that LOQ definitions are assay- and application-dependent, and the criteria applied here were selected to demonstrate the analytical performance of the device rather than to establish a clinically validated threshold. The 10% CV threshold was selected based on prior studies reporting that LAMP assays typically exhibit acceptable repeatability within a CV range of approximately 5–10%^7–9^.

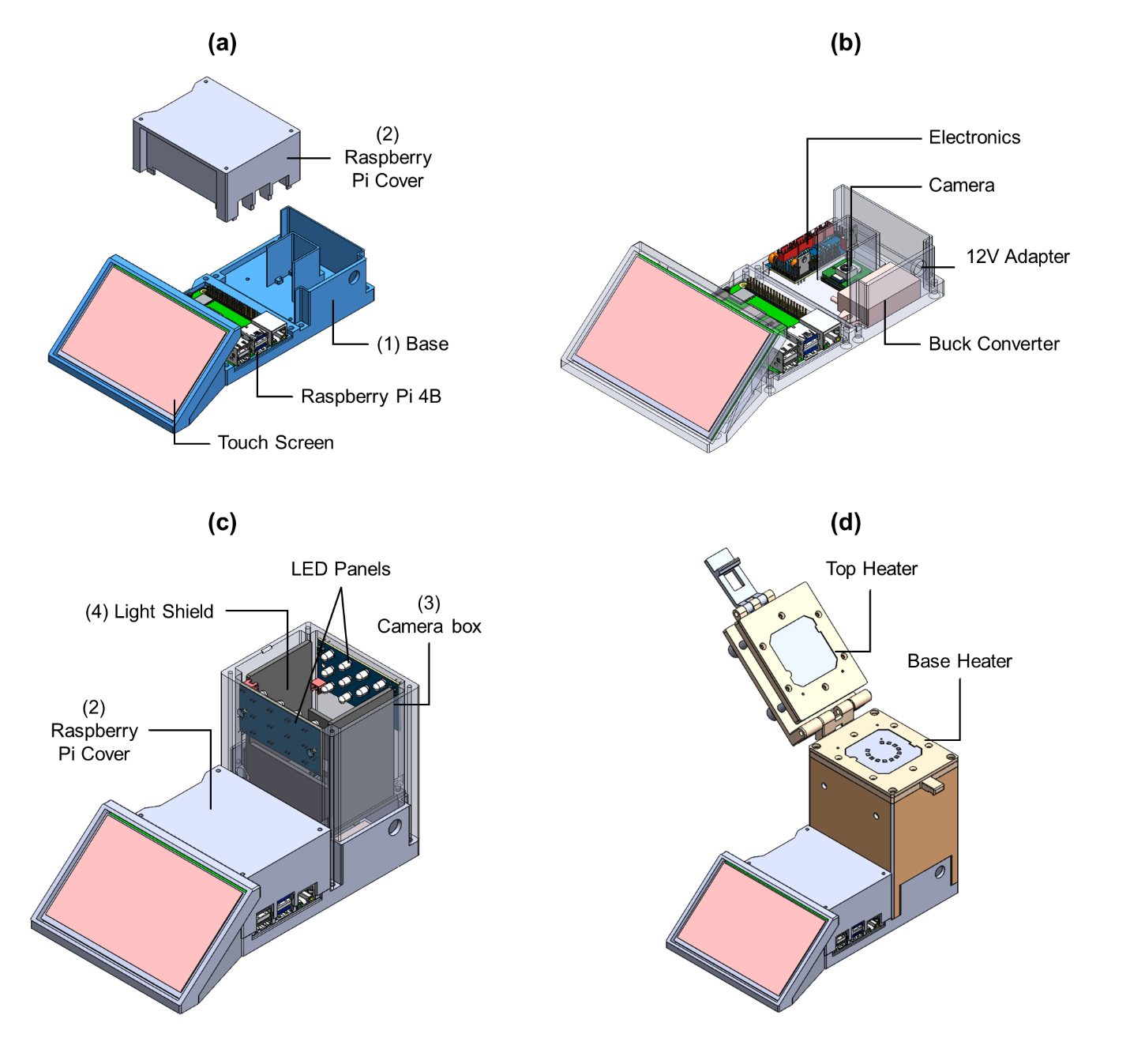

Figure S1. ThermiQuant™ VitroMini Assembly Instructions (Part I). 3D Printed design files are provided in the Supporting Information and are numbered as shown in this schematic. (a) Setup base: Place the Raspberry Pi (2) onto the base (1) and secure it with screws then place the touchscreen and connect the display cable to the Raspberry Pi. Do not install the Raspberry Pi cover (2) yet. (b) Set up camera and electronics: Place the electronics control board and camera onto the base (1), secure them with screws, and connect all required wires to the Raspberry Pi GPIO pins. After that install the 12 V adapter and connect it to the buck converter. Connect the 5 V output of the buck converter to the Raspberry Pi. Finally place the Raspberry Pi cover (2). (c) Assemble camera box: Screw the two LED panels onto the camera box (3), facing each other from opposite sides. Place the light shield (4) inside the camera box (3) and secure the camera box (3) to the base (1) using screws. (d) Install top and base heater: Refer to Figure S2 for the instruction.

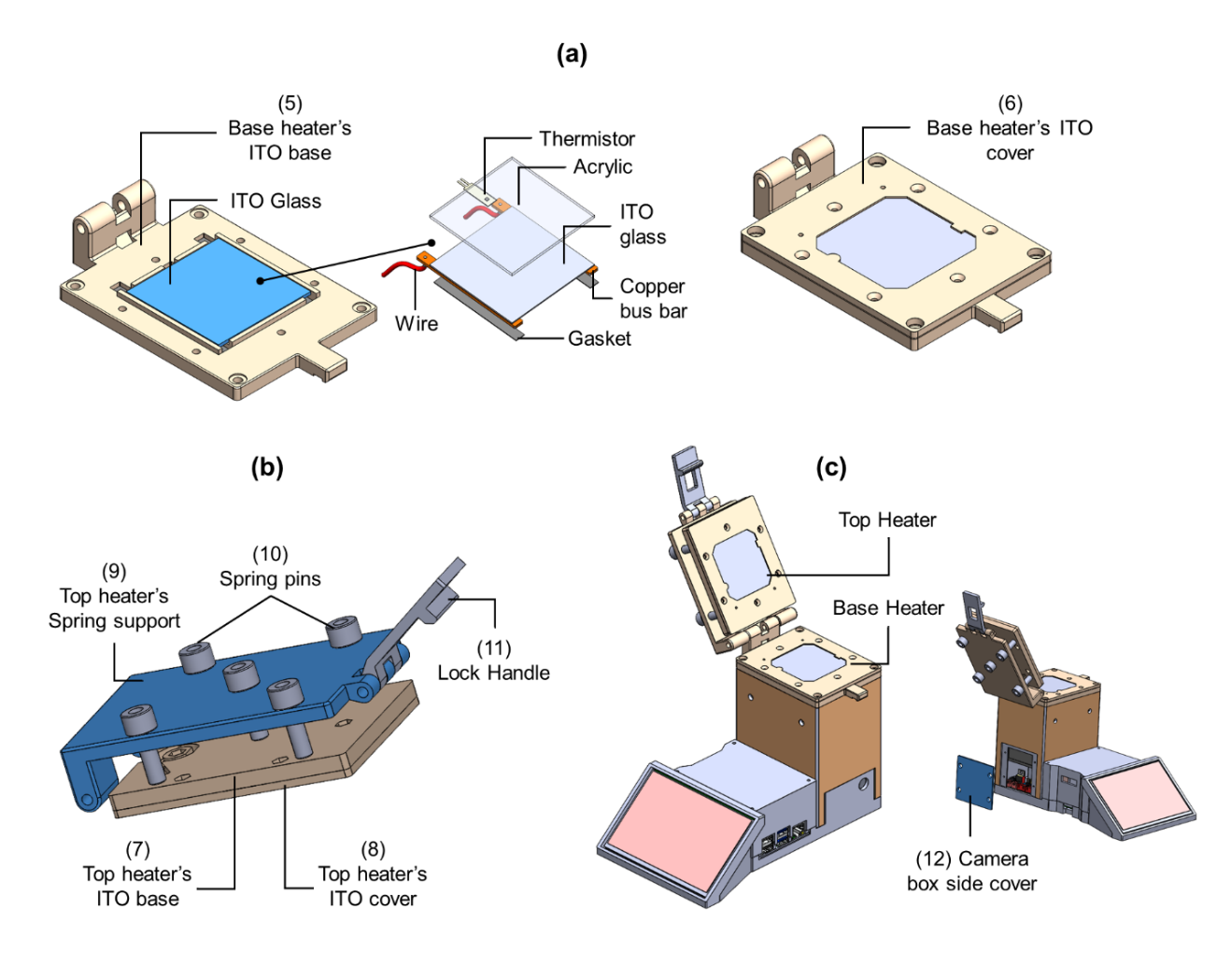

Figure S2. ThermiQuant™ VitroMini Heaters Assembly Instructions (Part II; continued from Figure S1; 3D-printed design files are available in the Supporting Information and are numbered to match this schematic). (a) Install base heater: To assemble the base heater, first place the two gaskets into the gasket grooves of the base heater’s ITO base (5). Next, place the copper bus bar with the soldered wire on top, followed by the ITO glass, ensuring that the ITO-coated side is in contact with the copper bus bar. Place the thermistor into the thermistor groove in the acrylic plate using double-sided adhesive. Then place the acrylic plate on top of the ITO glass so that the thermistor is sandwiched between them. Finally, cover the entire assembly with the base heater’s ITO cover (6). (b) Install top heater: To assemble the top heater, following the same procedure as the base heater assembly, first place the two gaskets into the gasket grooves of the top heater’s ITO base (7). Next, place the copper bus bar with the soldered wire on top, followed by the ITO glass, ensuring that the ITO-coated side is in contact with the copper bus bar. Place the thermistor into the thermistor groove in the 0.5 mm-thick thermal pad attached to the 1 mm-thick aluminum plate. Use double-sided adhesive to secure the thermal pad to the aluminum plate, then place the aluminum plate on top of the ITO glass so that the thermistor is sandwiched between them (Figure not shown). Before covering the entire assembly with the top heater’s ITO cover (8), screw the spring pins (10) to the top heater’s ITO cover (8) through the top heater’s spring support (9), ensuring that the springs are positioned between the top heater’s ITO base (7) and the top heater’s spring support (9), as shown in the schematics. Finally, install the lock handle (11) using a 4 mm-diameter bronze rod. (c) Final Assembly: Assemble the top and base heater assemblies together, then route the wires from both heaters through the hinge and into the camera box (3). Next, secure the base heater assembly to the camera box (3) using screws. Use the camera box side cover (12) to access the electronics board for maintenance or repair.

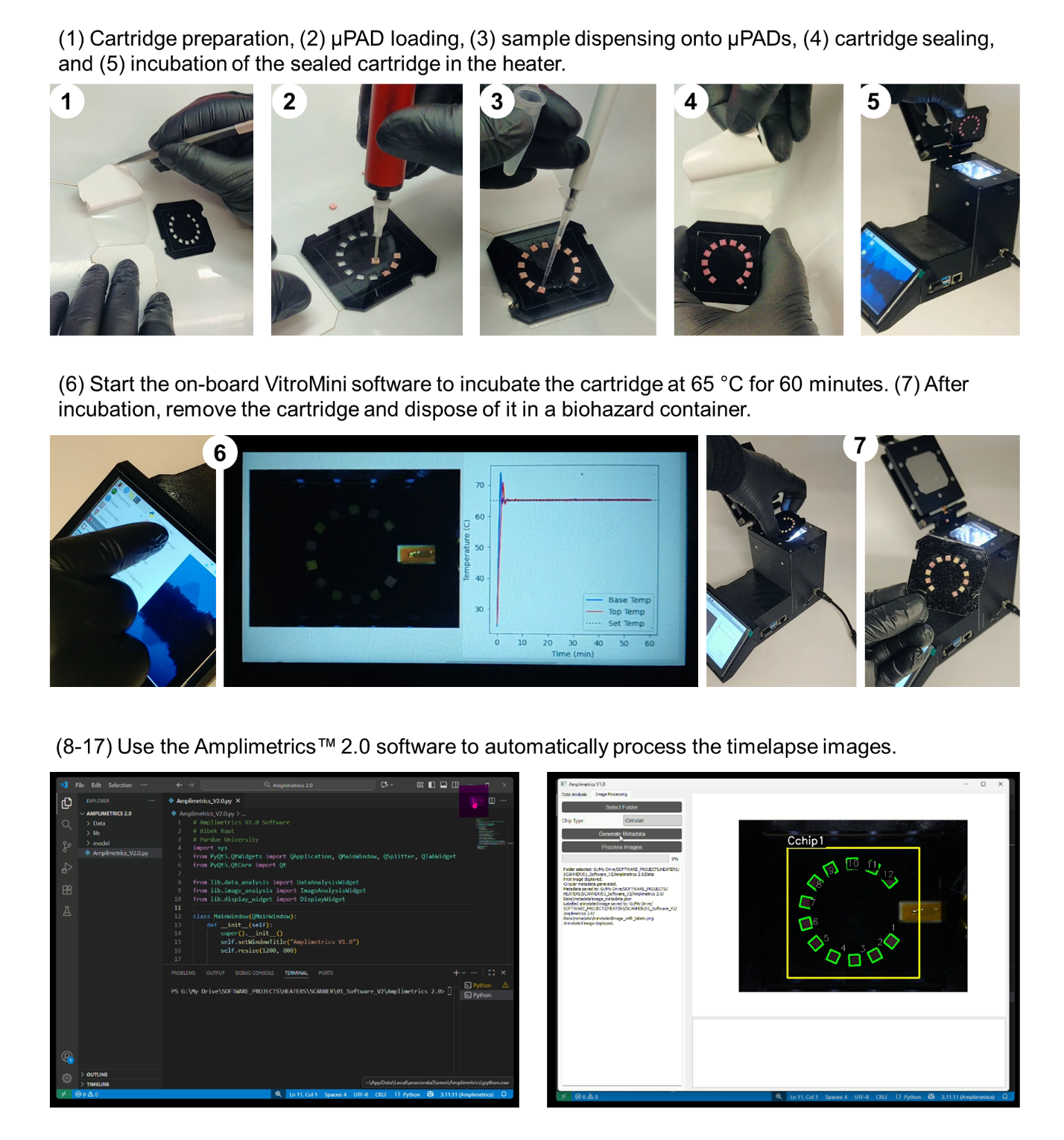

Figure S3. Step-by-step user guide for preparing the microfluidic paper-based analytical device (µPAD) cartridge for LAMP and corresponding software workflow. See Note 1.2 for the full written instructions and “Video1_Instrument_Demo_SD.mp4” in SI for video demo.

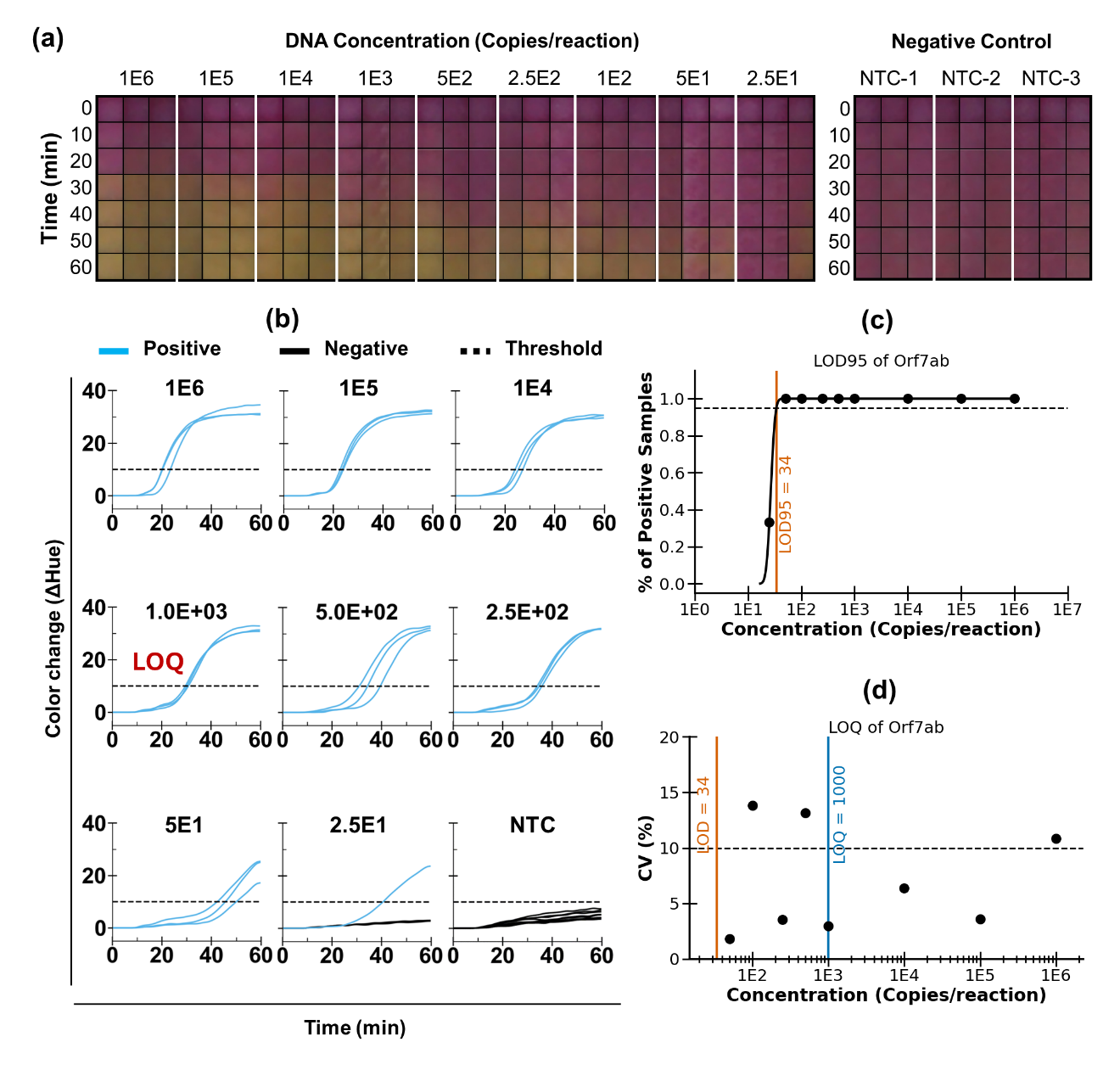

Figure S4. Analytical performance of colorimetric LAMP on µPADs using the ThermiQuant™ VitroMini. (A) Time-stamped cropped images (0–60 min, 10-min intervals) from µPADs loaded with serial dilutions of synthetic SARS-CoV-2 orf7ab DNA (10^6^–25 copies per reaction), including three technical replicates per concentration and nine no-template controls (NTCs; three each from three independent runs using the 12-µPAD cartridge). (B) Hue-versus-time trajectories generated from processed time-lapse images for each input concentration. A threshold of 10 hue units was applied to distinguish positive from negative amplification curves. (C) The limit of detection at 95% probability (LOD95) was determined using a probit fit, yielding a value of 34 copies per reaction. (D) Coefficient of variation (CV) in quantification time (Tq) as a function of concentration, with the limit of quantification (LOQ) defined as the lowest concentration with CV <10% (see definition in SI Note 1.6 under “Determination of LOD95 and LOQ”).

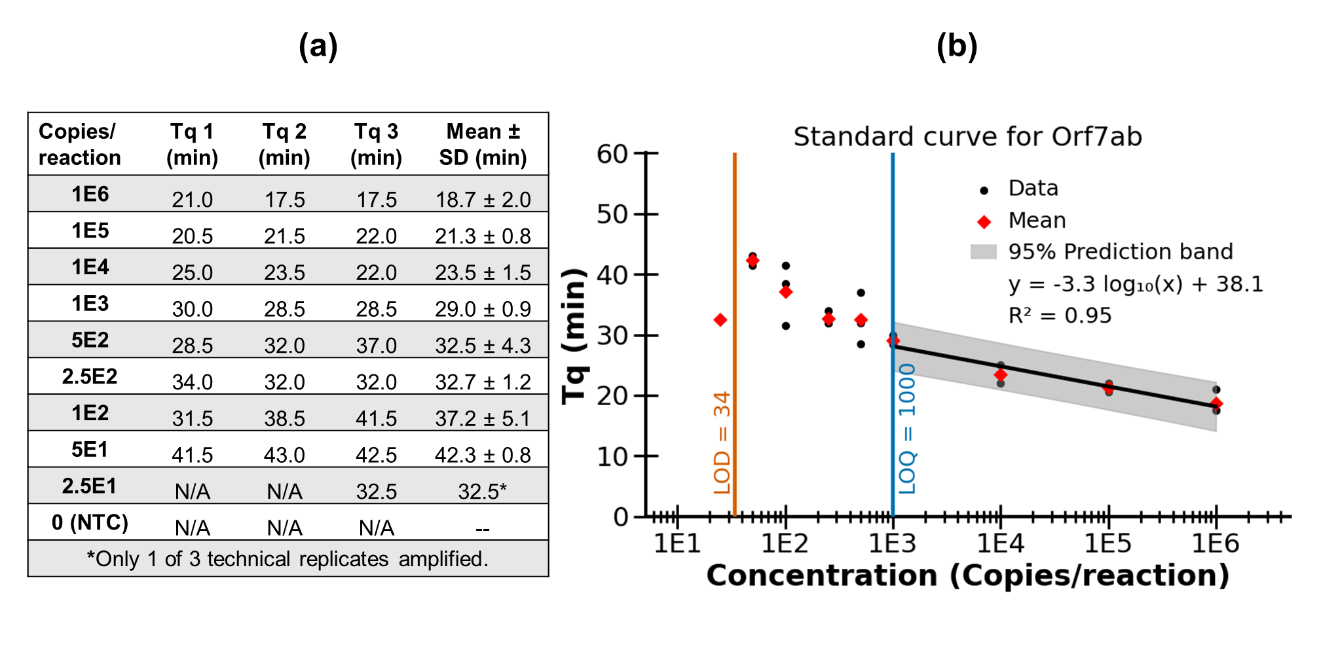

Figure S5. Standard calibration curve. (A) Summary table listing quantification time (Tq) values for each replicate and the corresponding mean ± SD across all concentrations. (D) Calibration curve plotting Tq versus input concentration, demonstrating a log-linear relationship from concentration at and above LOQ of 10^3^ copies per reaction.

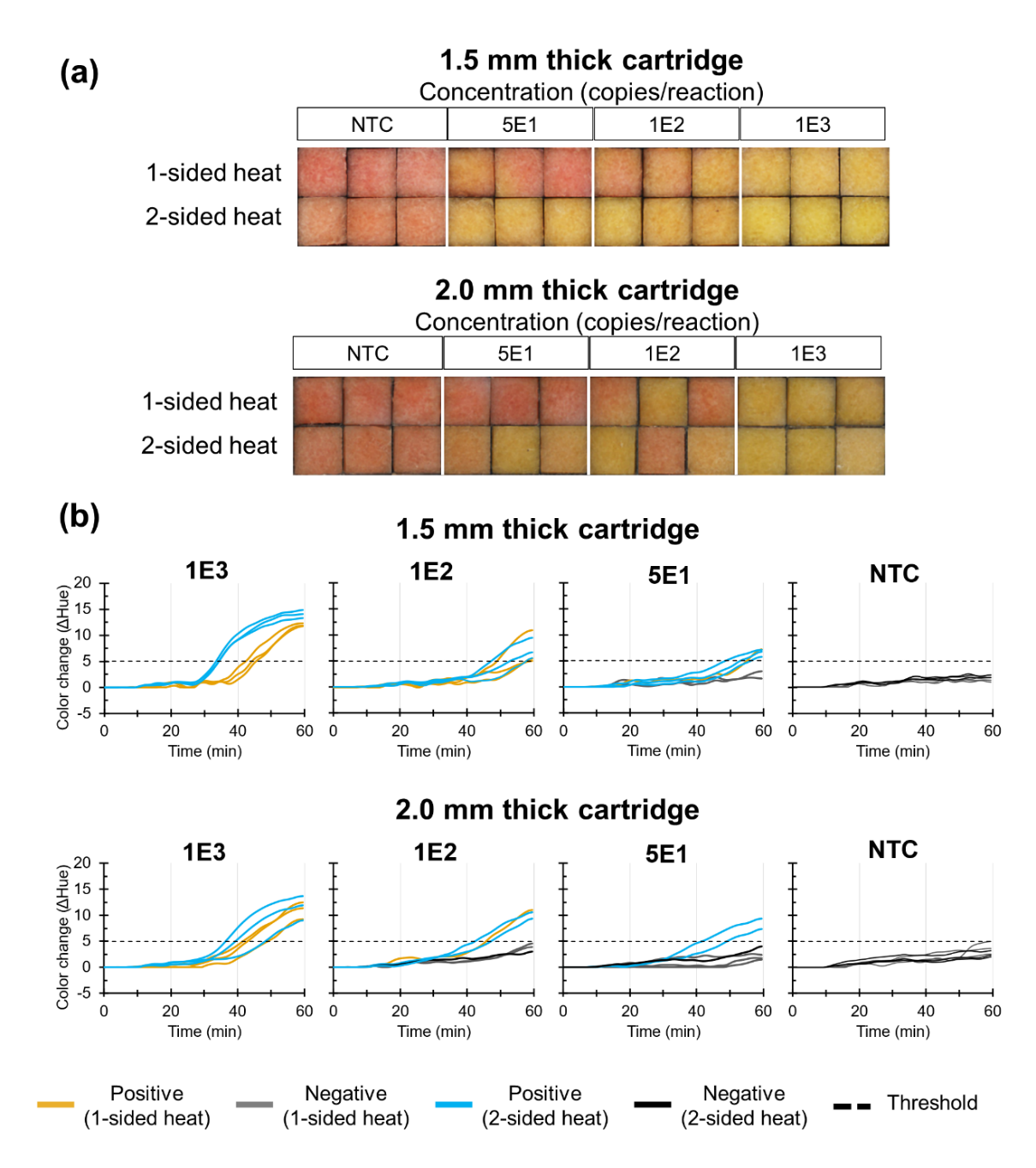

Figure S6. Effect of single-sided and dual-sided heating on limit of detection (LOD). (A) Endpoint (60 min) colorimetric images captured using a flatbed scanner (ThermiQuant™ MegaScan) for µPAD-based LAMP assays targeting orf7ab at 1E3, 1E2, 5E1 copies per reaction, along with no-template controls (NTCs). µPADs were housed in acrylic cartridges of two thicknesses (1.5 mm and 2.0 mm). Image enhanced with Microsoft PowerPoint with +40% contrast. (B) Color (net hue) versus time trajectories obtained using ThermiQuant™ VitroMini (this study) demonstrate reduced amplification efficiency under single-sided heating compared to dual-sided heating, particularly at lower input concentrations (10² and 5×10² copies per reaction), which are near the LOD95 (34 copies per reaction). A hue threshold of 5 was used to distinguish positive (sigmoidal) amplification from negative (flat/linear) responses. This threshold differed from that used in previous experiments (threshold = 10, Figure S4) due to a lower change in hue in the present µPAD batch, arising from differences in initial substrate color.

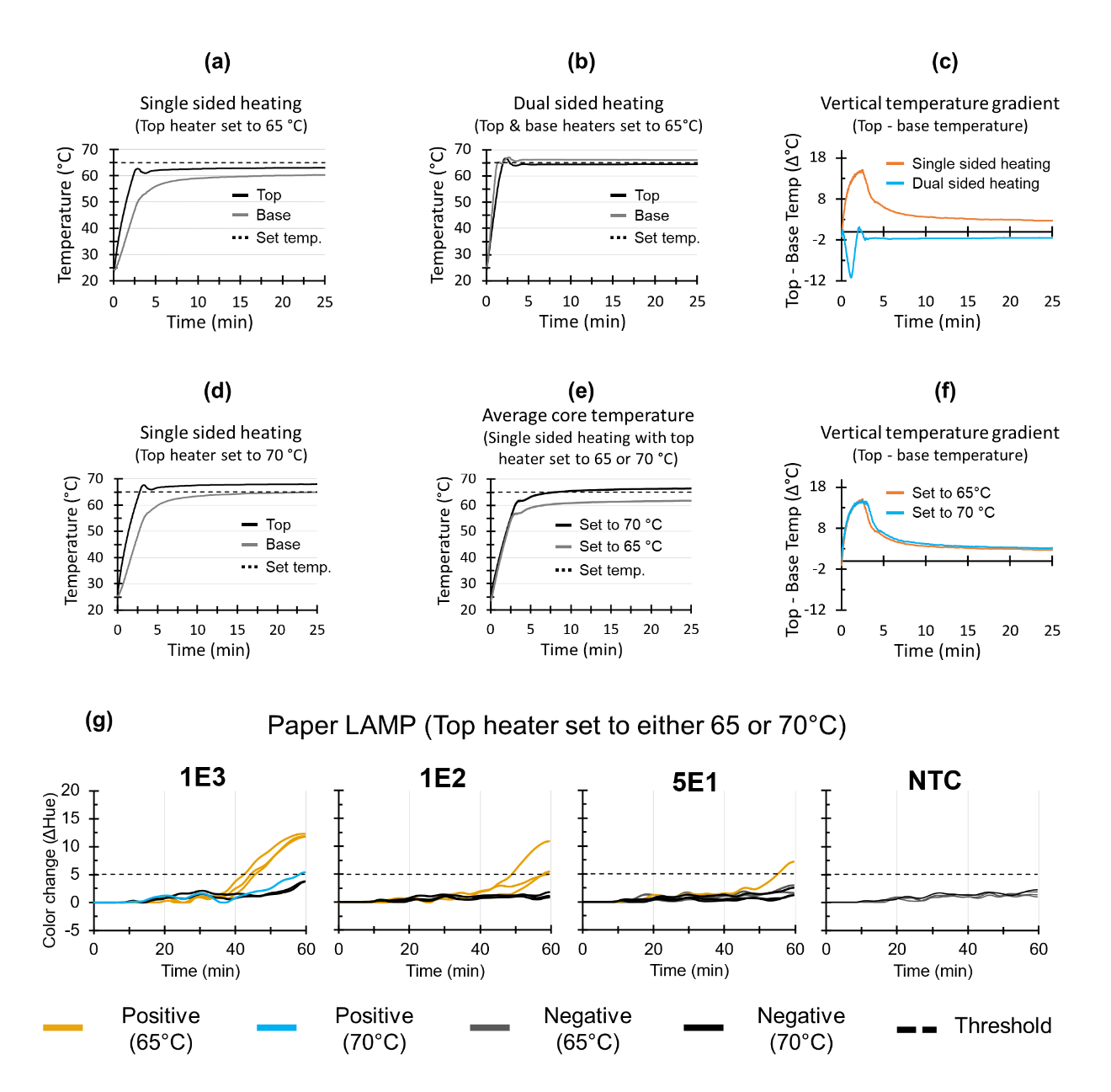

Figure S7. Vertical (through-thickness) thermal gradients and their impact on µPAD-LAMP performance under single- and dual-sided heating, including elevated temperature conditions. (A) Average temperature measured at the top surface (near the heater) and at the base (exposed to ambient conditions, ~23 °C) of a 2 mm thick acrylic cartridge (probe to probe distance 1.5 mm) under single-sided heating (top heater set to 65 °C). (B) Corresponding temperature measurements under dual-sided heating, with both top and base heaters set to 65 °C. (C) Through-thickness temperature difference (top minus base) for single- and dual-sided heating (65 °C setpoint), highlighting the larger thermal gradient in the single-sided configuration. (D) Average top and base temperatures under single-sided heating with an elevated top heater setpoint (70 °C). (E) Average core temperature within a cartridge under single-sided heating with a 65 or 70 °C top heater setpoint. (F) Through-thickness temperature difference (top minus base) under single-sided heating at top heater setpoints of 65 °C and 70 °C. (G) Hue-versus-time trajectories of µPAD-LAMP reactions under single-sided heating at 65 °C and 70 °C (elevated to achieve ~65 °C core temperature), showing reduced amplification at low DNA concentrations (1E3, 1E2, and 5E1 copies per reaction) at the higher setpoint (70 °C).

Table S1. Comparison of isothermal nucleic acid amplification tests (NAAT) instruments with optical (color/fluorescence) tracking capability published between 2020 to 2026. Acronyms: ITO- Indium tin oxide; LAMP- loop mediated isothermal amplification; PCB - printed circuit board; PTC - positive temperature coefficient; n/a - not available.

| Instruments | Heater type | Temperature precision | Color sensing | Cost (USD) | Size and weight |
| --- | --- | --- | --- | --- | --- |
| ThermiQuan™ VitroMini (this work) | Thin film (dual sided ITO) | 65 ± 0.5 °C | Camera (Hue) | $231 | 22.5 x 13 x 17.5 cm, 0.95 kg |
| Ubiquitous NAAT (UbiNAAT)^10^ | Thin film  (single sided PCB heater) | To achieve 63 °C, set to 71.5 °C | Smartphone with fluorescence optics | $1,458 | ~13 x 3 x 8 cm |
| Smartphone operated LAMP (SMART-LAMP)^11^ | Thin film (single sided Polyamide heater) | 65 ± 0.9 °C | TCS3472 color sensor | €300 (~$350) | 12.5 × 7.8 × 8.8 cm, 0.85 kg |
| Integrated Gene Box (i-Genbox)^12^ | Thin film (Single sided) | 65 ± 1-2 °C | Smartphone (Hue) | n/a | 8 cm × 5 cm × 6 cm |
| Quantitative Colorimetric LAMP (qcLAMP) device^13^ | Thin film (single sided PCB heater) | 63 ± 1-2 °C | Mini camera (RGB color differences) | n/a | 11 × 10 × 10 cm,  0.37 kg |
| Microfluidics Integrated LED-Photodiode (MILP) device ^14^ | Thin film (single sided PTC heater) | 65 °C  (precision: n/a) | Fluorescence optics | $1,763 | Test kit: 4.5 × 2.0 × 2.0 cm, device: n/a |
| Human papillomavirus (HPV) LAMP reader^15^ | Thin film (Aluminum film wrapped around reaction zone) | 65 ± 0.5 °C | Color sensor | n/a | 0.29 × 0.21 × 10.6 cm |
| LAMP Assay Reader Instrument (LARI)^16^ | Volumetric (Aluminum heat block) | 69 ± 0.4 °C | Digital light sensor | $1,500 | Bigger than 15 cm |
| ThermiQuant™ MegaScan^1^ | Volumetric (water bath) | 65 ± 0.5 °C | Scanner (Hue) | $1,573 | 50 × 31 × 25 cm,  8 kg |
| ThermiQuan™ AquaStream^17^ | Volumetric (water bath) | 65 ± 0.5 °C | Camera (Hue) | $327 | 20 × 15 × 16 cm,  5 kg |
| Field Applicable Rapid Microbial LAMP (FARM-LAMP)^18^ | Volumetric (water bath) | 65 °C (precision: n/a) | Camera (Hue) | n/a | 16.4 x 13.5 x 19.3 cm |

Table S2. Bill of materials of ThermiQuant™ VitroMini. All reported prices are for Nov 2025.

| Bill of materials for the ThermiQuant™ VitroMini | | | | | | | |
| --- | --- | --- | --- | --- | --- | --- | --- |
| S.N. | Parts Name | Manufacturer | Vendor | Vendor Parts Number | Units | Price/Pack (USD) | Total Price (USD) |
| 1 | Raspberry Pi 4B (4GB) | Raspberry Pi  Ltd | Digikey,  USA | 2648-SC0194(9)-ND | 1 | 55.00 | 55.00 |
| 2 | microSD card (64 GB) | Amazon | Amazon,  USA | ASIN:  B08TJTB8XS | 1 | 8.00 | 7.54 |
| 3 | Power Supply Adapter (12V, 5A) | Facmogu | Amazon,  USA | ASIN:  B0BYVK73BP | 1 | 12.00 | 11.99 |
| 4 | Power socket female (2.5 x 5.5mm Socket) | AJDPOI | Amazon,  USA | ASIN:  B09ZJ2G6B1 | 1 | 2.00 | 2.20 |
| 5 | 12V to 5V Buck Converter | Acridine | Amazon,  USA | ASIN:  B0BNQ5JNWZ | 1 | 6.00 | 6.38 |
| 6 | Touchscreen Monitor (5 inch, 800x480 DSI) | iPistBit | Amazon,  USA | ASIN:  B0CRR7XMCQ | 1 | 36.99 | 36.99 |
| 7 | Arducam 16 Megapixel camera | Arducam | Arducam,  USA | IMX519 | 1 | 30.00 | 29.99 |
| 8 | 2.4GHz Wireless Keyboard and Mouse for Raspberry Pi | Vilros | Amazon,  USA | ASIN:  B07LH6TZSZ | 1 | 27.00 | 26.99 |
| 9 | Duracell Coppertop AAA Batteries | Duracell | Amazon,  USA | ASIN: B004K95PBQ | 0.2 | 18.09 | 3.62 |
| 10 | 3D Printer Filament (PC, 1 kg) | OVERTURE | Amazon,  USA | ASIN:  B0BQRC3JDH | 0.5 | 29.99 | 15.00 |
| 11 | M3 Hex socket head screw, spacer & nuts kit (stainless steel) | WZHUIDA | Amazon,  USA | ASIN: ‎  B0CNJW37NJ | 0.25 | 8.99 | 2.25 |
| 12 | Raspberry Pi 4 Heatsink Kit | Pastall | Amazon,  USA | ASIN‏: ‎  B082RKKQ2D | 0.1 | 8.00 | 0.80 |
| 13 | Raspberry Pi Fan 40mm 5V DC | Easycargo | Amazon,  USA | ASIN‏: ‎  B08R1CXGCJ | 0.25 | 11.99 | 3.00 |
| 14 | 5 mm White Flat Top LED Diode (DC 3V-3.2V Volt 20mA) | CHANZON | Amazon,  USA | ASIN:  B01BUVHSSE | 0.12 | 7.00 | 0.84 |
| 15 | Indium tin Oxide (ITO) Coated Conductive Glass Slides 50×50×0.7 mm Resistance 7 ohm/sq | Qunguan Electronics | Amazon,  USA | ASIN:  ‎B0FJF54PZV | 0.25 | 35.91 | 8.98 |
| 16 | MF55 Film B3950 Thermistors with 0.15mm Thickness | Garosa | Amazon,  USA | ASIN:  B0BN8K7HPR | 0.04 | 17.00 | 0.69 |
| 17 | Transparent 1/16-Inch-Thick Acrylic Sheet 12 x 12 Inch | Outus | Amazon,  USA | ASIN:  B08C2JZKNG | 0.02 | 9.99 | 0.20 |
| 18 | Black 1/16-Inch-Thick Acrylic Sheet 12 x 12 Inch | Outus | Amazon,  USA | ASIN:  B09MRFHFTR | 0.02 | 9.00 | 0.18 |
| 19 | 6061 T651 Aluminum Sheet Metal 12" x 12" x 0.04"(1mm) Flat Plain Thin Aluminum Sheet | CeeDa | Amazon,  USA | ASIN:  B09JF1MSJS | 0.04 | 13.00 | 0.52 |
| 20 | Compression spring (10mm OD, 0.8mm Wire Size, 15mm Free Length) | uxcell | Amazon,  USA | ASIN:  B0C339Q4X5 | 1 | 6.00 | 6.49 |
| 21 | 4mm diameter Brass Rods | DYWISHKEY | Amazon,  USA | ASIN:  B0B3DM53VD | 0.05 | 10.00 | 0.50 |
| 22 | ADS1115 16 Bits 4 Channel Analog-to-Digital Converter | Teyleten Robot | Amazon,  USA | ASIN:  B0CNV9G4K1 | 0.33 | 10.00 | 3.30 |
| 23 | DS3231 Real Time Clock Module RTC | DORHEA | Amazon,  USA | ASIN:  B08X4H3NBR | 0.25 | 10.00 | 2.50 |
| 24 | IRLZ44N TO-220AB Power Mosfet N-Channel | YEGAFE | Amazon,  USA | ASIN:  B0BW8K31BF | 0.25 | 9.00 | 2.17 |
| 25 | Breakable Pin Header 2.54mm | Jabinco | Amazon,  USA | ASIN:  B0817JG3XN | 0.01 | 6.00 | 0.06 |
| 26 | 10nF 103 Ceramic Capacitors | ALLECIN | Amazon,  USA | ASIN:  B0DFWNJLXQ | 0.02 | 7.00 | 0.14 |
| 27 | 220-ohm Resistor 1/2w (0.5Watt) ±1% Tolerance | EDGELEC | Amazon,  USA | ASIN:  B07QK9ZBVZ | 0.08 | 6.00 | 0.48 |
| 28 | 100-ohm Resistor 1/2w (0.5Watt) ±1% Tolerance | EDGELEC | Amazon,  USA | ASIN:  B07QG1VL1Q | 0.02 | 6.00 | 0.12 |
| 29 | 10K ohm Resistor 1/2w (0.5Watt) ±1% Tolerance | EDGELEC | Amazon,  USA | ASIN:  B07QJB31M7 | 0.08 | 6.00 | 0.48 |
| 30 | Jst Xh 2.54 4 Pin Connector Plug Male with 200mm Wire & Female Connector | Daier | Amazon,  USA | ASIN:  B01DUC1S14 | 0.05 | 9.00 | 0.43 |
| 31 | Mini Micro JST-XH 2.54mm Pitch 2 Pin Male Connector with 10cm Silicone Wire Cables and Female Socket | ZSDZFYLLK | Amazon,  USA | ASIN:  B0D6KSMK1Q | 0.15 | 7.00 | 1.05 |
| 32 | Female to Female Breadboard Jumper Wires (10 cm length) | EDGELEC | Amazon,  USA | ASIN:  B07GD312VG | 0.1 | 7.00 | 0.70 |
| 33 | PCB (Heater control board) without shipping cost | JLCPCB | JLCPCB,  China | Custom PCB parts | 0.2 | 5.00 | 1.00 |
| 34 | PCB (LED Panel) without shipping cost | JLCPCB | JLCPCB,  China | Custom PCB parts | 0.2 | 5.00 | 1.00 |
| Total | | | | | | | **230.84** |

Table S3. LAMP primers taken from previously published work^1,3^.

| Primers | Sequence (5’-3’) | Vendor |
| --- | --- | --- |
| SC2.orf7ab.1_F3 | CGGCGTAAAACACGTCTA | IDT |
| SC2.orf7ab.1_B3 | GCTAAAAAGCACAAATAGAAG TC | IDT |
| SC2.orf7ab.1_FIP | GGAGAGTAAAGTTCTTGAACTT CCTAGTTACGTGCCAGATCAG | IDT |
| SC2.orf7ab.1_BIP | TGCGGCAATAGTGTTTATAAC ACTATGAAAGTTCAATCATTCT GTCT | IDT |
| SC2.orf7ab.1_LF | TGTCTGATGAACAGTTTAGGT GAAA | IDT |
| SC2.orf7ab.1_LB | TTGCTTCACACTCAAAAGAA | IDT |

Table S4. *Orf7ab* synthetic DNA sequence (NCBI Reference Sequence: NC_045512.2)

**Sequence (5’-3’) target sequence of *orf7ab***

ATGAAAATTATTCTTTTCTTGGCACTGATAACACTCGCTACTTGTGAGCTTTATCACTACCAAGAGTGTGTTAGAGGTACAACAGTACTTTTAAAAGAACCTT

GCTCTTCTGGAACATACGAGGGCAATTCACCATTTCATCCTCTAGCTGATAACAAATTTGCACTGACTTGCTTTAGCACTCAATTTGCTTTTGCTTGTCCTGAC

GGCGTAAAACACGTCTATCAGTTACGTGCCAGATCAGTTTCACCTAAACTGTTCATCAGACAAGAGGAAGTTCAAGAACTTTACTCTCCAATTTTTCTTATTGT

TGCGGCAATAGTGTTTATAACACTTTGCTTCACACTCAAAAGAAAGACAGAATGATTGAACTTTCATTAATTGACTTCTATTTGTGCTTTTTAGCCTTTCTGCTATTCCTTGTTTTAATTATGCTTATTATCTTTTGGTTCTCACTTGAACTGCAAGATCATAATGAAACTTGTCACGCCTAA

Table S5. Digital PCR (dPCR) primers and probes taken from previously published work^1^.

| **dPCR Primers** | **Sequence (5’-3’)** | **Vendor** |
| --- | --- | --- |
| orf7ab_PCR2_FWD | GAGGGCAATTCACCATTTCATC | Life Science Tech |
| orf7ab_PCR2_REV | AAACTGATCTGGCACGTAACT | Life Science Tech |
| Probe | /56-FAM/TT TGC TTG T/ZEN/C CTG ACG GCG TAA A/3IABkFQ/ | IDT |
