## Supplementary material for "A solid-state heater-imager for quantitative evaluation of colorimetric isothermal nucleic acid amplification on paper": Software: Amplimetrics_V2.0.py

```
# Amplimetrics V2.0 Software
# Bibek Raut
# Purdue University
import sys
from PyQt5.QtWidgets import QApplication, QMainWindow, QSplitter, QTabWidget
from PyQt5.QtCore import Qt

from lib.data_analysis import DataAnalysisWidget
from lib.image_analysis import ImageAnalysisWidget
from lib.display_widget import DisplayWidget

class MainWindow(QMainWindow):
    def __init__(self):
        super().__init__()
        self.setWindowTitle("Amplimetrics V1.0")
        self.resize(1200, 800)
        
        main_splitter = QSplitter(Qt.Horizontal)
        self.setCentralWidget(main_splitter)
        
        # Left side: Tabs for Data Analysis and Image Processing.
        self.left_tabs = QTabWidget()
        main_splitter.addWidget(self.left_tabs)
        self.left_tabs.setMinimumWidth(300)
        
        # Right side: Common display widget.
        self.display_widget = DisplayWidget()
        main_splitter.addWidget(self.display_widget)
        
        # Create the two functional widgets.
        self.data_analysis_widget = DataAnalysisWidget(self.display_widget)
        self.image_processing_widget = ImageAnalysisWidget(self.display_widget)
        
        self.left_tabs.addTab(self.data_analysis_widget, "Data Analysis")
        self.left_tabs.addTab(self.image_processing_widget, "Image Processing")

if __name__ == "__main__":
    app = QApplication(sys.argv)
    window = MainWindow()
    window.show()
    sys.exit(app.exec_())
```
