## Supplementary material for "A solid-state heater-imager for quantitative evaluation of colorimetric isothermal nucleic acid amplification on paper": Software: VitroMini.py

```
#!/usr/bin/env python3
"""
Author: Bibek Raut
Purdue Biomedical Engineering

Multiprocessing :
- The HEATER process runs a PID loop, logging time in minutes to a CSV.
- The CAMERA process captures images up to 60 minutes.
- A PyQt5 GUI in the main process shows the latest image (left) and a continuous temperature plot (right)
  in minutes, with a dotted line at set temp.
  
The code has been modified to include signal handlers and to set the GPIO pins to a safe state on startup
and shutdown.
"""

import os
import sys
import time
import math
import queue
import datetime
import threading
import multiprocessing
import csv
import signal  # For handling termination signals

# Camera imports
from picamera2 import Picamera2, Preview
from libcamera import controls

# Heater control imports
import RPi.GPIO as GPIO
import Adafruit_ADS1x15
from simple_pid import PID

# Plot / GUI imports
import matplotlib
matplotlib.use("Qt5Agg")  # ensure PyQt5 backend
import matplotlib.pyplot as plt
from matplotlib.backends.backend_qt5agg import FigureCanvasQTAgg

from PyQt5.QtCore import QTimer, Qt
from PyQt5.QtWidgets import QApplication, QWidget, QLabel, QHBoxLayout
from PyQt5.QtGui import QPixmap

# ----------------------------------------------------------------------
#                   Overall Timing Configuration
# ----------------------------------------------------------------------
TOTAL_RUN_TIME = 3600  # 60 minutes in seconds

def create_data_folders():
    """Create a timestamped folder under ~/Desktop/Raw_Data."""
    desktop_path = os.path.join(os.path.expanduser("~"), "Desktop")
    raw_data_path = os.path.join(desktop_path, "Raw_Data")
    os.makedirs(raw_data_path, exist_ok=True)

    now_str = datetime.datetime.now().strftime("%Y%m%d_%H%M%S")
    main_folder = os.path.join(raw_data_path, now_str)
    os.makedirs(main_folder, exist_ok=True)
    return main_folder

# ----------------------------------------------------------------------
#                   Camera Configuration
# ----------------------------------------------------------------------
CYCLES = 999999         # Large number so we rely on TOTAL_RUN_TIME
CYCLE_INTERVAL = 30     # Seconds per capture cycle
SHORT_INTERVAL = 2      # Delay between multiple gains in one cycle
GAIN_SETTINGS = [2.0,4.0,6.0,8.0]   # [1.0, 2.0]. Either single gain or loop through multiple gain
DISCARD_WAIT = 3
INITIAL_SETUP_DURATION = 2
EXPOSURE_TIME = 10000   # microseconds

# ----------------------------------------------------------------------
#                   Heater Configuration
# ----------------------------------------------------------------------
VCC = 3.3
R_FIXED = 10000
BETA = 3950
T0 = 298.15   # Kelvin
R0 = 10000

BASE_MOSFET_PIN = 13  # GPIO BCM 13 (physical pin 33)
TOP_MOSFET_PIN = 19   # GPIO BCM 19 (physical pin 35)
PWM_FREQUENCY = 1000

BASE_SETPOINT = 65.0
TOP_SETPOINT = 65.0

base_kp = 2.0
base_ki = 0.5
base_kd = 1.0

top_kp = 2.0
top_ki = 0.5
top_kd = 1.0

# Two safety timeouts:
TIMEOUT_SETPOINT = 90       # Must reach setpoint in 90s or we stop
TIMEOUT_OVERALL = TOTAL_RUN_TIME  # match camera's 60 min total

MAX_TEMPERATURE = 80.0
MIN_TEMPERATURE = 0.0

PLOT_UPDATE_INTERVAL = 1000  # ms for GUI timer

# ----------------------------------------------------------------------
#                   Camera Process Function
# ----------------------------------------------------------------------
def camera_process_func(
    main_folder,
    gain_settings,
    cycles,
    cycle_interval,
    short_interval,
    discard_wait,
    initial_setup_duration,
    exposure_time,
    total_run_time
):
    """
    Capture images in cycles until total_run_time is reached or cycles are done.
    Save them in subfolders for each gain, plus 'latest.jpg' for the GUI.
    """
    # Register signal handlers for graceful shutdown.
    # When a termination signal is received, raise KeyboardInterrupt.
    def signal_handler(signum, frame):
        raise KeyboardInterrupt
    signal.signal(signal.SIGTERM, signal_handler)
    signal.signal(signal.SIGINT, signal_handler)

    # Create subfolders for each gain
    subfolders = {}
    for g in gain_settings:
        sf = os.path.join(main_folder, f"Gain_{int(g)}")
        os.makedirs(sf, exist_ok=True)
        subfolders[g] = sf

    # Initialize camera
    picam2 = Picamera2()
    preview_config = picam2.create_preview_configuration(
        controls={"AfMode": 1, "AfTrigger": 0},
        main={"size": (640, 480)},
        buffer_count=1
    )
    picam2.configure(preview_config)

    picam2.start_preview(Preview.NULL)
    picam2.start()

    start_time = time.time()

    # Discard an initial frame
    time.sleep(discard_wait)
    _ = picam2.capture_array()  # discard
    time.sleep(initial_setup_duration)

    # Optional autofocus
    picam2.set_controls({"AfTrigger": 1})
    lens_position = None
    for _ in range(3):
        time.sleep(1)
        md = picam2.capture_metadata()
        lens_position = md.get("LensPosition")
        if lens_position is not None:
            break
    if lens_position is not None:
        picam2.set_controls({"AfMode": 0, "LensPosition": lens_position})

    frame_counts = {g: 1 for g in gain_settings}

    try:
        for cycle_idx in range(cycles):
            elapsed = time.time() - start_time
            if elapsed >= total_run_time:
                break

            cycle_start = time.time()

            for g in gain_settings:
                elapsed = time.time() - start_time
                if elapsed >= total_run_time:
                    break

                # Set camera controls
                picam2.set_controls({"ExposureTime": exposure_time, "AnalogueGain": g})
                time.sleep(0.3)

                # Discard warm-up frame
                _ = picam2.capture_array()
                time.sleep(0.2)

                # Capture the official image
                fname = f"Gain{int(g)}_Exp{exposure_time}_Frame{frame_counts[g]}.jpg"
                save_path = os.path.join(subfolders[g], fname)
                picam2.capture_file(save_path)
                frame_counts[g] += 1

                # Copy to "latest.jpg" for GUI display
                import shutil
                latest_path = os.path.join(main_folder, "latest.jpg")
                try:
                    shutil.copyfile(save_path, latest_path)
                except Exception as e:
                    print(f"[Camera] Error copying latest image: {e}")

                time.sleep(short_interval)

            # Sleep until the next cycle
            cycle_elapsed = time.time() - cycle_start
            remain = cycle_interval - cycle_elapsed
            if remain > 0:
                time.sleep(remain)

            if (time.time() - start_time) >= total_run_time:
                break
    except KeyboardInterrupt:
        print("[Camera] Received interrupt signal. Shutting down camera process.")
    finally:
        picam2.stop()
        picam2.close()

# ----------------------------------------------------------------------
#                   Heater Process Function
# ----------------------------------------------------------------------
def read_thermistor_temp(adc, ch1, ch2):
    """
    Reads temperature from a thermistor (Wheatstone bridge) on 2 ADC channels.
    Returns temperature in Celsius (float) or None if an error occurs.
    """
    try:
        raw1 = adc.read_adc(ch1, gain=1)
        raw2 = adc.read_adc(ch2, gain=1)

        v1 = raw1 * (4.096 / 32768.0)
        v2 = raw2 * (4.096 / 32768.0)

        diff = abs(v1 - v2)
        if diff >= VCC:
            return None

        num = R_FIXED * v2
        den = (2.0 * v1) - v2
        if den == 0:
            return None

        R_therm = num / den
        if R_therm <= 0:
            return None

        temp_k = 1.0 / ((1.0/T0) + (1.0/BETA)*math.log(R_therm/R0))
        return temp_k - 273.15
    except Exception as e:
        print(f"[Heater] Error reading thermistor: {e}")
        return None
def heater_process_func(main_folder, csv_queue, total_run_time):
    """
    Runs the heater PID logic until total_run_time or a safety condition occurs.
    Logs time (in minutes), temperatures, and PID outputs to a CSV file.
    Sends (time, base_temp, top_temp) tuples to the GUI via a multiprocessing.Queue.
    """
    # Register signal handlers for graceful shutdown.
    def signal_handler(signum, frame):
        raise KeyboardInterrupt
    signal.signal(signal.SIGTERM, signal_handler)
    signal.signal(signal.SIGINT, signal_handler)

    csv_path = os.path.join(main_folder, "temperature_data.csv")
    with open(csv_path, "w", newline="") as f:
        writer = csv.writer(f)
        writer.writerow(["Time_min", "BaseTemp_C", "TopTemp_C", "BasePWM", "TopPWM"])

    # Setup GPIO with initial safe state (LOW)
    GPIO.setmode(GPIO.BCM)
    GPIO.setup(BASE_MOSFET_PIN, GPIO.OUT, initial=GPIO.LOW)
    GPIO.setup(TOP_MOSFET_PIN, GPIO.OUT, initial=GPIO.LOW)

    base_pwm = GPIO.PWM(BASE_MOSFET_PIN, PWM_FREQUENCY)
    top_pwm = GPIO.PWM(TOP_MOSFET_PIN, PWM_FREQUENCY)
    base_pwm.start(0)
    top_pwm.start(0)

    # Setup ADC
    adc = Adafruit_ADS1x15.ADS1115(address=0x48, busnum=1)

    # Initialize PID controllers
    pid_base = PID(base_kp, base_ki, base_kd, setpoint=BASE_SETPOINT)
    pid_base.output_limits = (0, 100)
    pid_base.sample_time = 1.0

    pid_top = PID(top_kp, top_ki, top_kd, setpoint=TOP_SETPOINT)
    pid_top.output_limits = (0, 100)
    pid_top.sample_time = 1.0

    start_time = time.time()
    base_reached = False
    top_reached = False

    try:
        while True:
            elapsed_s = time.time() - start_time
            if elapsed_s >= total_run_time:
                break

            base_temp = read_thermistor_temp(adc, 0, 1)
            top_temp  = read_thermistor_temp(adc, 2, 3)

            if base_temp is not None and top_temp is not None:
                # Check if setpoint has been reached
                if not base_reached and base_temp >= BASE_SETPOINT:
                    base_reached = True
                if not top_reached and top_temp >= TOP_SETPOINT:
                    top_reached = True

                # Safety checks
                if elapsed_s > TIMEOUT_SETPOINT:
                    if not base_reached or not top_reached:
                        print("[Heater] Error: setpoint not reached in time.")
                        break
                if elapsed_s > TIMEOUT_OVERALL:
                    print("[Heater] Overall internal timeout reached.")
                    break
                if base_temp > MAX_TEMPERATURE or top_temp > MAX_TEMPERATURE:
                    print("[Heater] ALERT: Over max temp!")
                    break
                if base_temp < MIN_TEMPERATURE or top_temp < MIN_TEMPERATURE:
                    print("[Heater] ALERT: Below min temp!")
                    break

                # Calculate new duty cycles using the PID controllers
                base_duty = pid_base(base_temp)
                top_duty  = pid_top(top_temp)

                base_pwm.ChangeDutyCycle(base_duty)
                top_pwm.ChangeDutyCycle(top_duty)

                # Convert elapsed time to minutes (with full float for GUI and CSV rounded to 3 decimals)
                time_min_float = elapsed_s / 60.0
                time_min_str   = f"{time_min_float:.3f}"

                # Append the new data to the CSV log
                with open(csv_path, "a", newline="") as fcsv:
                    writer = csv.writer(fcsv)
                    writer.writerow([
                        time_min_str,
                        f"{base_temp:.2f}",
                        f"{top_temp:.2f}",
                        f"{base_duty:.1f}",
                        f"{top_duty:.1f}"
                    ])

                # Send data to the GUI
                csv_queue.put((time_min_float, base_temp, top_temp))
            else:
                print("[Heater] Could not read sensors.")

            time.sleep(1.0)
    except KeyboardInterrupt:
        print("[Heater] Received interrupt signal. Shutting down heater process.")
    finally:
        # Safely shut down the heater by setting duty cycles to 0
        base_pwm.ChangeDutyCycle(0)
        top_pwm.ChangeDutyCycle(0)
        base_pwm.stop()
        top_pwm.stop()
        GPIO.cleanup()
        print("[Heater] Cleanup done.")

# ----------------------------------------------------------------------
#                   PyQt5 GUI in Main Process
# ----------------------------------------------------------------------
class MatplotCanvas(FigureCanvasQTAgg):
    def __init__(self, parent=None):
        import matplotlib.figure as mplfig
        fig = mplfig.Figure()
        self.ax = fig.add_subplot(111)
        super().__init__(fig)
        self.setParent(parent)

class MainWindow(QWidget):
    def __init__(self, main_folder, csv_queue):
        super().__init__()
        self.setWindowTitle("Heater + Camera Monitor (60 min)")

        self.main_folder = main_folder
        self.csv_queue   = csv_queue

        layout = QHBoxLayout()

        self.image_label = QLabel("No image yet.")
        self.image_label.setAlignment(Qt.AlignCenter)
        layout.addWidget(self.image_label, stretch=1)

        self.canvas = MatplotCanvas(self)
        layout.addWidget(self.canvas, stretch=1)

        self.setLayout(layout)

        # Data storage for plotting
        self.time_data = []
        self.base_data = []
        self.top_data  = []

        self.update_timer = QTimer(self)
        self.update_timer.timeout.connect(self.update_gui)
        self.update_timer.start(PLOT_UPDATE_INTERVAL)

    def update_gui(self):
        # 1) Update the image from the latest capture.
        latest_img = os.path.join(self.main_folder, "latest.jpg")
        if os.path.exists(latest_img):
            pix = QPixmap(latest_img)
            if not pix.isNull():
                scaled_pix = pix.scaled(
                    self.image_label.width(),
                    self.image_label.height(),
                    Qt.KeepAspectRatio,
                    Qt.SmoothTransformation
                )
                self.image_label.setPixmap(scaled_pix)

        # 2) Drain the CSV queue for new temperature data.
        while True:
            try:
                item = self.csv_queue.get_nowait()
                # Each item is a tuple: (time_min_float, base_temp, top_temp)
                self.time_data.append(item[0])
                self.base_data.append(item[1])
                self.top_data.append(item[2])
            except queue.Empty:
                break

        # 3) Update the plot.
        ax = self.canvas.ax
        ax.clear()
        ax.set_xlabel("Time (min)")
        ax.set_ylabel("Temperature (C)")
        ax.grid(False)
        if self.time_data:
            ax.plot(self.time_data, self.base_data, 'b-', label="Base Temp")
            ax.plot(self.time_data, self.top_data, 'r-', label="Top Temp")

        # Dotted line showing the set temperature
        ax.axhline(y=BASE_SETPOINT, color='black', linestyle=':', label="Set Temp")
        ax.legend()
        ax.relim()
        ax.autoscale_view()
        self.canvas.draw()

# ----------------------------------------------------------------------
#                   Main Function
# ----------------------------------------------------------------------
def main():
    main_folder = create_data_folders()
    csv_queue   = multiprocessing.Queue()

    # Start the camera process (non-daemon for proper cleanup)
    camera_proc = multiprocessing.Process(
        target=camera_process_func,
        args=(
            main_folder,
            GAIN_SETTINGS,
            CYCLES,
            CYCLE_INTERVAL,
            SHORT_INTERVAL,
            DISCARD_WAIT,
            INITIAL_SETUP_DURATION,
            EXPOSURE_TIME,
            TOTAL_RUN_TIME
        )
    )

    # Start the heater process (non-daemon for proper cleanup)
    heater_proc = multiprocessing.Process(
        target=heater_process_func,
        args=(main_folder, csv_queue, TOTAL_RUN_TIME)
    )

    camera_proc.start()
    heater_proc.start()

    # Launch the Qt GUI
    app = QApplication(sys.argv)
    gui = MainWindow(main_folder, csv_queue)
    gui.resize(1200, 600)
    gui.show()

    # When the GUI closes, terminate the child processes gracefully.
    def on_exit():
        print("[Main] Shutting down child processes.")
        if camera_proc.is_alive():
            camera_proc.terminate()
        if heater_proc.is_alive():
            heater_proc.terminate()
        camera_proc.join(timeout=5)
        heater_proc.join(timeout=5)
    app.aboutToQuit.connect(on_exit)

    app.exec_()

    print("[Main] End of 60-minute run or GUI closed.")

if __name__ == "__main__":
    main()
```
