## Supplementary figures and images for "A solid-state heater-imager for quantitative evaluation of colorimetric isothermal nucleic acid amplification on paper"

### Gain8_Exp1000_Frame9.jpg

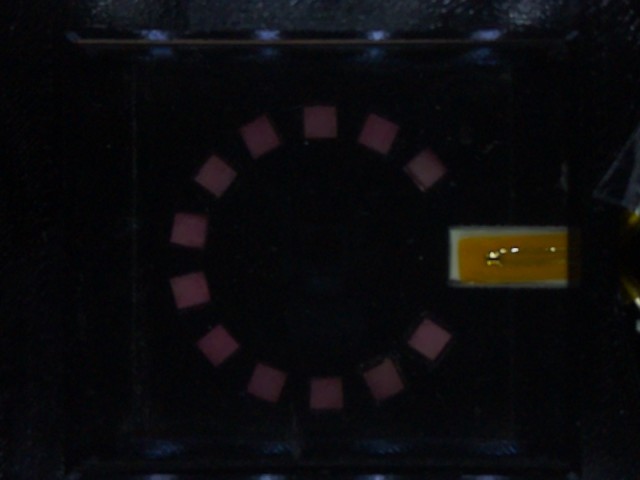

### Gain8_Exp1000_Frame81.jpg

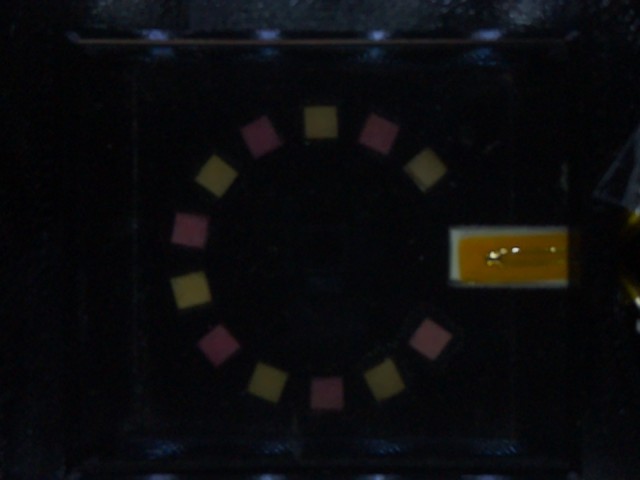

### Gain8_Exp1000_Frame82.jpg

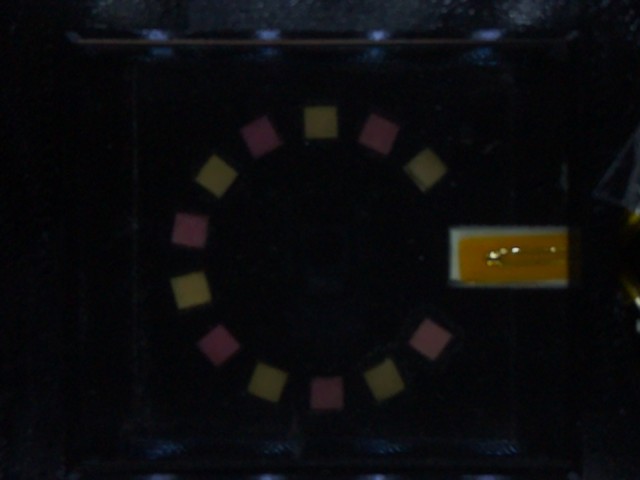

### Gain8_Exp1000_Frame83.jpg

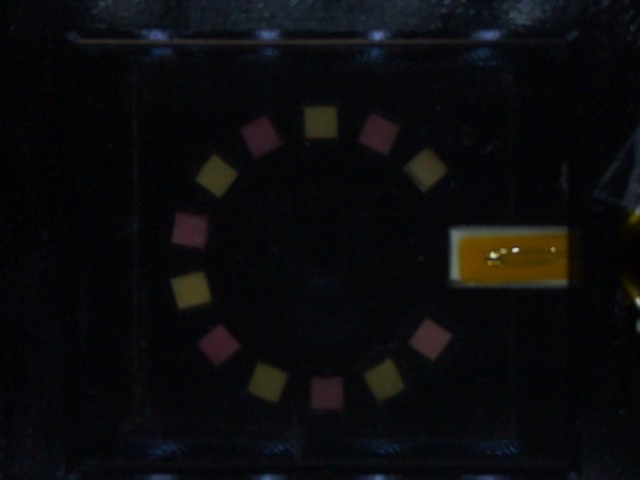

### Gain8_Exp1000_Frame84.jpg

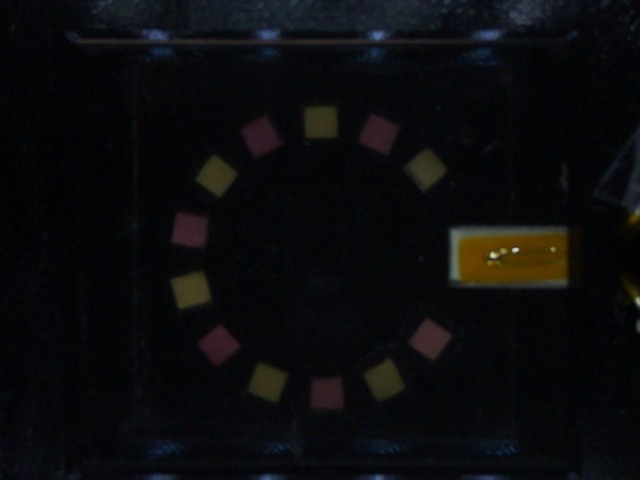

### Gain8_Exp1000_Frame85.jpg

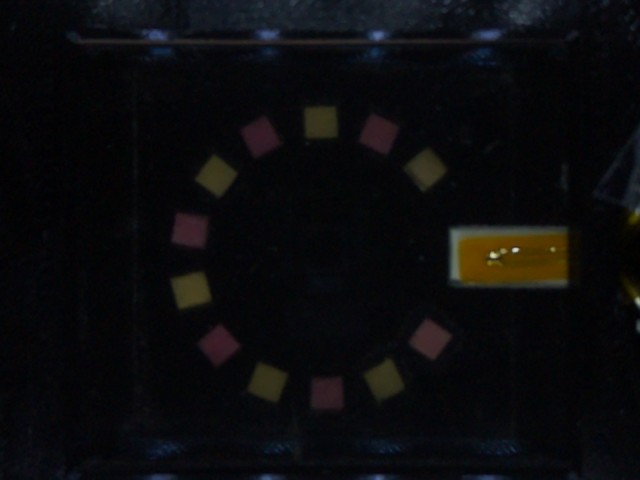

### Gain8_Exp1000_Frame86.jpg

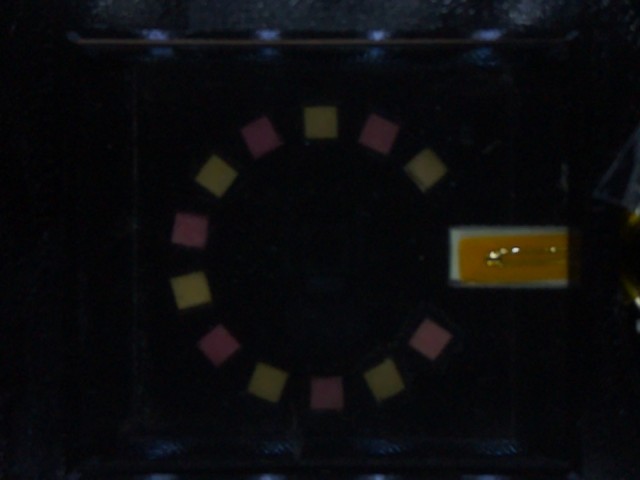

### Gain8_Exp1000_Frame87.jpg

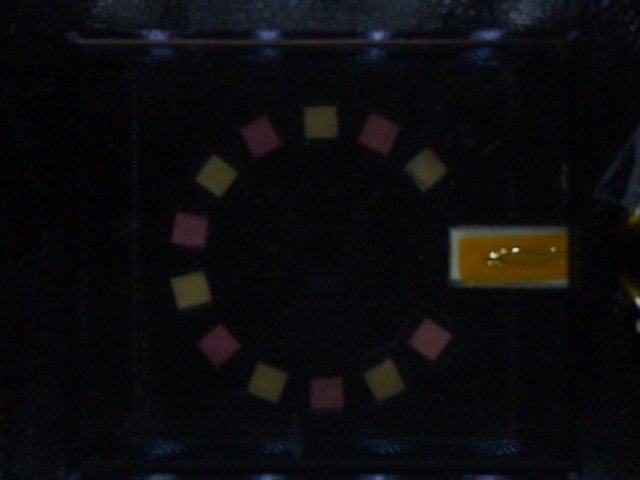

### Gain8_Exp1000_Frame88.jpg

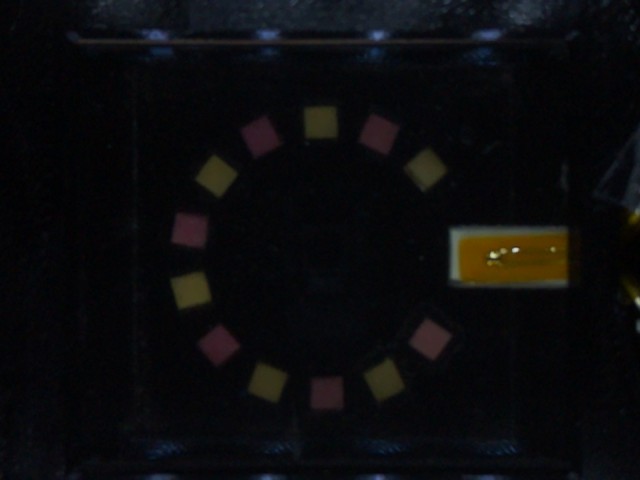

### Gain8_Exp1000_Frame89.jpg

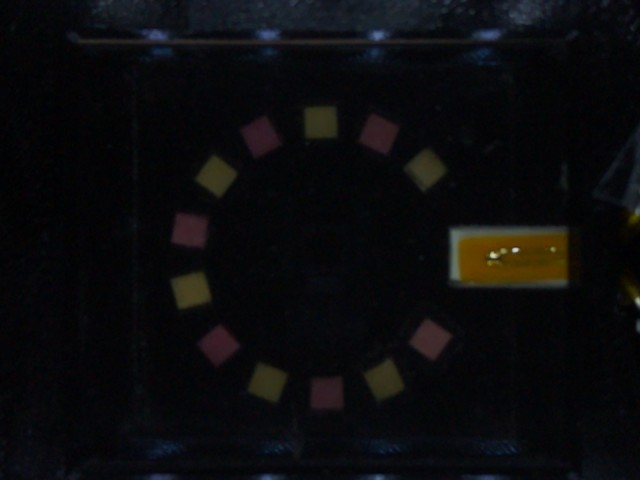

### Gain8_Exp1000_Frame90.jpg

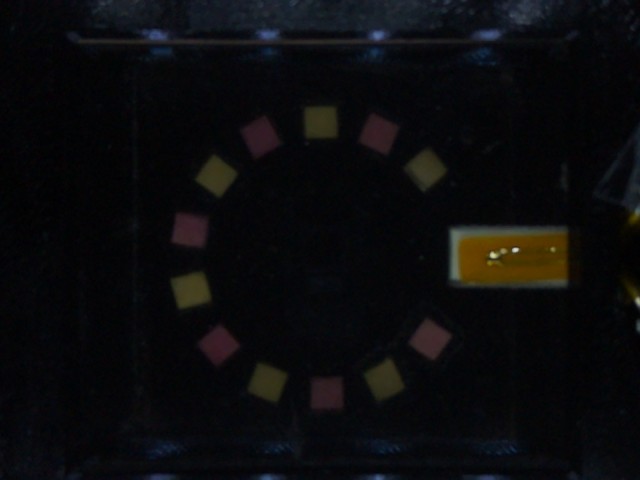

### Gain8_Exp1000_Frame91.jpg

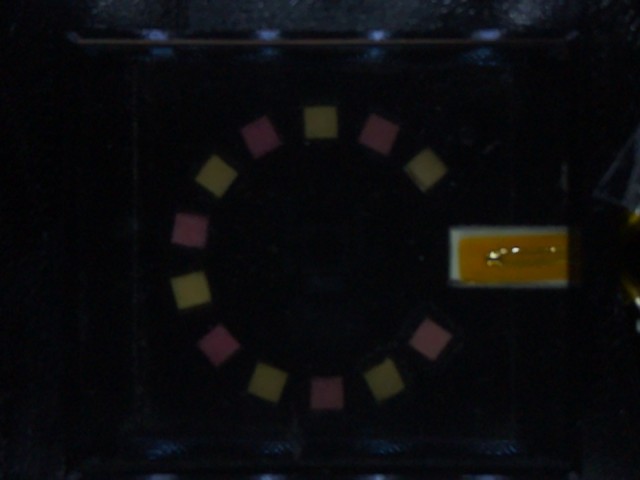

### Gain8_Exp1000_Frame92.jpg

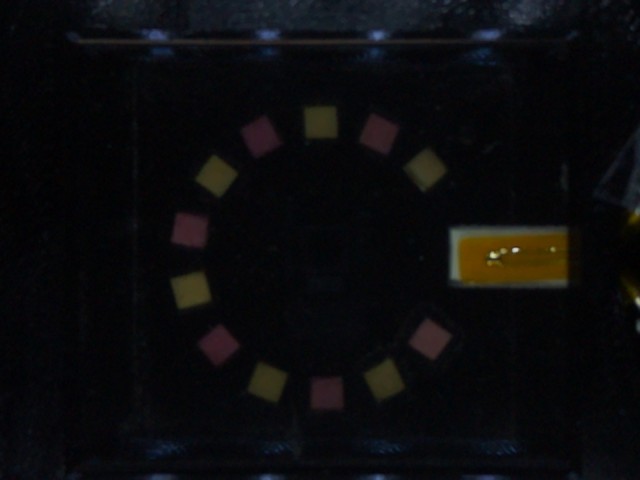

### Gain8_Exp1000_Frame93.jpg

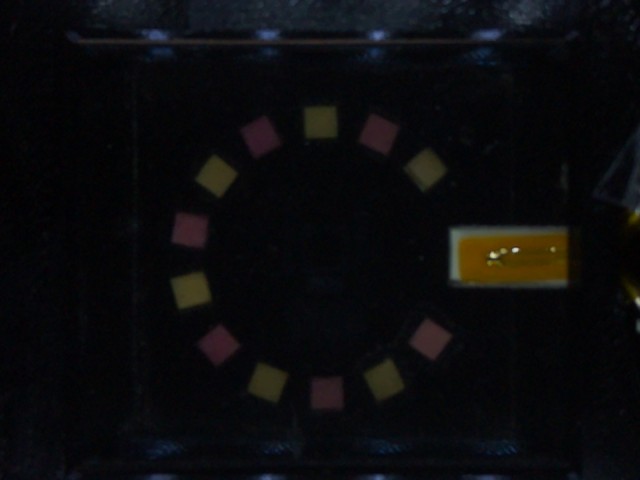

### Gain8_Exp1000_Frame94.jpg

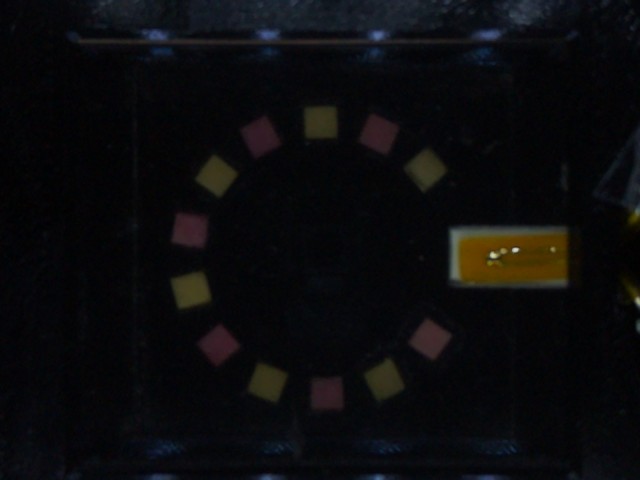

### Gain8_Exp1000_Frame95.jpg

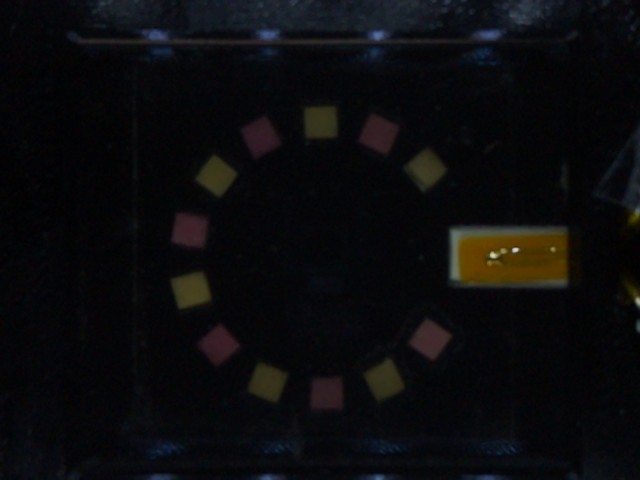

### Gain8_Exp1000_Frame96.jpg

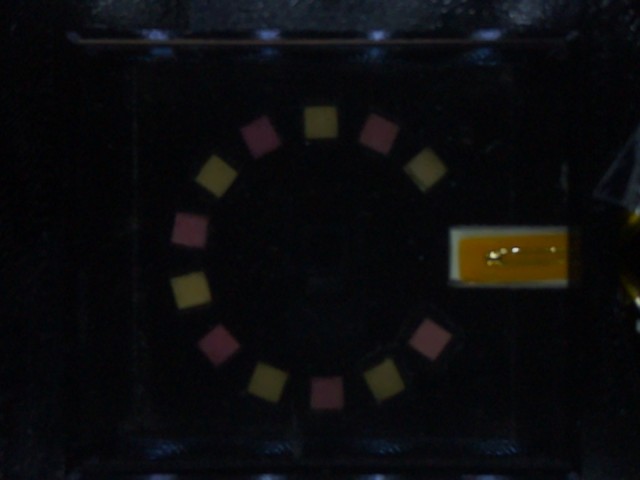

### Gain8_Exp1000_Frame97.jpg

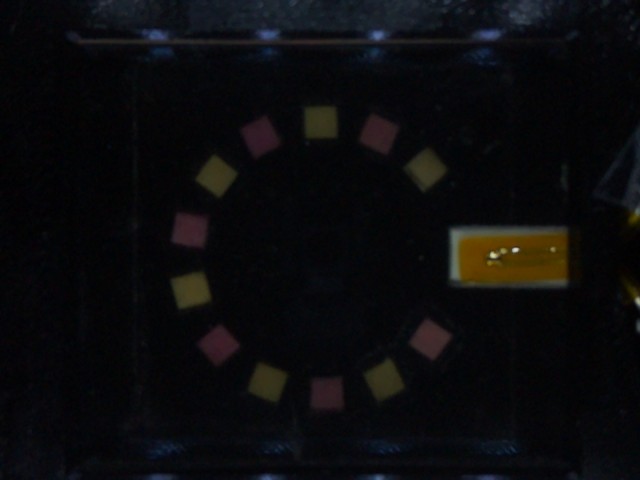

### Gain8_Exp1000_Frame98.jpg

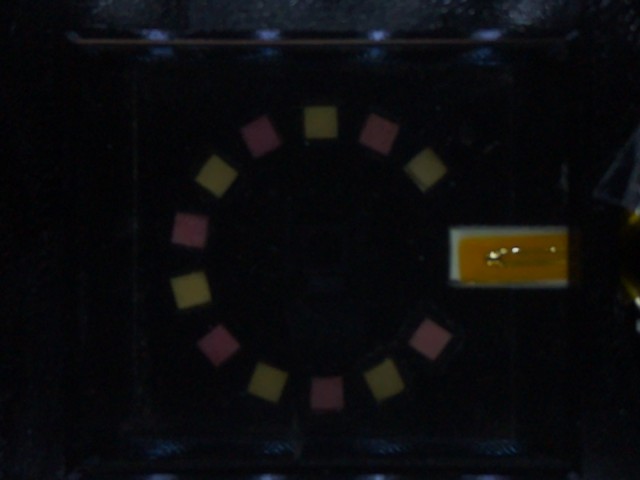

### Gain8_Exp1000_Frame99.jpg

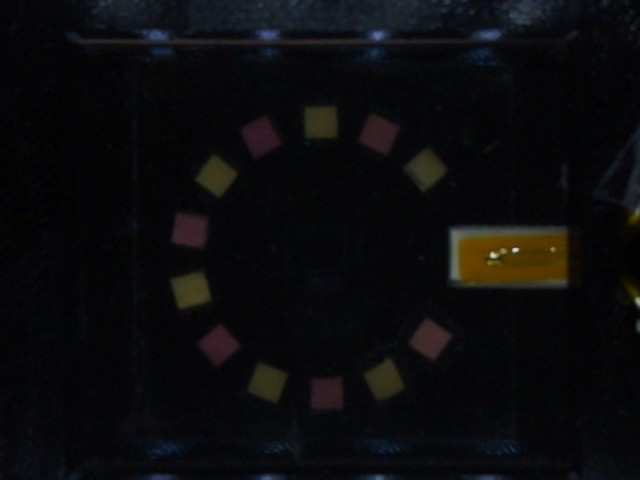
