## Supplementary figures and images for "A solid-state heater-imager for quantitative evaluation of colorimetric isothermal nucleic acid amplification on paper"

### Gain8_Exp1000_Frame1.jpg

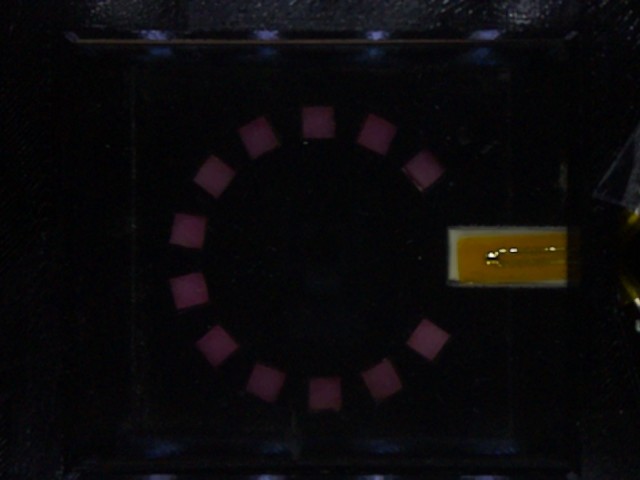

### Gain8_Exp1000_Frame2.jpg

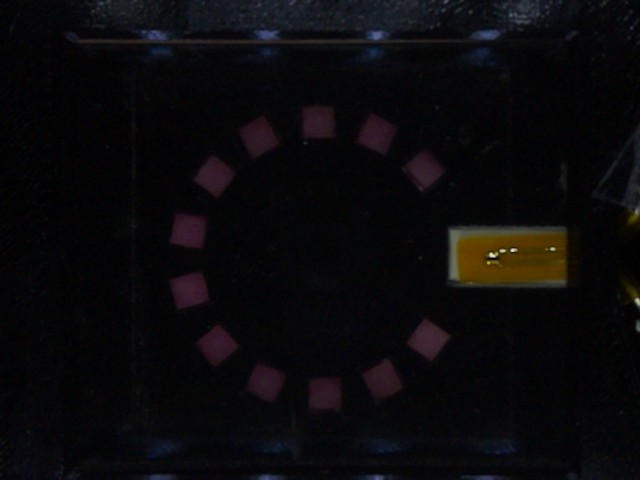

### Gain8_Exp1000_Frame3.jpg

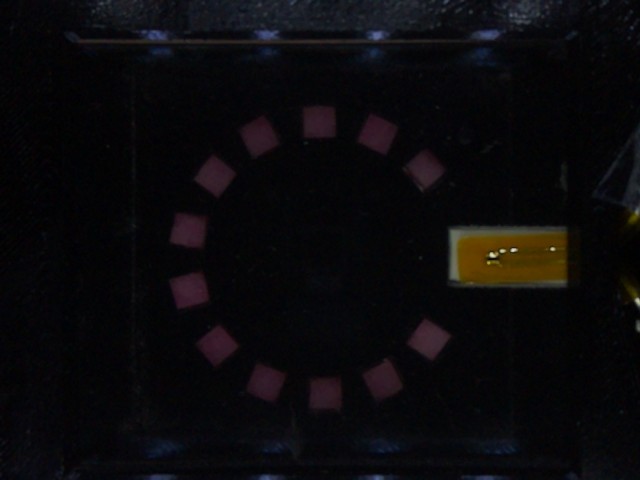

### Gain8_Exp1000_Frame4.jpg

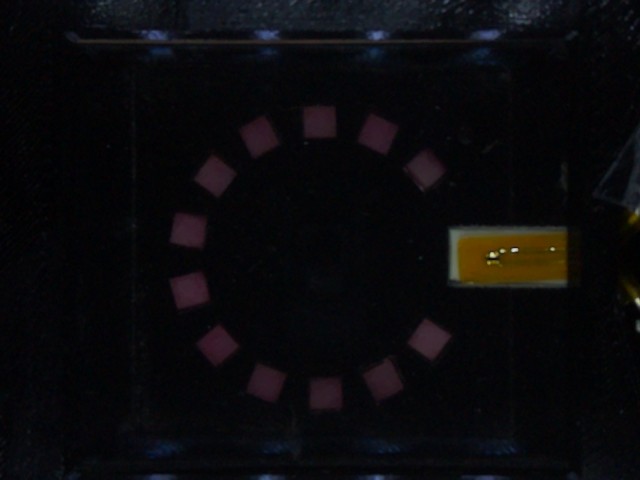

### Gain8_Exp1000_Frame5.jpg

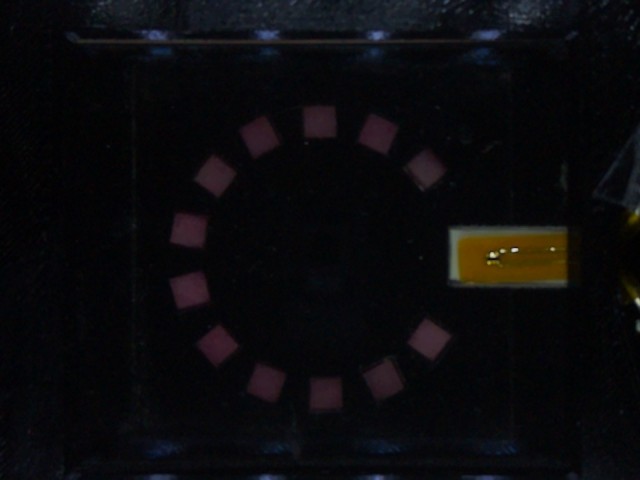

### Gain8_Exp1000_Frame6.jpg

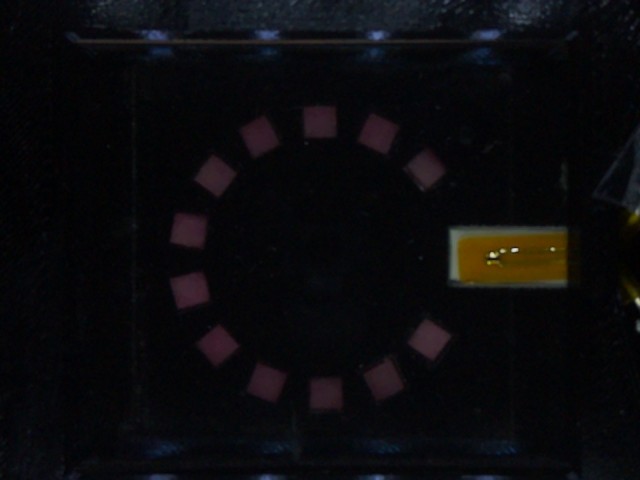

### Gain8_Exp1000_Frame7.jpg

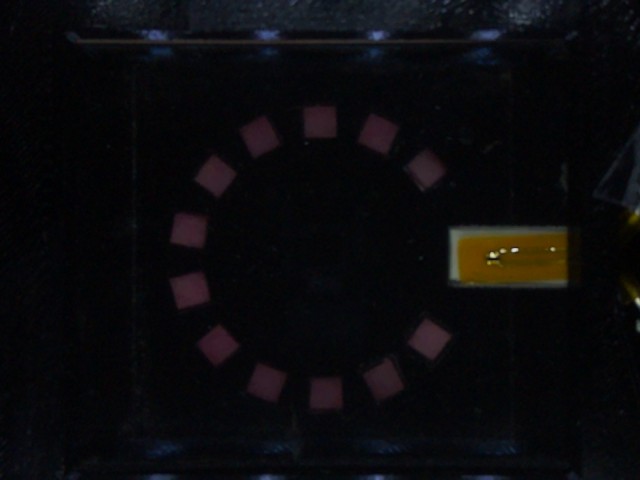

### Gain8_Exp1000_Frame8.jpg

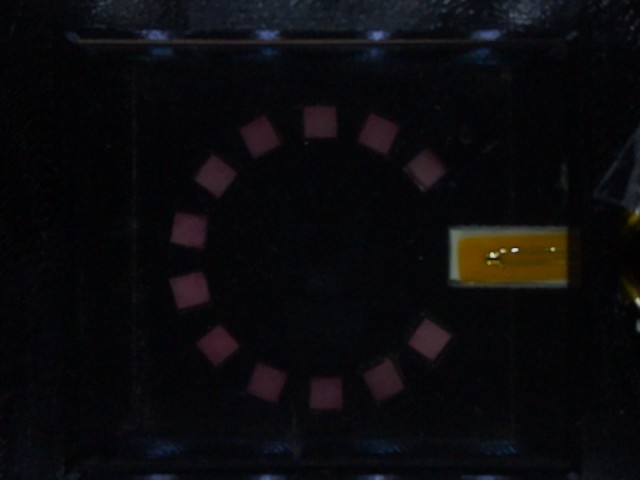

### Gain8_Exp1000_Frame10.jpg

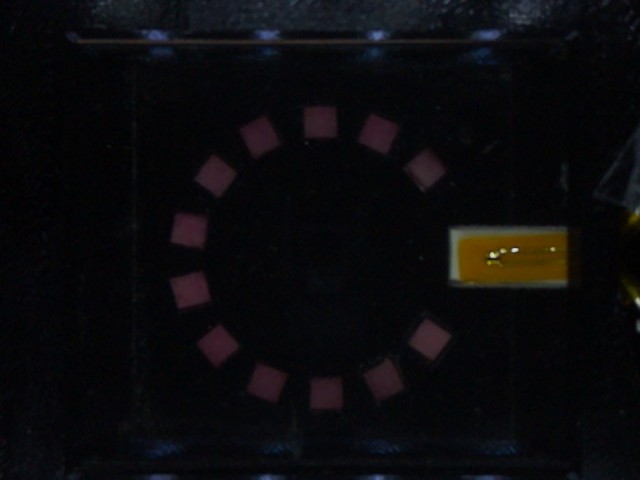

### Gain8_Exp1000_Frame11.jpg

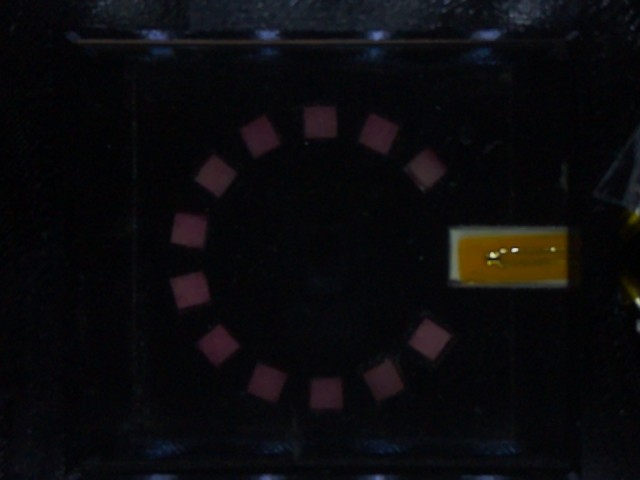

### Gain8_Exp1000_Frame12.jpg

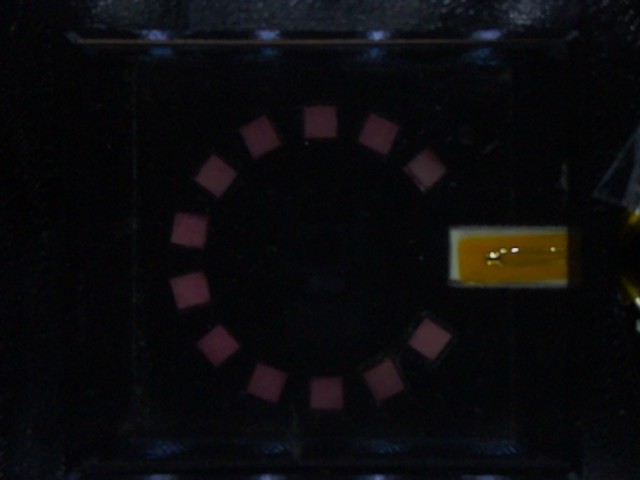

### Gain8_Exp1000_Frame13.jpg

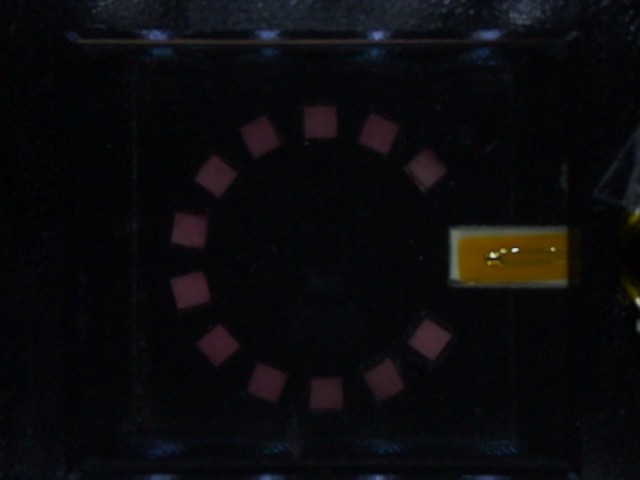

### Gain8_Exp1000_Frame14.jpg

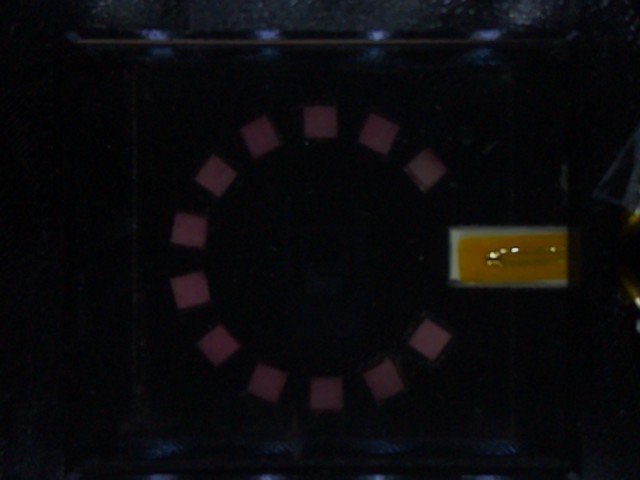

### Gain8_Exp1000_Frame15.jpg

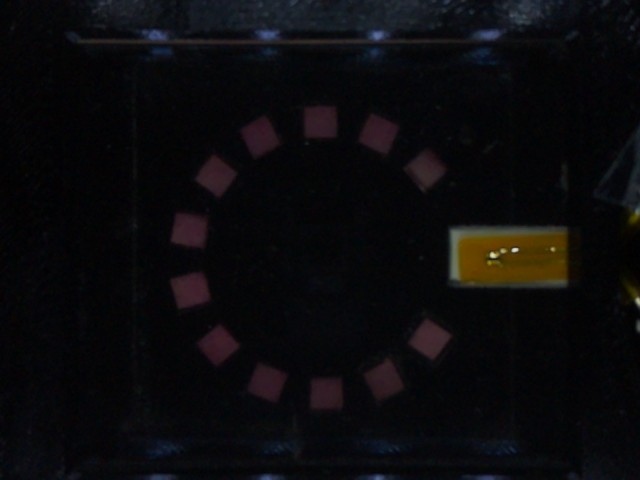

### Gain8_Exp1000_Frame16.jpg

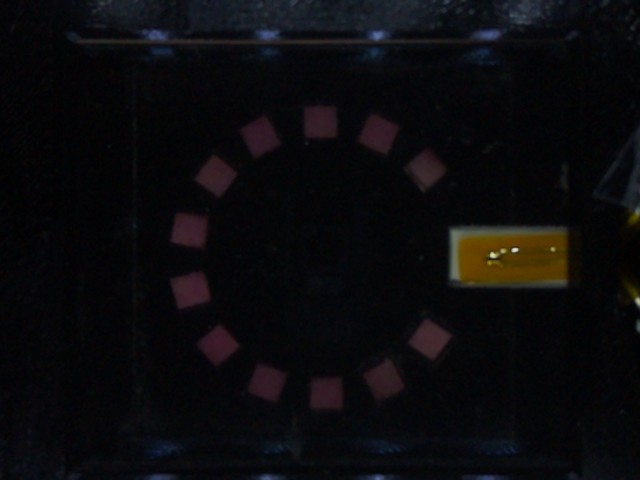

### Gain8_Exp1000_Frame17.jpg

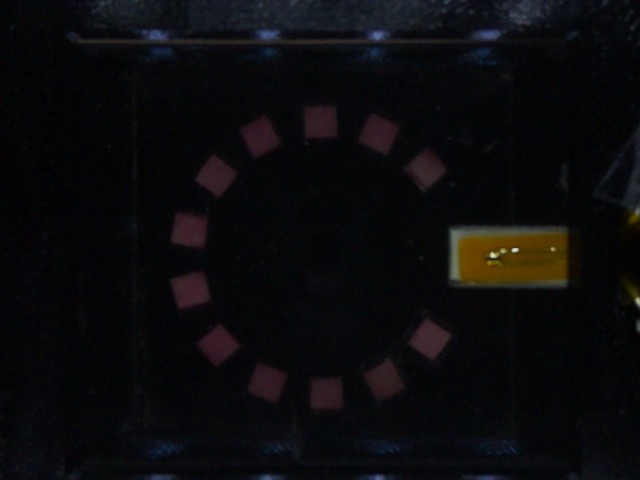

### Gain8_Exp1000_Frame18.jpg

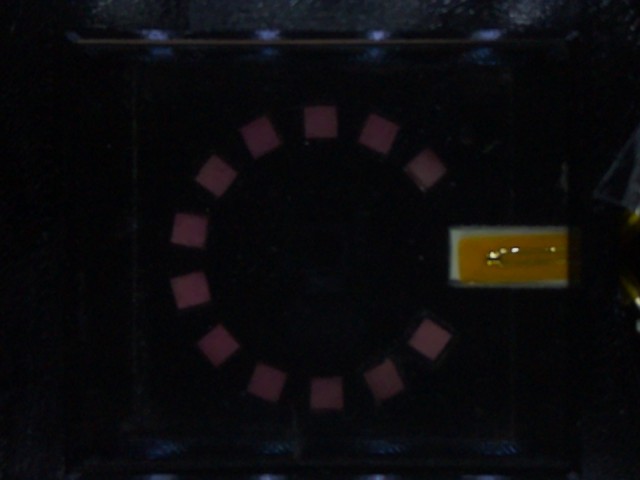

### Gain8_Exp1000_Frame19.jpg

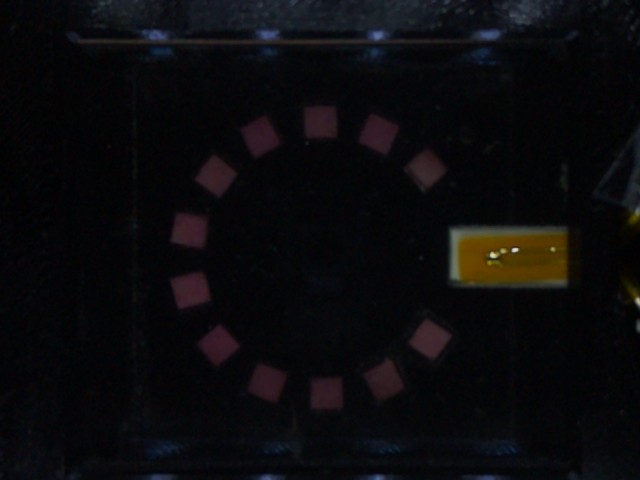

### Gain8_Exp1000_Frame20.jpg

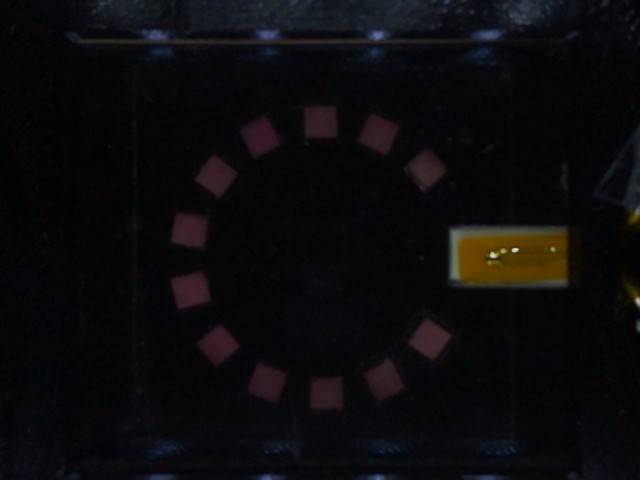

### Gain8_Exp1000_Frame21.jpg

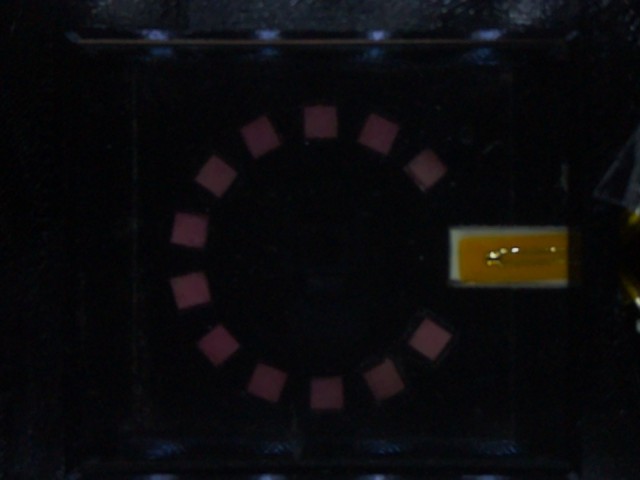

### Gain8_Exp1000_Frame22.jpg

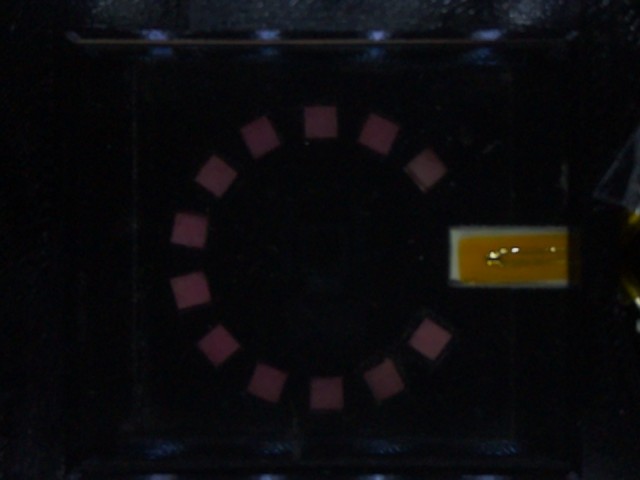

### Gain8_Exp1000_Frame23.jpg

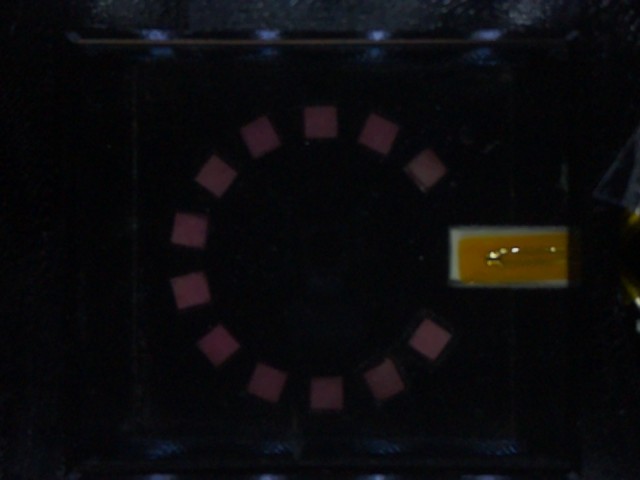

### Gain8_Exp1000_Frame24.jpg

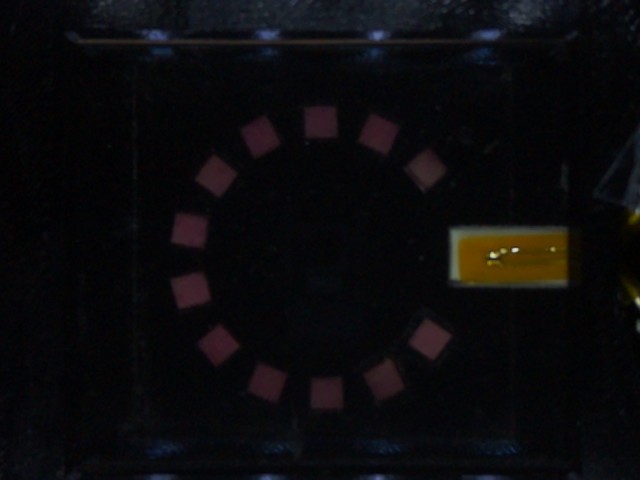

### Gain8_Exp1000_Frame25.jpg

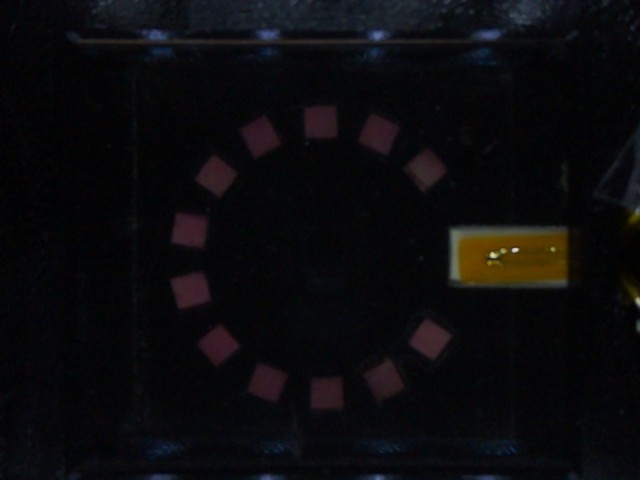

### Gain8_Exp1000_Frame26.jpg

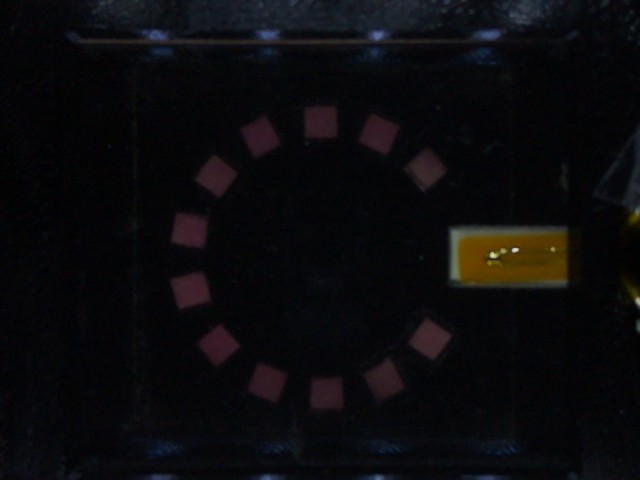

### Gain8_Exp1000_Frame27.jpg

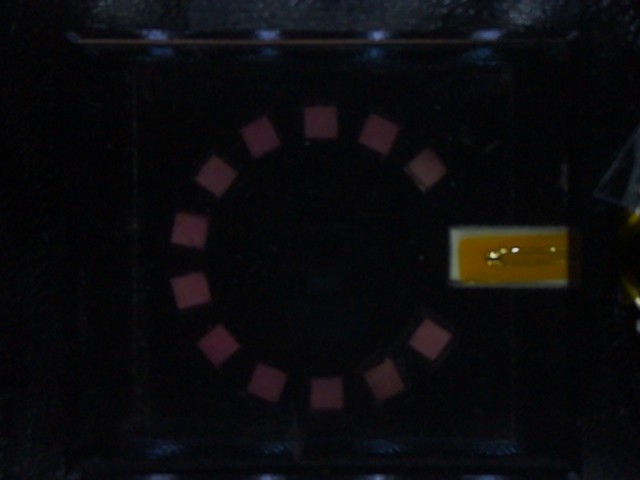

### Gain8_Exp1000_Frame28.jpg

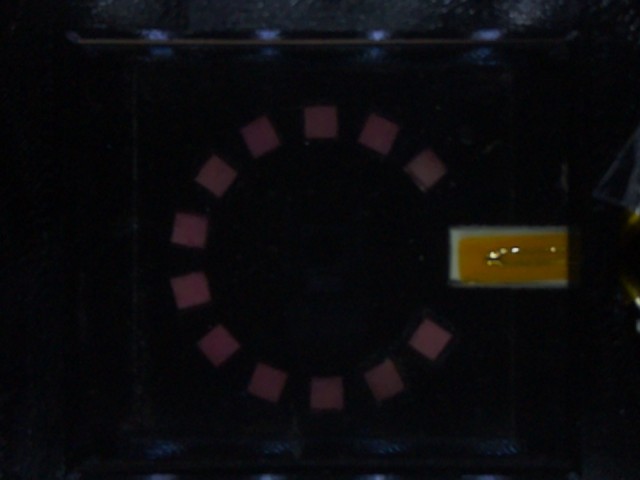

### Gain8_Exp1000_Frame29.jpg

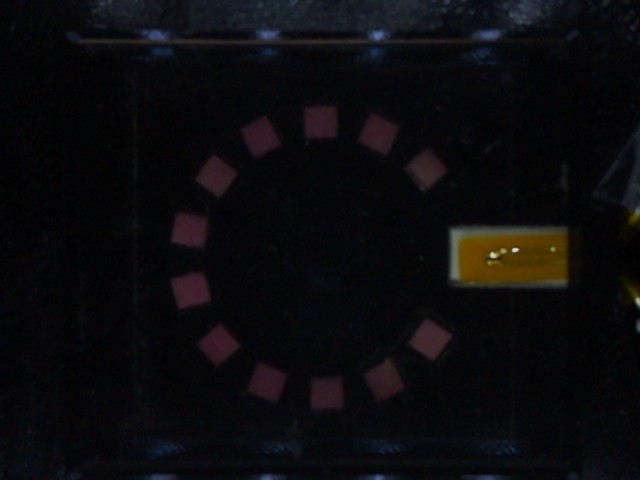

### Gain8_Exp1000_Frame30.jpg

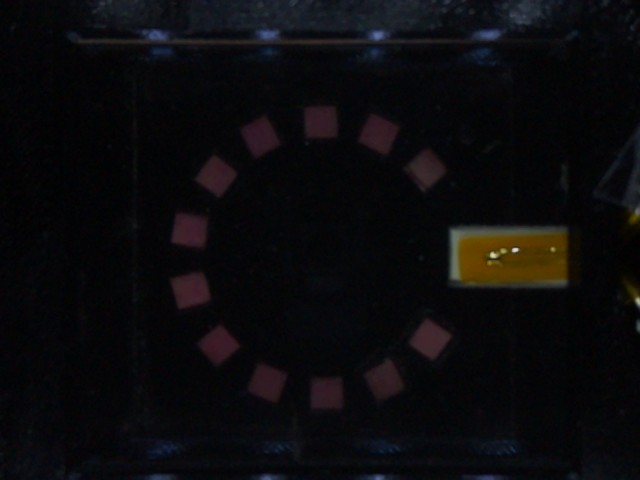

### Gain8_Exp1000_Frame31.jpg

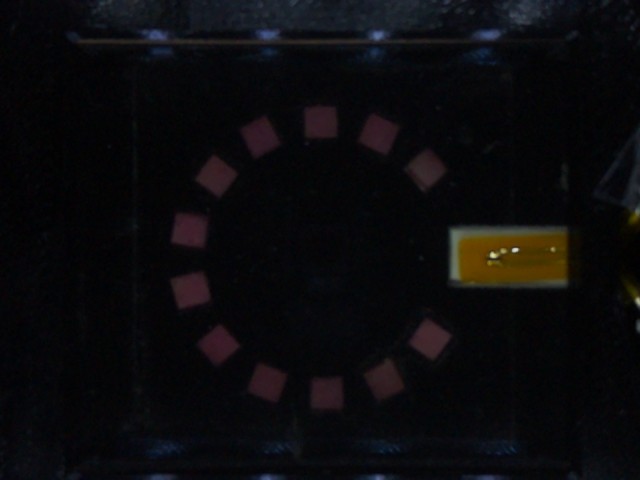

### Gain8_Exp1000_Frame32.jpg

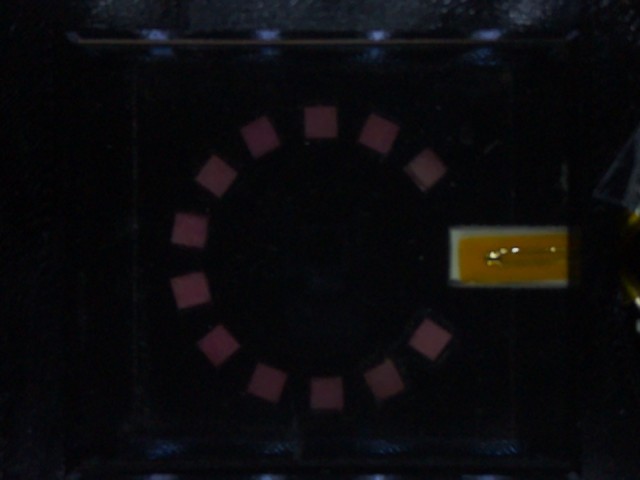

### Gain8_Exp1000_Frame33.jpg

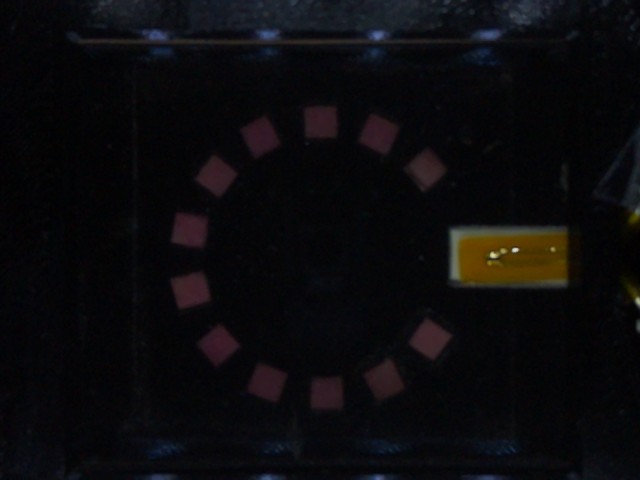

### Gain8_Exp1000_Frame34.jpg

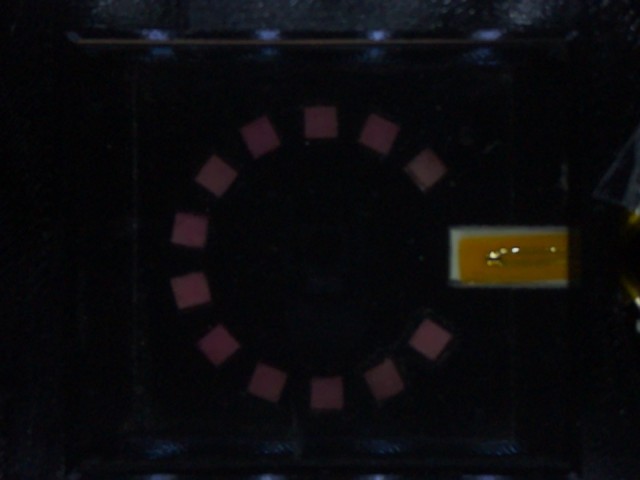

### Gain8_Exp1000_Frame35.jpg

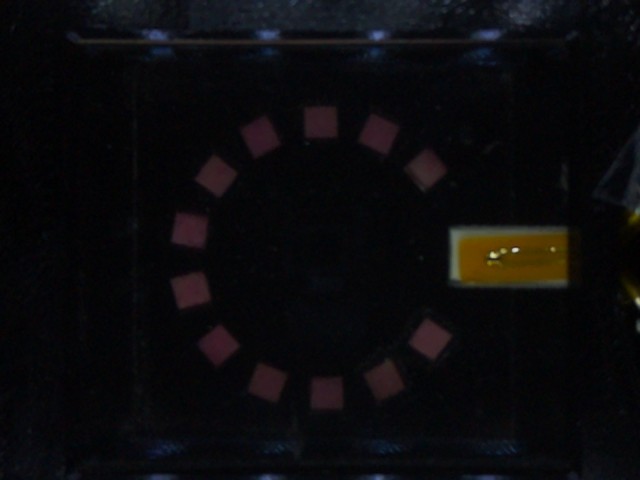

### Gain8_Exp1000_Frame36.jpg

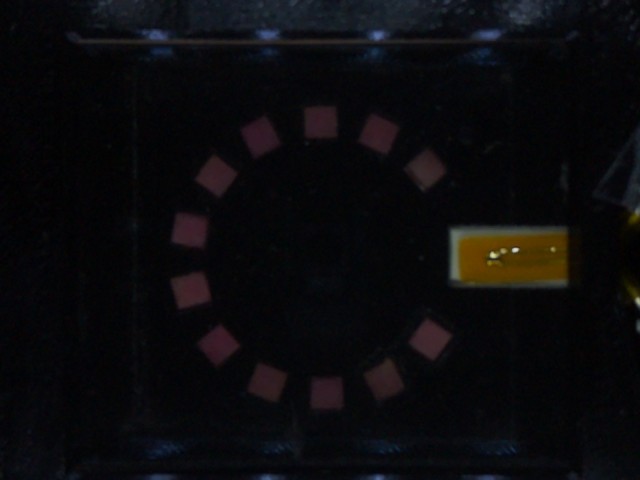

### Gain8_Exp1000_Frame37.jpg

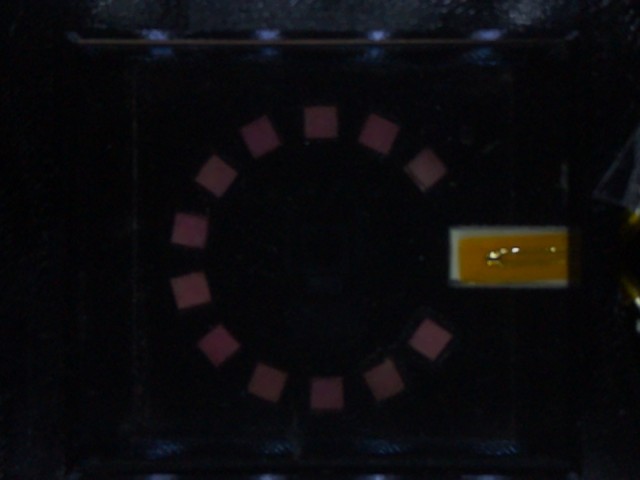

### Gain8_Exp1000_Frame38.jpg

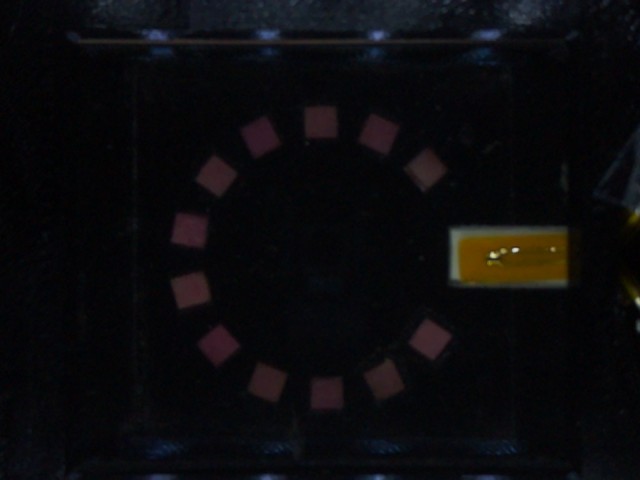

### Gain8_Exp1000_Frame39.jpg

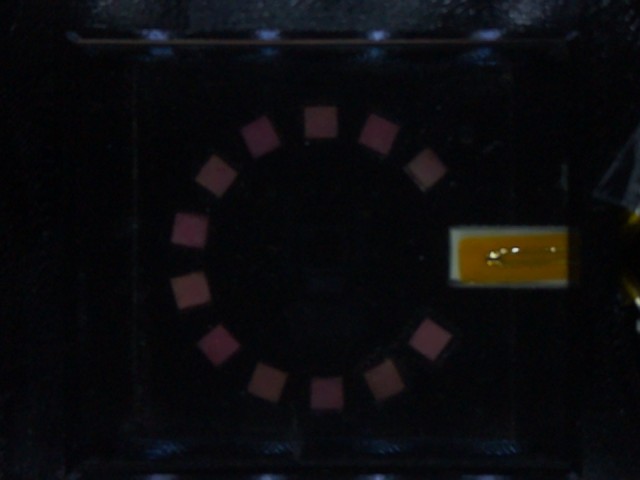

### Gain8_Exp1000_Frame40.jpg

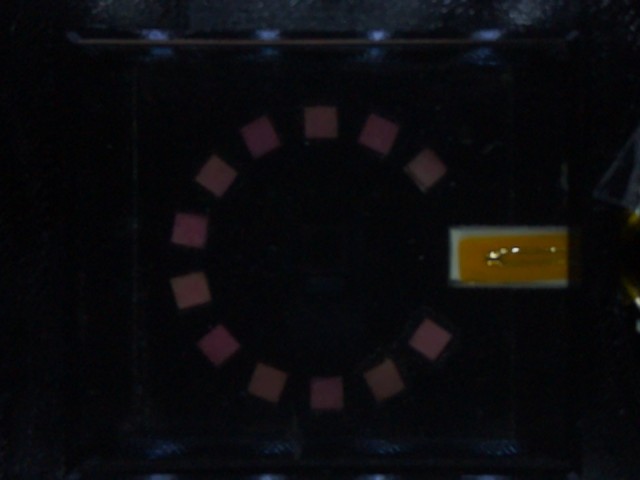

### Gain8_Exp1000_Frame41.jpg

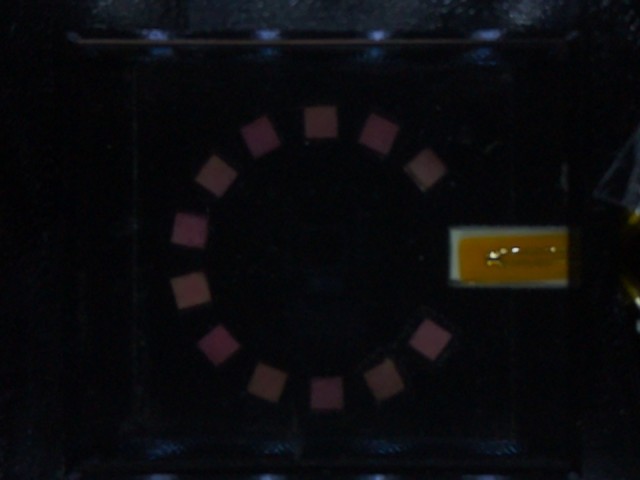

### Gain8_Exp1000_Frame42.jpg

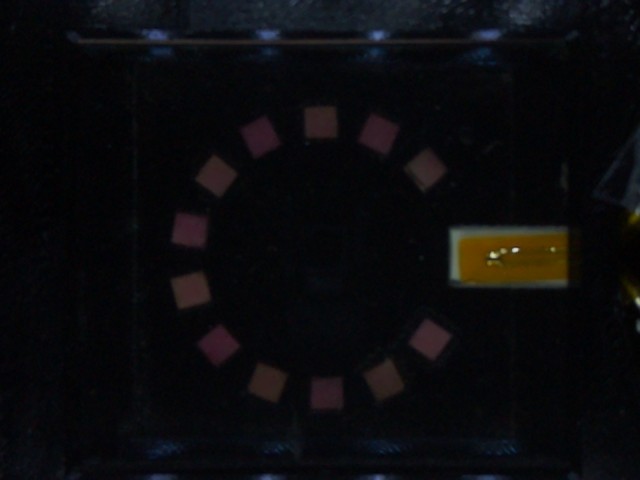

### Gain8_Exp1000_Frame43.jpg

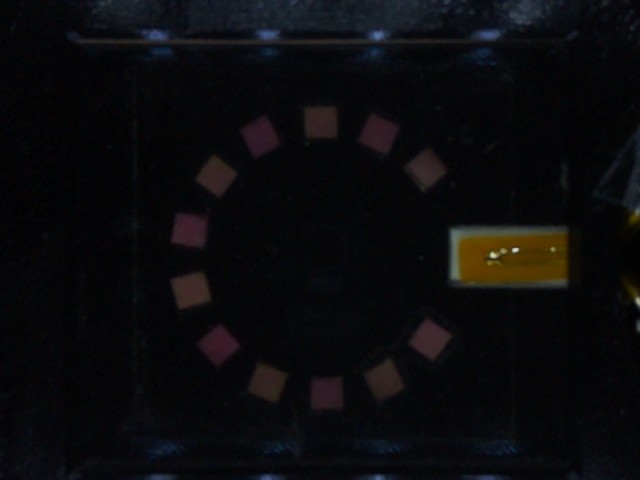

### Gain8_Exp1000_Frame44.jpg

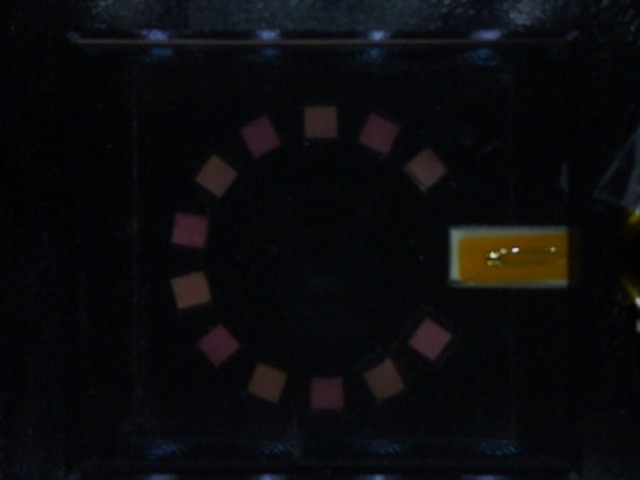

### Gain8_Exp1000_Frame45.jpg

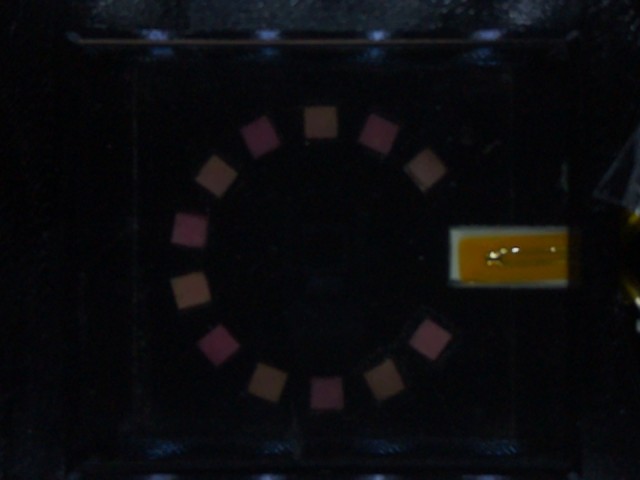

### Gain8_Exp1000_Frame46.jpg

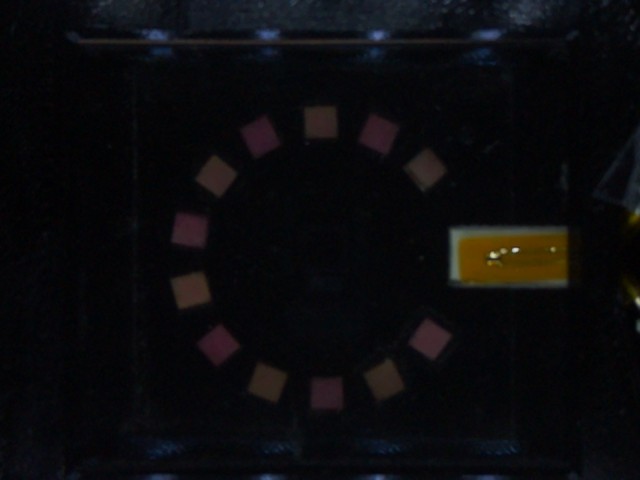

### Gain8_Exp1000_Frame47.jpg

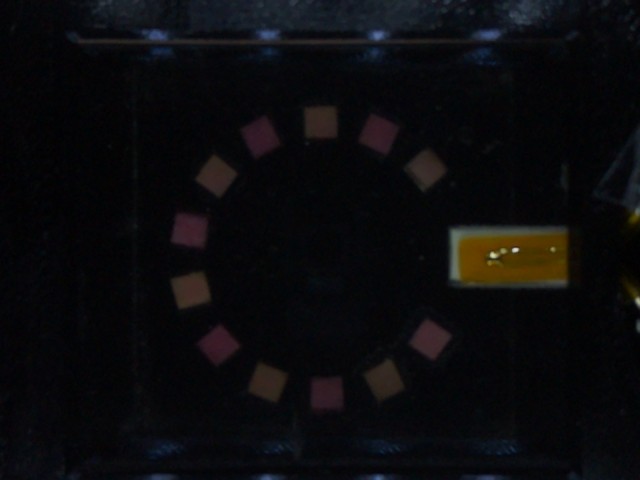

### Gain8_Exp1000_Frame48.jpg

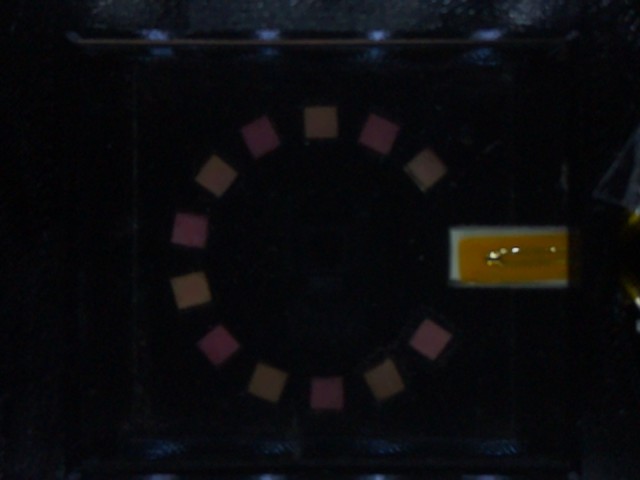

### Gain8_Exp1000_Frame49.jpg

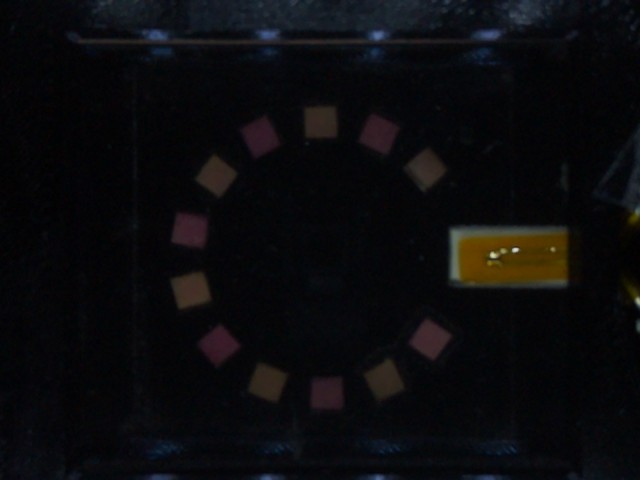

### Gain8_Exp1000_Frame50.jpg

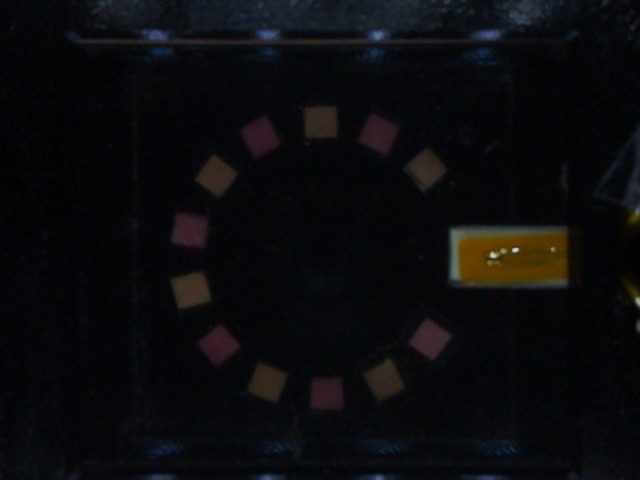

### Gain8_Exp1000_Frame51.jpg

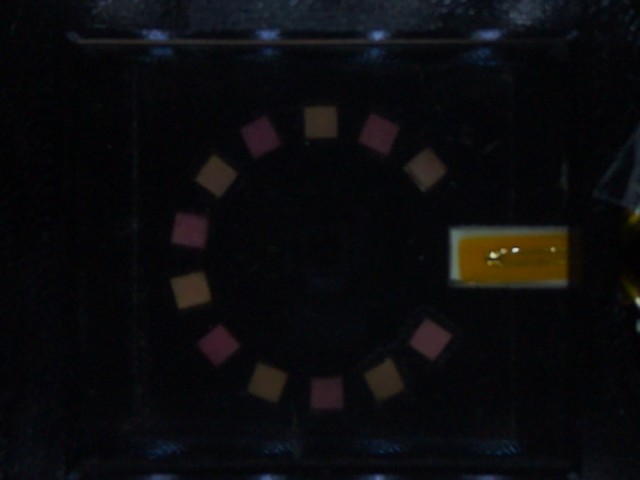

### Gain8_Exp1000_Frame52.jpg

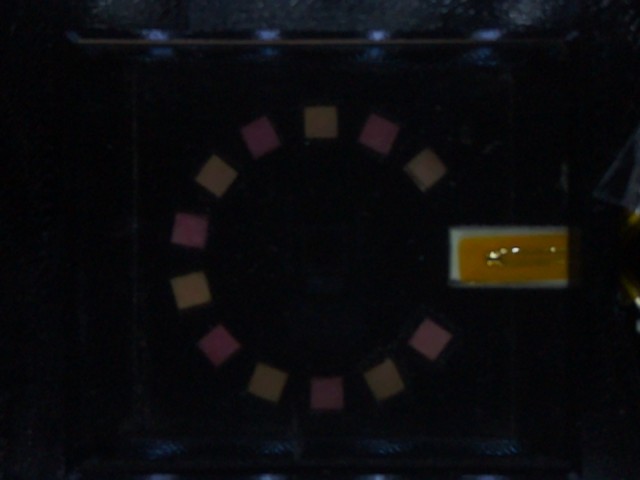

### Gain8_Exp1000_Frame53.jpg

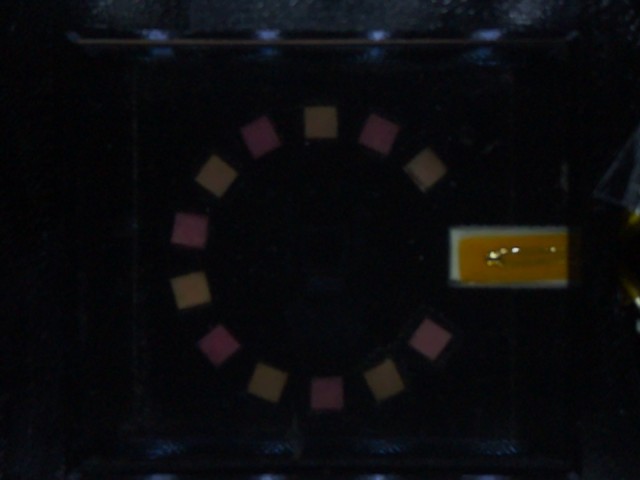

### Gain8_Exp1000_Frame54.jpg

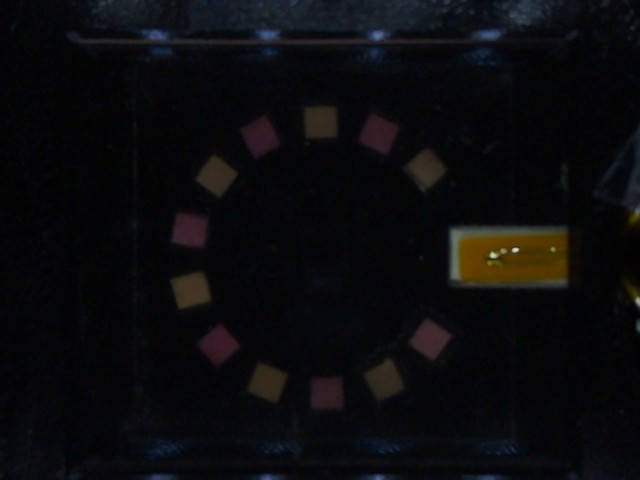

### Gain8_Exp1000_Frame55.jpg

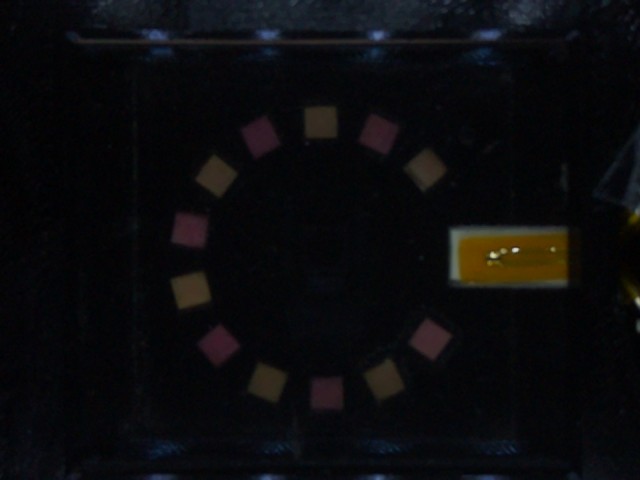

### Gain8_Exp1000_Frame56.jpg

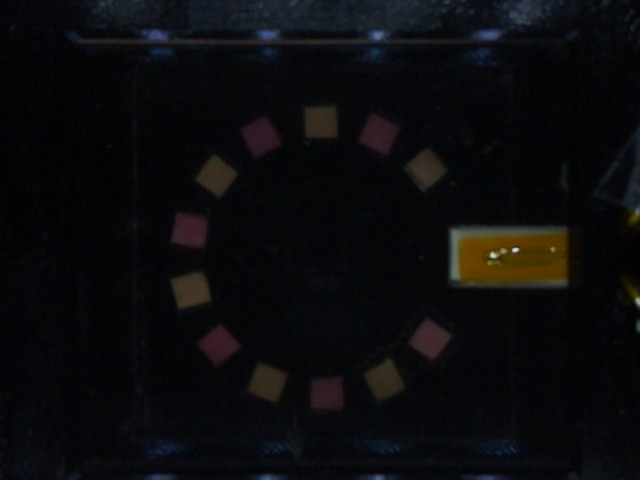

### Gain8_Exp1000_Frame57.jpg

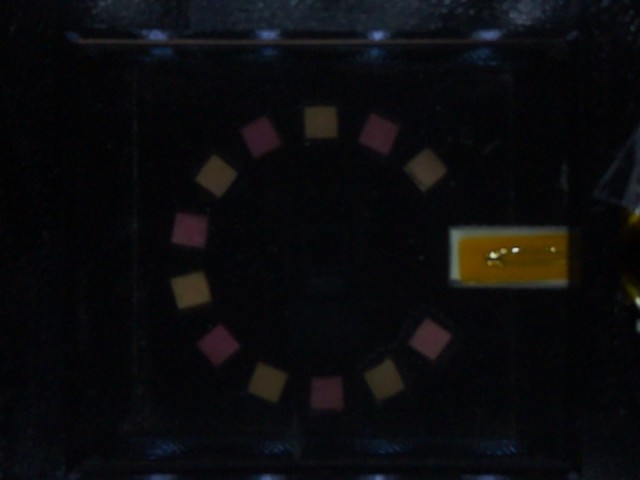

### Gain8_Exp1000_Frame58.jpg

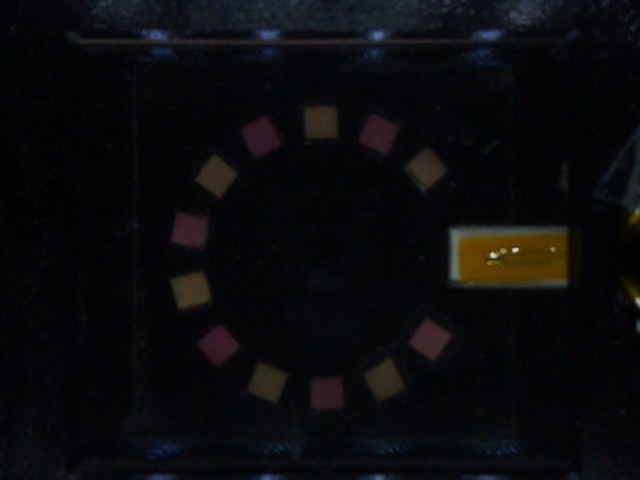

### Gain8_Exp1000_Frame59.jpg

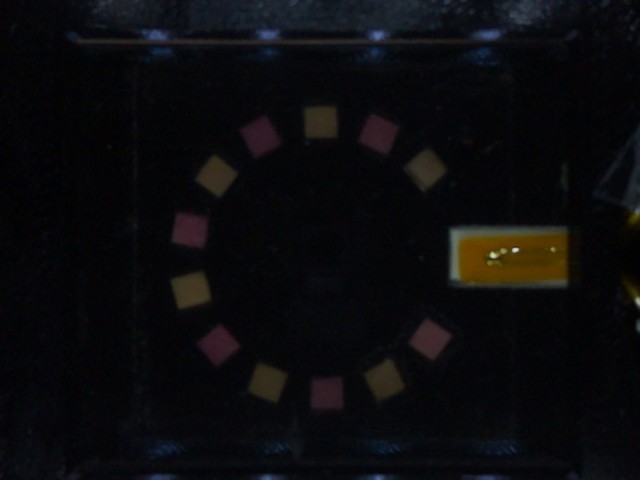

### Gain8_Exp1000_Frame60.jpg

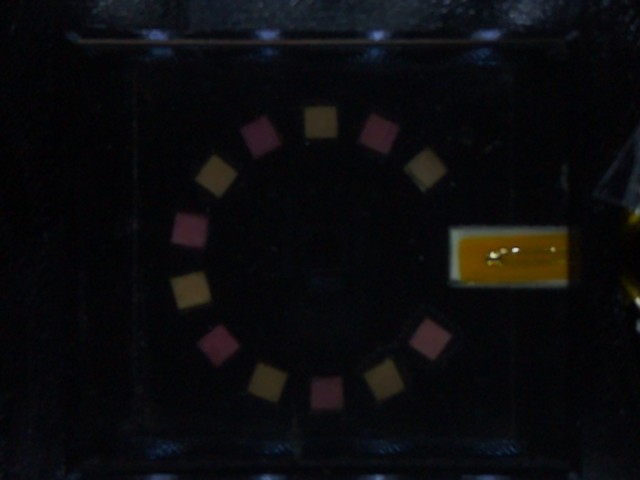

### Gain8_Exp1000_Frame61.jpg

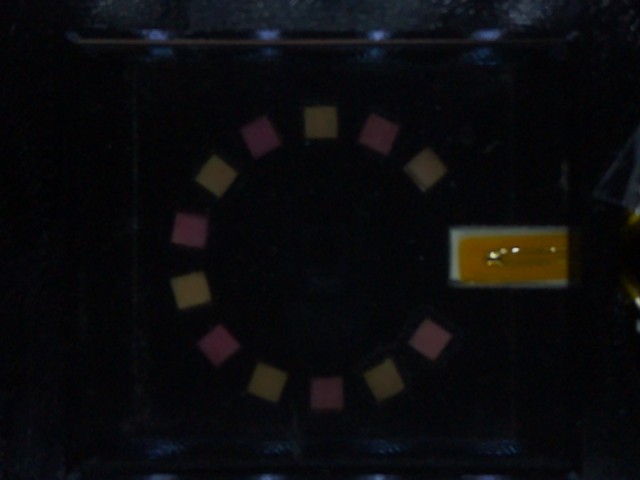

### Gain8_Exp1000_Frame62.jpg

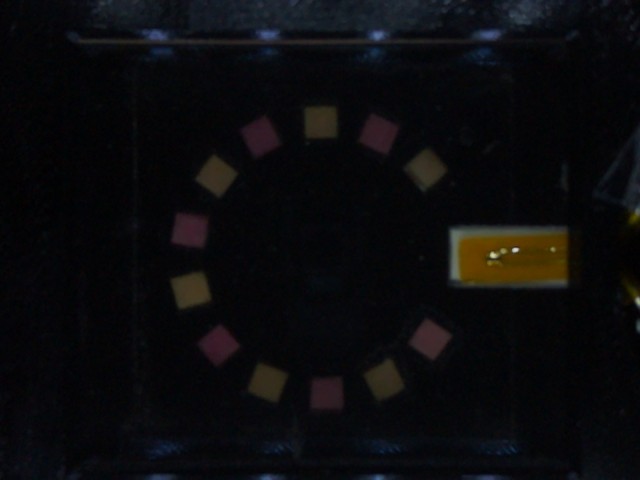

### Gain8_Exp1000_Frame63.jpg

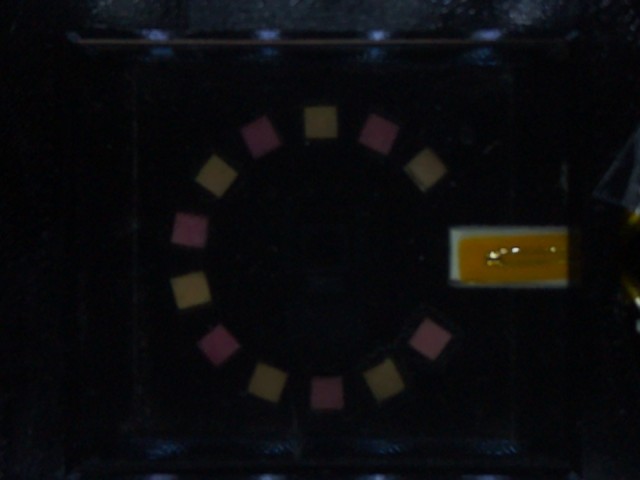

### Gain8_Exp1000_Frame64.jpg

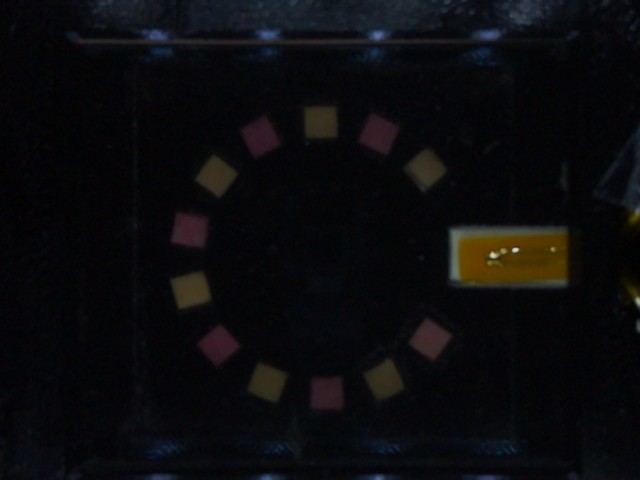

### Gain8_Exp1000_Frame65.jpg

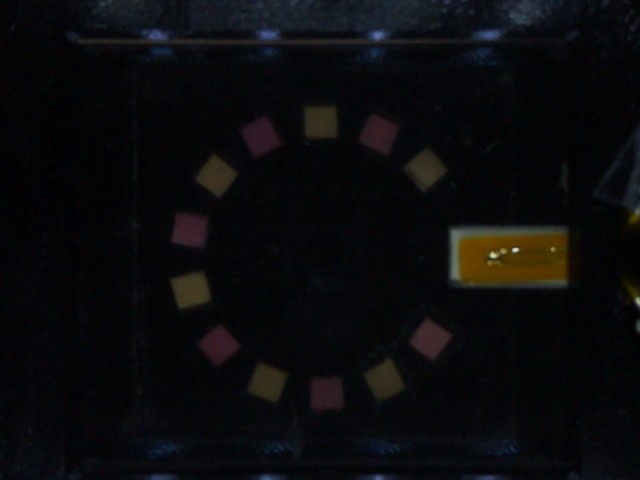

### Gain8_Exp1000_Frame66.jpg

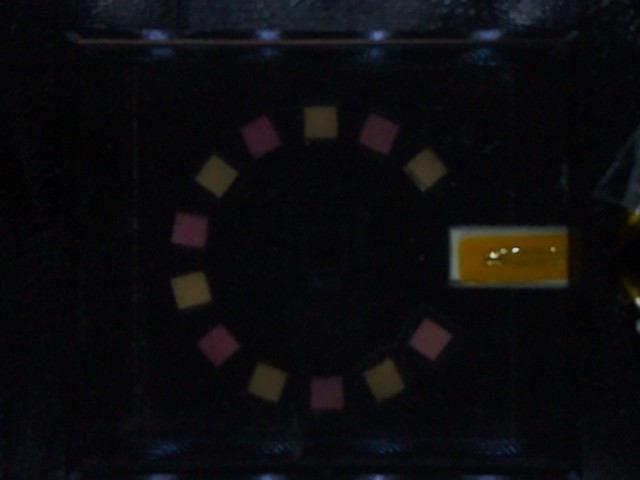

### Gain8_Exp1000_Frame67.jpg

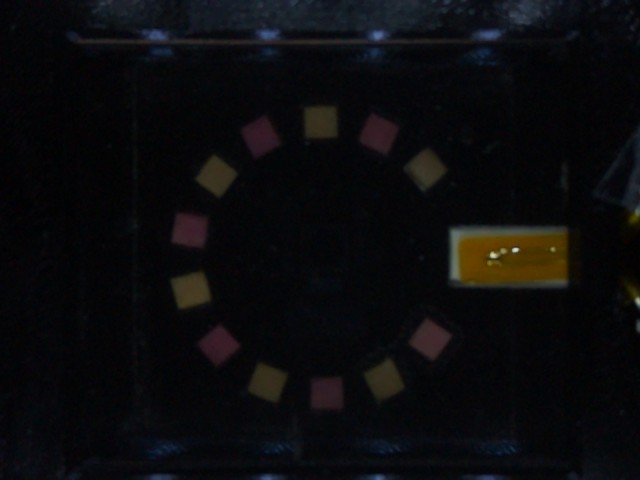

### Gain8_Exp1000_Frame68.jpg

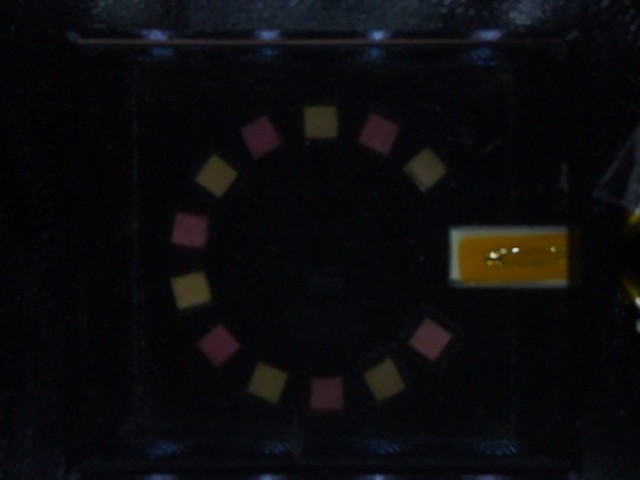

### Gain8_Exp1000_Frame69.jpg

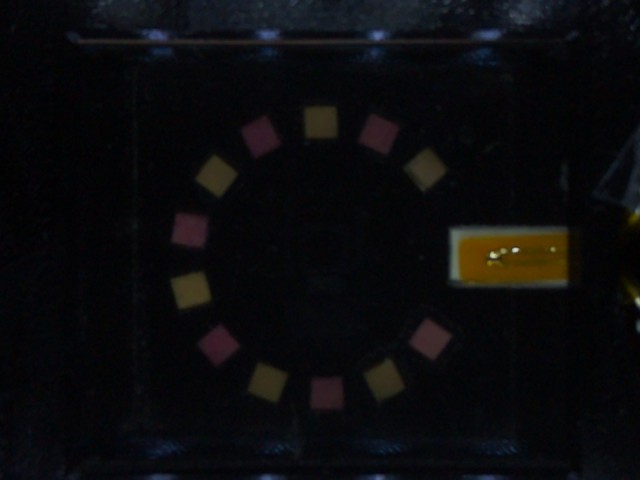

### Gain8_Exp1000_Frame70.jpg

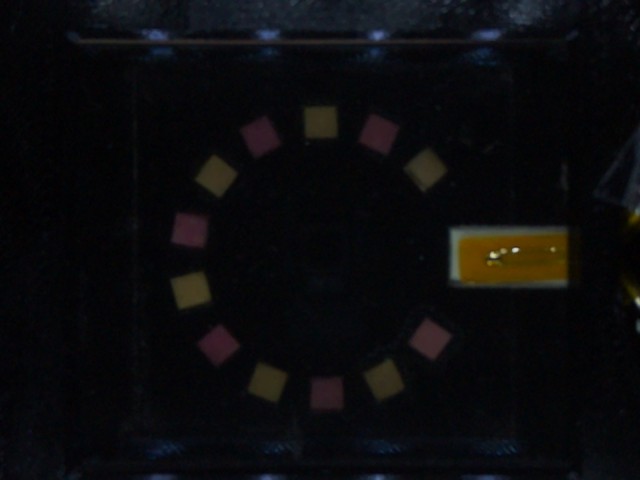

### Gain8_Exp1000_Frame71.jpg

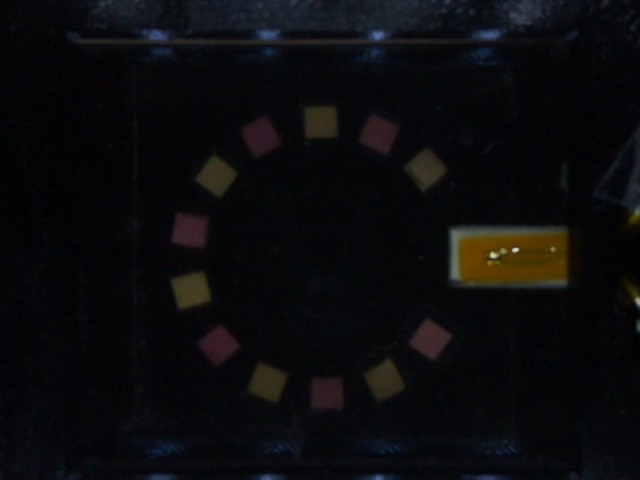

### Gain8_Exp1000_Frame72.jpg

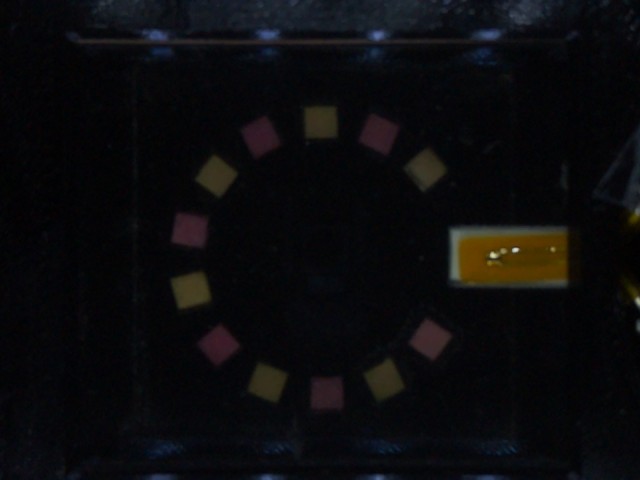

### Gain8_Exp1000_Frame73.jpg

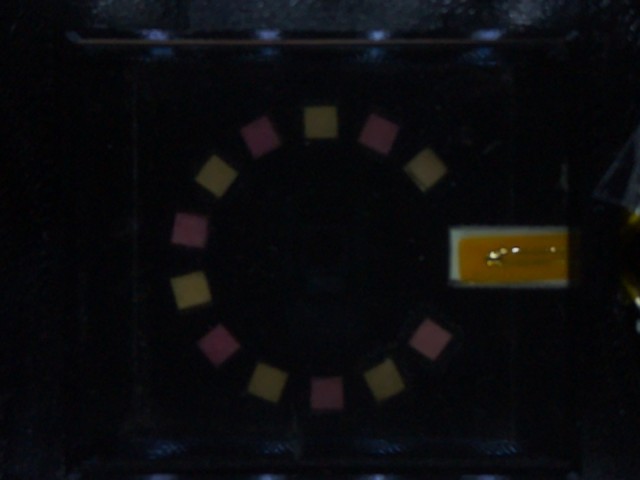

### Gain8_Exp1000_Frame74.jpg

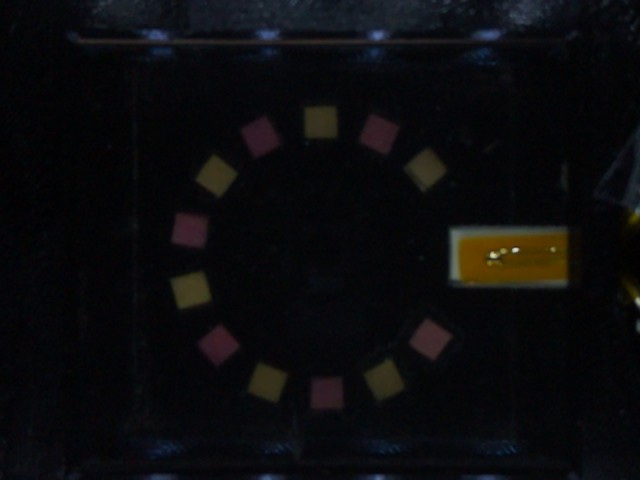

### Gain8_Exp1000_Frame75.jpg

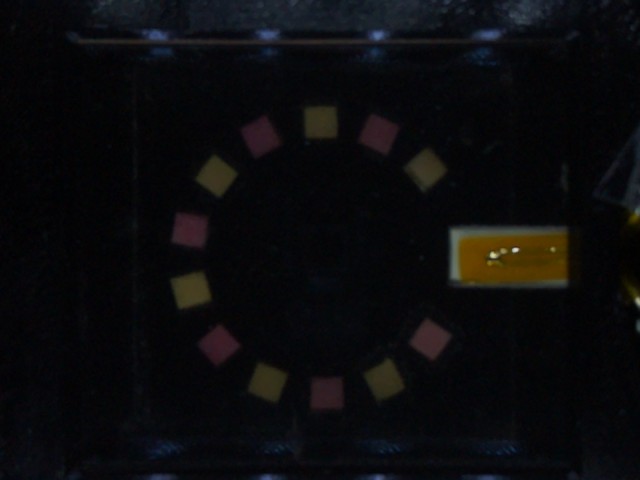

### Gain8_Exp1000_Frame76.jpg

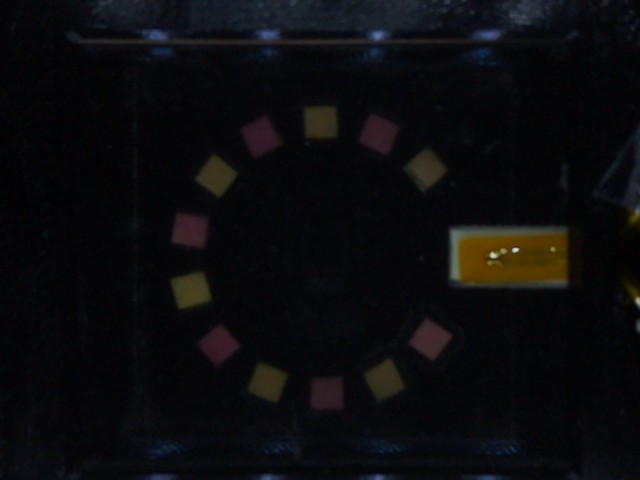

### Gain8_Exp1000_Frame77.jpg

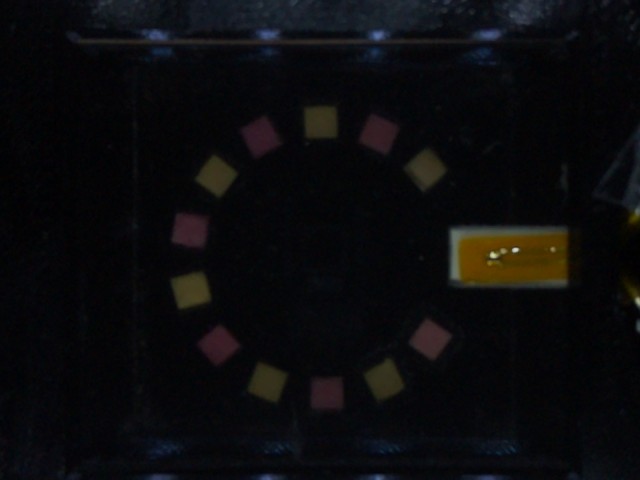

### Gain8_Exp1000_Frame78.jpg

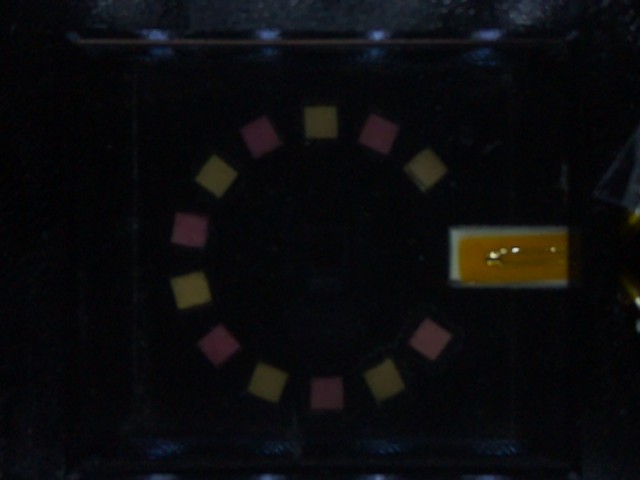

### Gain8_Exp1000_Frame79.jpg

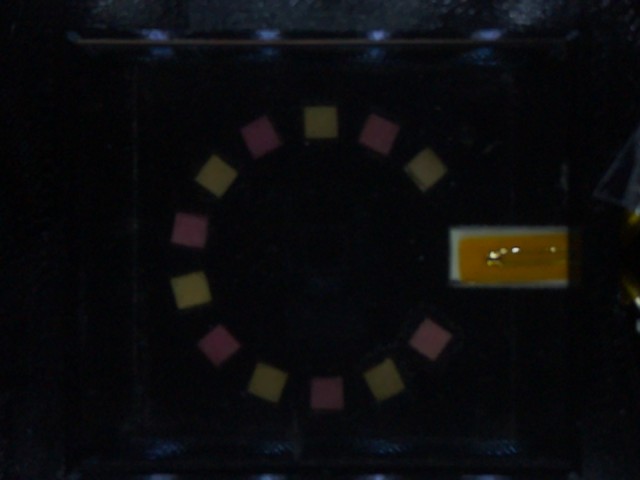

### Gain8_Exp1000_Frame80.jpg

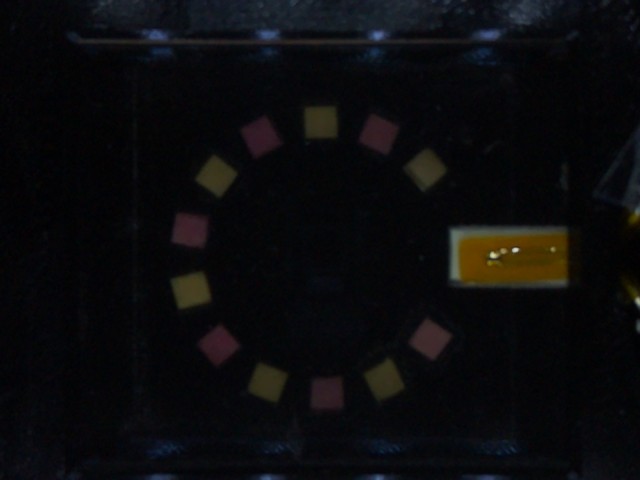

### Gain8_Exp1000_Frame100.jpg

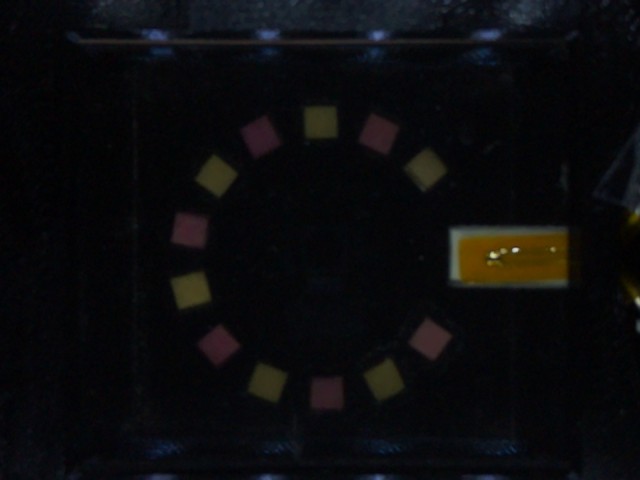

### Gain8_Exp1000_Frame101.jpg

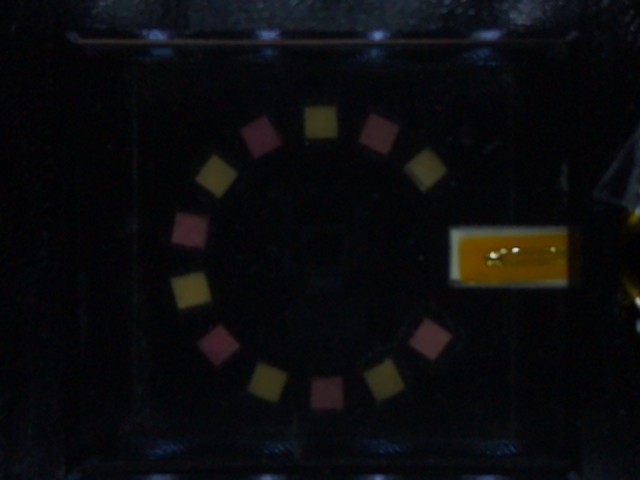

### Gain8_Exp1000_Frame102.jpg

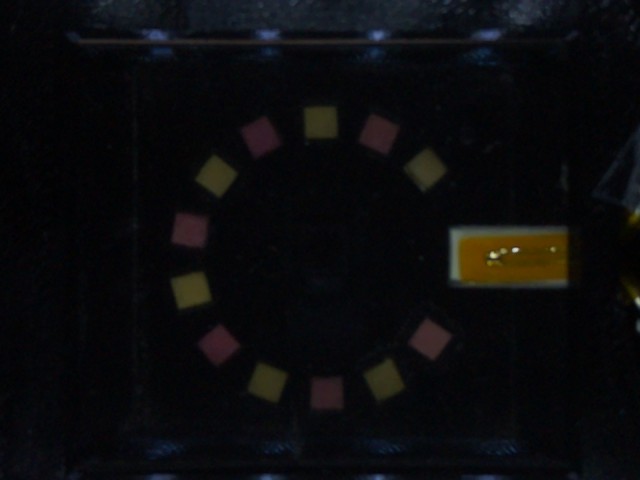

### Gain8_Exp1000_Frame103.jpg

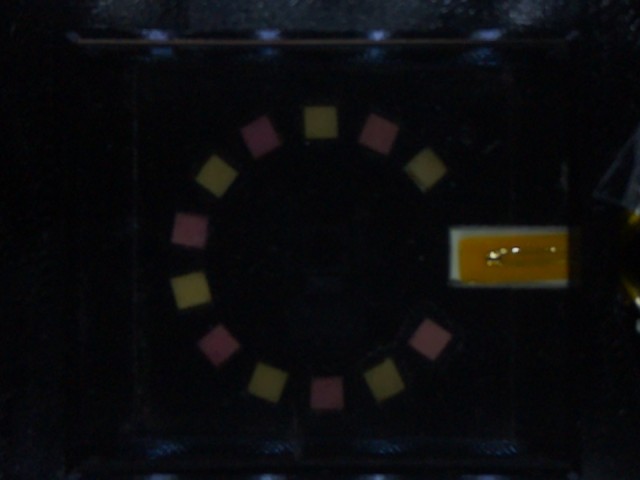

### Gain8_Exp1000_Frame104.jpg

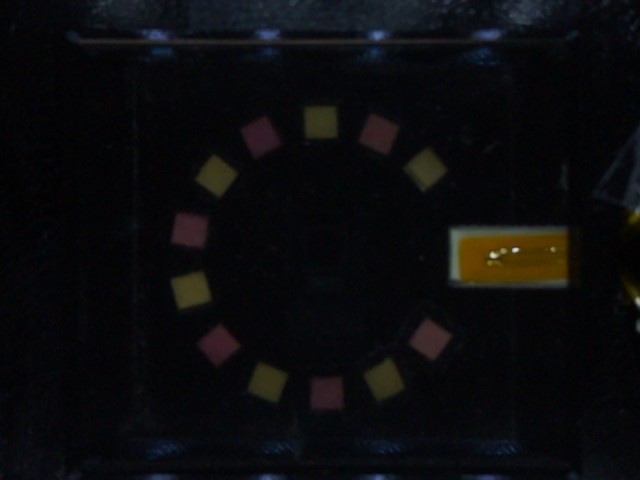

### Gain8_Exp1000_Frame105.jpg

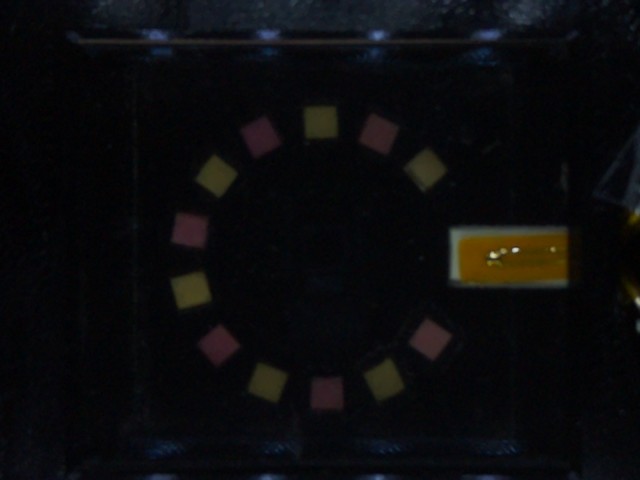

### Gain8_Exp1000_Frame106.jpg

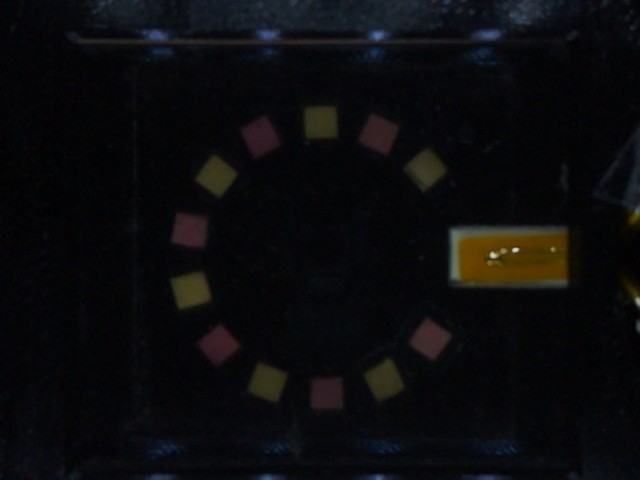

### Gain8_Exp1000_Frame107.jpg

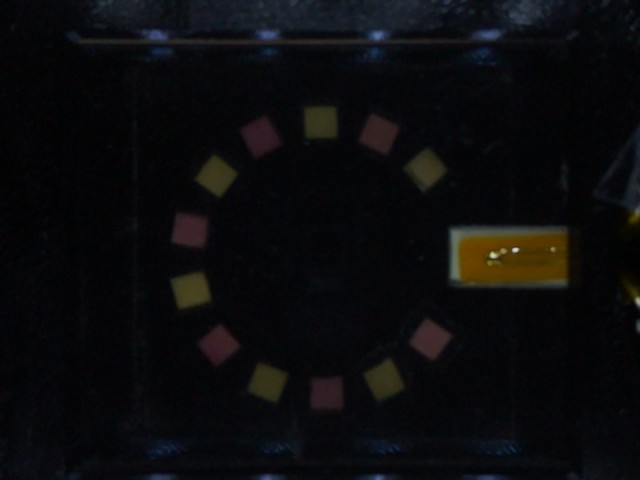

### Gain8_Exp1000_Frame108.jpg

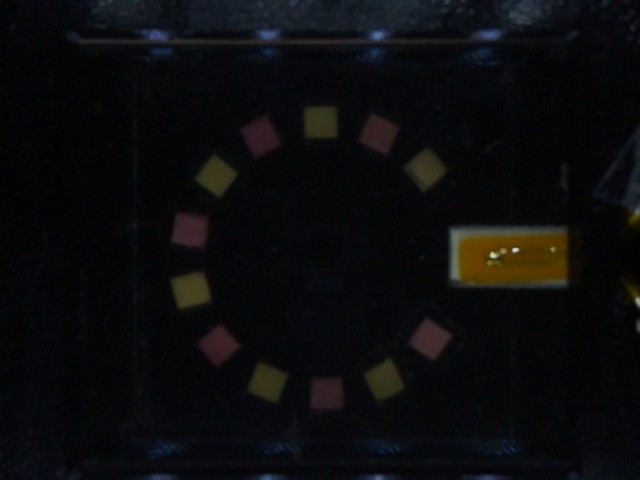

### Gain8_Exp1000_Frame109.jpg

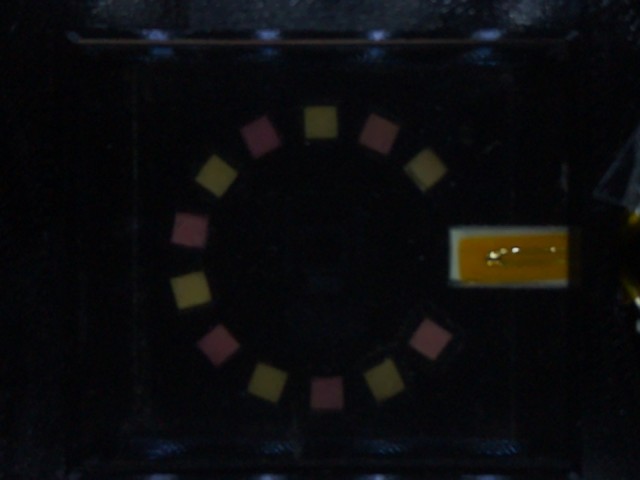

### Gain8_Exp1000_Frame110.jpg

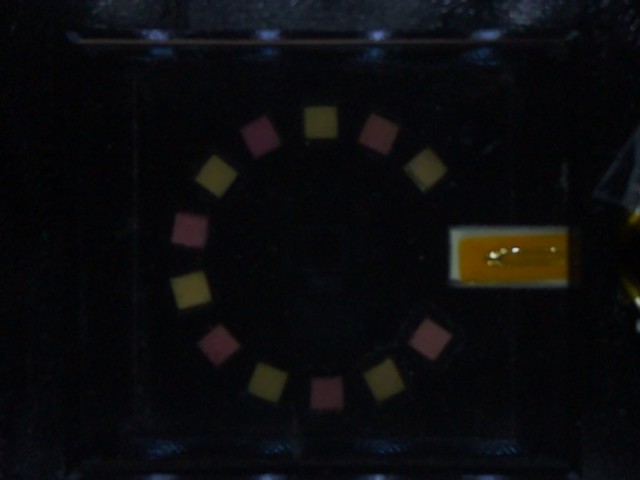

### Gain8_Exp1000_Frame111.jpg

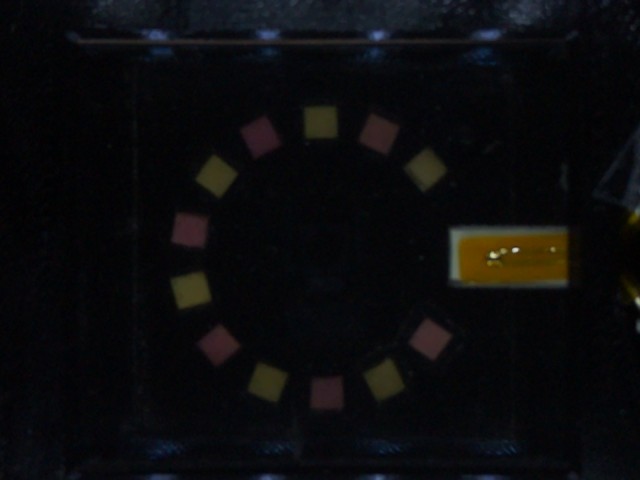

### Gain8_Exp1000_Frame112.jpg

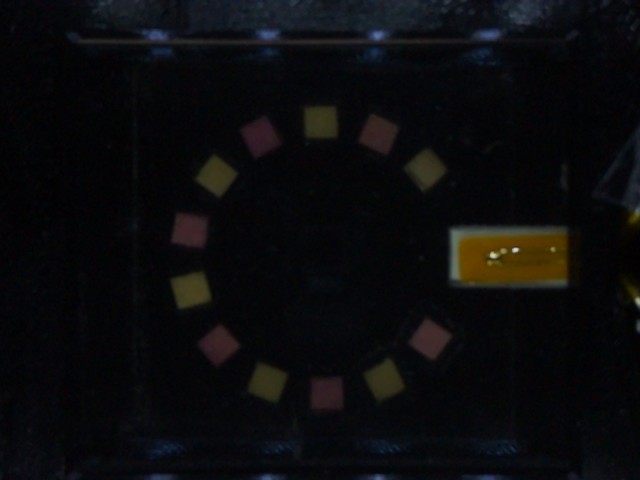

### Gain8_Exp1000_Frame113.jpg

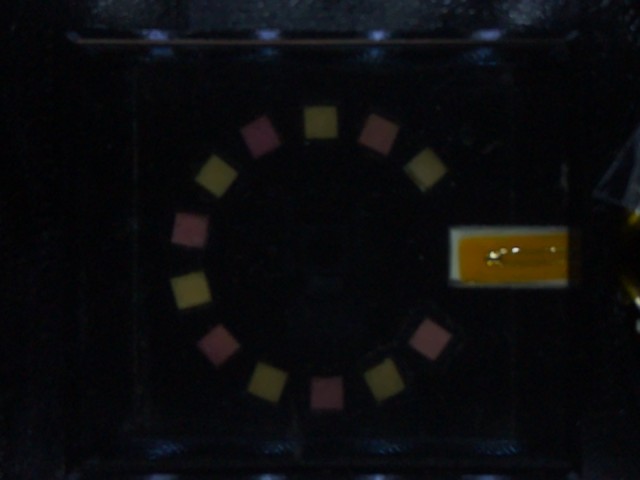

### Gain8_Exp1000_Frame114.jpg

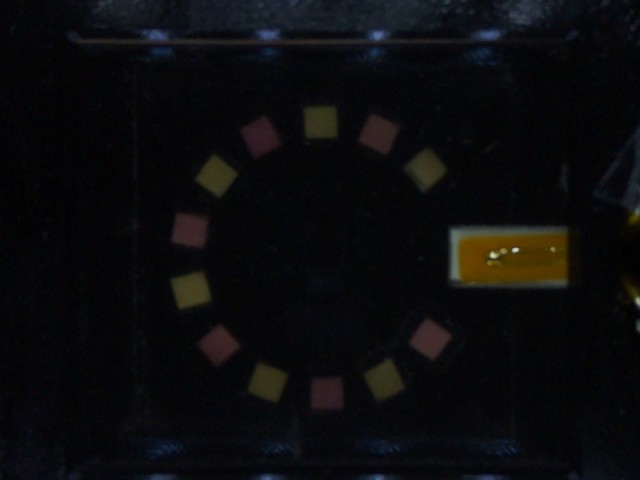

### Gain8_Exp1000_Frame115.jpg

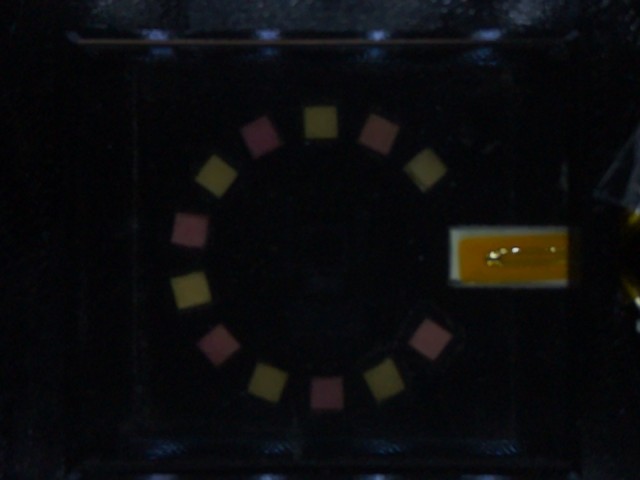

### Gain8_Exp1000_Frame116.jpg

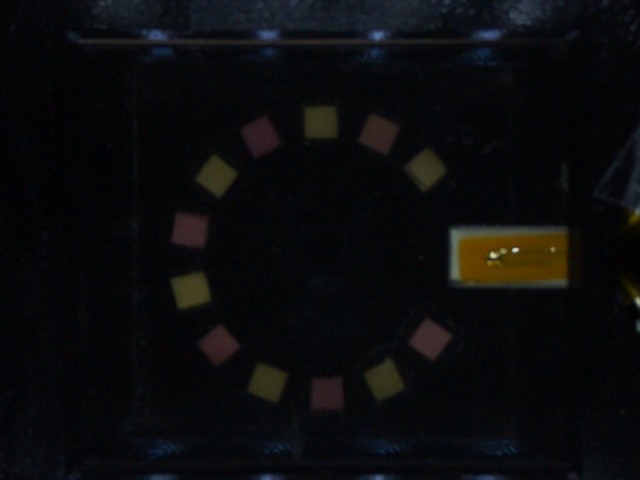

### Gain8_Exp1000_Frame117.jpg

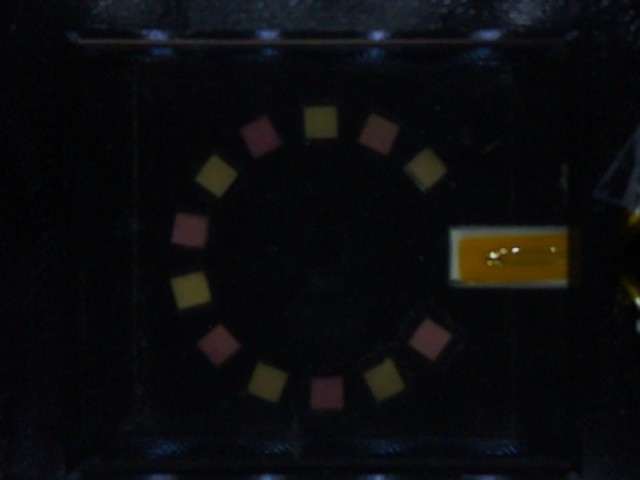

### Gain8_Exp1000_Frame118.jpg

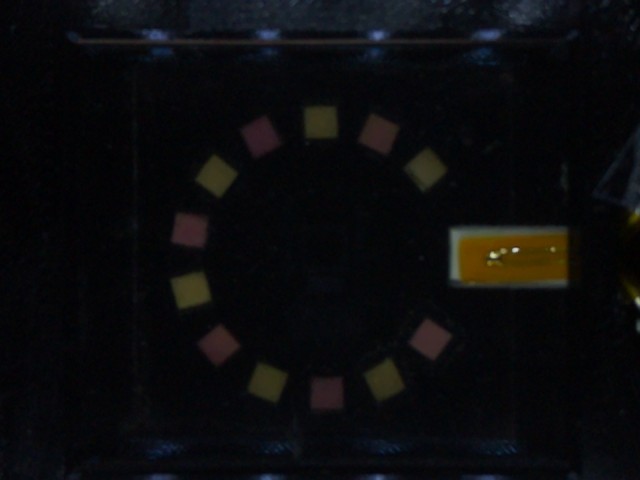

### Gain8_Exp1000_Frame119.jpg

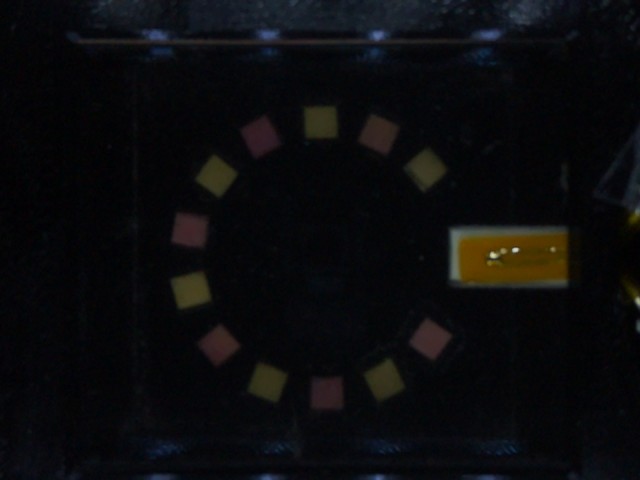

### Gain8_Exp1000_Frame120.jpg

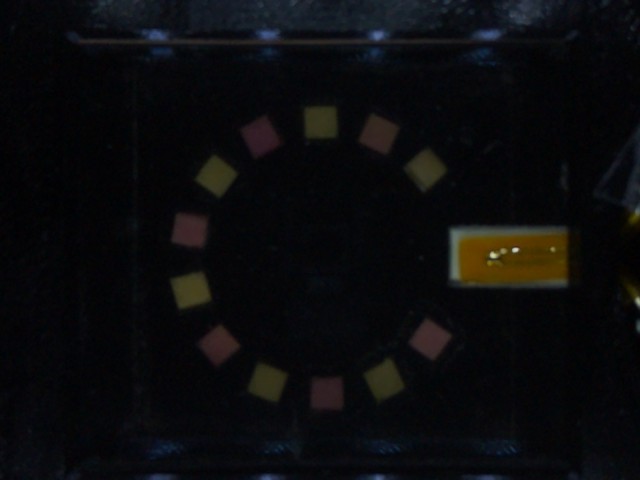
